# A Topological Data Analytic Approach for Discovering Biophysical Signatures in Protein Dynamics

**DOI:** 10.1101/2021.07.28.454240

**Authors:** Wai Shing Tang, Gabriel Monteiro da Silva, Henry Kirveslahti, Erin Skeens, Bibo Feng, Timothy Sudijono, Kevin K. Yang, Sayan Mukherjee, Brenda Rubenstein, Lorin Crawford

## Abstract

Identifying structural differences among proteins can be a non-trivial task. When contrasting ensembles of protein structures obtained from molecular dynamics simulations, biologically-relevant features can be easily overshadowed by spurious fluctuations. Here, we present SINATRA Pro, a computational pipeline designed to robustly identify topological differences between two sets of protein structures. Algorithmically, SINATRA Pro works by first taking in the 3D atomic coordinates for each protein snapshot and summarizing them according to their underlying topology. Statistically significant topological features are then projected back onto a user-selected representative protein structure, thus facilitating the visual identification of biophysical signatures of different protein ensembles. We assess the ability of SINATRA Pro to detect minute conformational changes in five independent protein systems of varying complexities. In all test cases, SINATRA Pro identifies known structural features that have been validated by previous experimental and computational studies, as well as novel features that are also likely to be biologically-relevant according to the literature. These results highlight SINATRA Pro as a promising method for facilitating the non-trivial task of pattern recognition in trajectories resulting from molecular dynamics simulations, with substantially increased resolution.

**Author Summary:** Structural features of proteins often serve as signatures of their biological function and molecular binding activity. Elucidating these structural features is essential for a full understanding of underlying biophysical mechanisms. While there are existing methods aimed at identifying structural differences between protein variants, such methods do not have the capability to jointly infer both geometric and dynamic changes, simultaneously. In this paper, we propose SINATRA Pro, a computational framework for extracting key structural features between two sets of proteins. SINATRA Pro robustly outperforms standard techniques in pinpointing the physical locations of both static and dynamic signatures across various types of protein ensembles, and it does so with improved resolution.

## Introduction

Identifying structural features associated with macromolecular dynamics is crucial to our understanding of the underlying physical behavior of proteins and their broader impact on biology and health. Structural and dynamical properties of proteins often serve as signatures of their functions and activities [1]. Subtle topological changes in protein conformation can lead to dramatic changes in biological function [2,3], thus highlighting the importance of being able to accurately characterize protein conformational dynamics.

Conventionally, the structural dynamics of proteins have been modeled using molecular dynamics (MD) simulations, which work by sampling structural ensembles from conformational landscapes. In infinite timescales, such structural ensembles are expected to represent all physical states such that their ensemble-averaged observables converge to true physical values and are thus physically meaningful. While MD simulations have provided key insights into the atomistic motions that underpin many protein functions [4], biologically-relevant structural changes can be overshadowed by spurious statistical noise caused by the thermal fluctuations that naturally arise during the course of these simulations [5]. In practice, this can often make important structural features difficult to identify and robustly interpret from MD trajectories. Data from MD simulations are often analyzed in a strictly goal-dependent manner by using computational methods that quantify and assess specific protein characteristics. For example, geometric changes that arise as a result of ligand binding, point mutations, or post-translational modifications are usually inferred by analyzing the root mean square fluctuations (RMSF) of atomic positions or the per-domain radius of gyration with respect to a reference structure [6]. Unfortunately, these standard approaches are less powerful when the relevant changes in protein structure are overshadowed by fluctuations irrelevant to the biological process of interest.

Recently, more sophisticated methods have aimed to overcome these challenges by taking advantage of correspondences between the atomic positions on any two given proteins. For example, per-residue distance functions or contact maps can be calculated on each frame of a trajectory for clustering [7] or principal component analyses (PCA) [8,9], which project complex conformations onto a lower-dimensional space for ease of comparison. However, the downside to these methods is that they require diffeomorphisms between structures (i.e., the map from protein *A* to protein *B* must be differentiable). There are many scenarios in protein dynamics where no such transformation is guaranteed because atomic features can be gained or lost during the evolution of the system [10]. Indeed, there are 3D shape algorithms that construct more general “functional” correspondences and can be applied even across shapes having different topology [11, 12]; however, previous work has shown that the performance of these algorithms drops significantly when the assumed functional mapping input is even slightly misspecified [13].

In this work, we introduce SINATRA Pro: a topological data analytic pipeline for identifying biologically-relevant structural differences between two protein structural ensembles without the need for explicit contact maps or atomic correspondences. Our algorithm is an extension of a previous framework, SINATRA, which was broadly introduced to perform variable selection on physical features that best describe the variation between two groups of static 3D shapes [13]. Using a tool from integral geometry and differential topology called the Euler characteristic (EC) transform [14–17], SINATRA was shown to have the power to identify known morphological perturbations in controlled simulations and robustly identify anatomical aberrations in mandibular molars associated within four different suborders of primates. SINATRA Pro is an adaptation of the SINATRA framework for protein dynamics. Here, we develop a simplicial complex construction step to specifically model both 3D geometric and topological relationships between atomic positions on protein structures. We also utilize a new set of statistical parameters which we calibrate for complex protein systems.

In this study, we demonstrate SINATRA Pro’s ability to identify key structural and dynamical features in a hierarchy of proteins with increasingly challenging features to statistically resolve. The five proteins studied, TEM *β*-lactamase, the Abelson Kinase (Abl1), the HIV-1 protease, Elongation Factor Thermo Unstable (EF-Tu), and Importin-*β*, undergo structural changes in response to a wide range of well-studied biological phenomena, including mutations, interactions with partners, and small molecule binding. We show that SINATRA Pro outperforms standard analytic techniques including RMSF and PCA in consistently pinpointing physical locations of biologically-relevant conformational changes. Overall, we find that SINATRA Pro holds great promise for extracting topological differences between two sets of protein structures from meaningless statistical noise.

## Results

### Pipeline Overview

The SINATRA Pro pipeline involves five key steps (see Fig. 1). First, the algorithm begins by taking aligned structures from two all-atom protein MD simulation trajectories of different phenotypic states (e.g., wild-type versus mutant) as inputs (Fig. 1(a)). In the second step, SINATRA Pro uses the 3D Cartesian coordinates of each atom of the proteins to create mesh representations of their 3D structures (Fig. 1(b)). Here, atoms within a predetermined physical distance cutoff (e.g., *∼* 6 Ångströms (Å) apart) are connected by “edges” and then triangles enclosed by the connected edges are filled to create “faces.” In the third step, we convert the resulting triangulated meshes to a set of topological summary statistics using an invariant called the “differential Euler Characteristics (DEC)” transform (Fig. 1(c)). In the fourth step, SINATRA Pro implements a nonlinear Gaussian process model to classify the protein structures using the topological summary statistics, with which association measures are computed for each topological feature to provide a statistical notion of “significance” (Fig. 1(d)). In the last step of the pipeline, SINATRA Pro maps the association measures back onto the original protein structures (Fig. 1(e)), which produces “evidence scores” that reveal the spatial locations that best explain the variance between two protein ensembles. Theoretical details of our implementation are fully detailed in the Materials and Methods sections.

**Figure 1.**
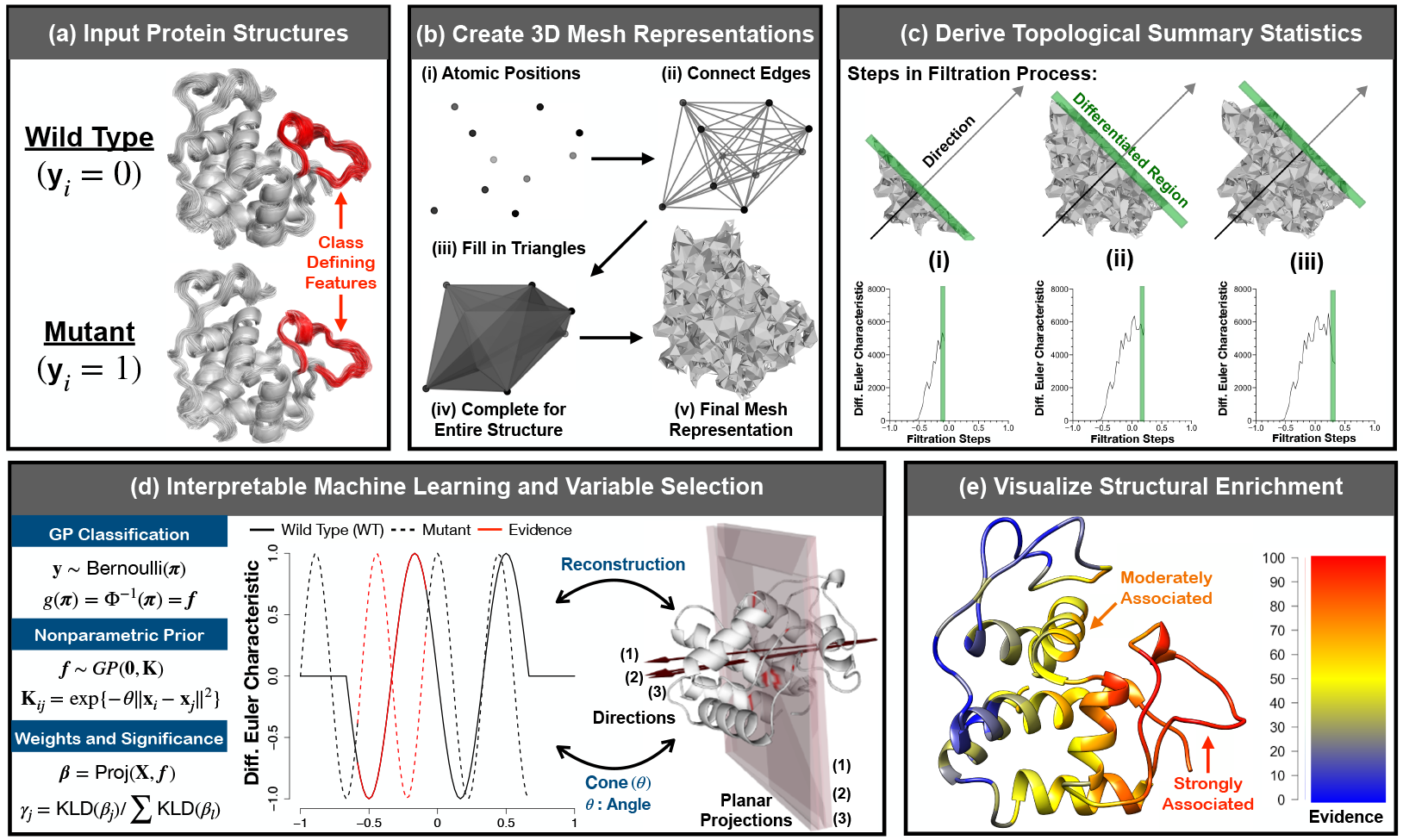
Schematic overview of SINATRA Pro: a novel framework for discovering biophysical signatures that differentiate classes of proteins. **(a)** The SINATRA Pro algorithm requires the following inputs: *(i)* (*x, y, z*)-coordinates corresponding to the structural position of each atom in every protein; *(ii)* **y**, a binary vector denoting protein class or phenotype (e.g., mutant versus wild-type); *(iii) r*, the cutoff distance for simplicial construction (i.e., constructing the mesh representation for every protein); *(iv) c*, the number of cones of directions; *(v) d*, the number of directions within each cone; *(vi) θ*, the cap radius used to generate directions in a cone; and *(vii) l*, the number of sublevel sets (i.e., filtration steps) used to compute the differential Euler characteristic (DEC) curve along a given direction. Guidelines for how to choose the free parameters are given in Table 1. **(b)** Using the atomic positions for each protein, we create mesh representations of their 3D structures. First, we draw an edge between any two atoms if the Euclidean distance between them is smaller than some value *r*, namely dist | (*x*_1_, *y*_1_, *z*_1_), (*x*_2_, *y*_2_, *z*_2_) | *< r*. Next, we fill in all of the triangles (or faces) formed by these connected edges. We treat the resulting triangulated mesh as a simplicial complex with which we can perform topological data analysis. **(c)** We select initial positions uniformly on a unit sphere. Then for each position, we generate a cone of *d* directions within angle *θ* using Rodrigues’ rotation formula [88], resulting in a total of *m* = *c* × *d* directions. For each direction, we compute DEC curves with *l* sublevel sets. We concatenate the DEC curves along all the directions for each protein to form vectors of topological features of length *J* = *l* × *m*. Thus, for a study with *N*-proteins, an *N* × *J* design matrix is statistically analyzed using a Gaussian process classification model. **(d)** Evidence of association measures for each topological feature vector are determined using relative centrality measures. We reconstruct corresponding protein structures by identifying the atoms on the shape that correspond to “statistically associated” topological features. **(e)** The reconstruction enables us to visualize the enrichment of biophysical signatures that best explain the variance between the two classes of proteins. The heatmaps display atomic (or residue-level, which we define as a collection of atoms) evidence potential on a scale from [0 − 100], with a score of 100 meaning most enriched.

### Software Overview

The software for SINATRA Pro requires the following inputs: *(i)* 3D Cartesian coordinates corresponding to the atomic positions in each protein structure; *(ii)* **y**, a binary vector denoting protein class or phenotype (e.g., *y*_*i*_ = 0 for wild-type or *y*_*i*_ = 1 for mutant); *(iii) r*, the cutoff distance for simplicial construction (i.e., constructing the mesh representation for every protein); *(iv) c*, the number of cones of directions; *(v) d*, the number of directions within each cone; *(vi) θ*, the cap radius used to generate directions in a cone; and *(vii) l*, the number of sublevel sets (i.e., filtration steps) used to compute the differential Euler characteristic (DEC) curve along any given direction. In addition to real data analyses, we implement a controlled benchmark simulation study designed to assess SINATRA Pro’s performance at identifyin structurally-perturbed regions in protein dynamics relative to other methods. Results for the controlled benchmark simulations were done using free parameters {*r* = 1.0 Å, *c* = 20, *d* = 8, *θ* = 0.80, *l* = 120}, and results for the real data analyses were done using parameters {*r* = 6.0 Å, *c* = 20, *d* = 8, *θ* = 0.80, *l* = 120}. All values were chosen via a grid search. Guidelines for how to choose the free parameters for the software are given in Table 1. Tables detailing the scalability of the current algorithmic implementation of SINATRA Pro can also be found in Supporting Information (see Tables S1-S3).

**Table 1.** General guidelines for choosing values for the free parameters in the SINATRA Pro pipeline software. The guidelines provided are based off of intuition gained through the simulation studies provided in the main text and Supporting Information. In practice, we suggest specifying multiple cones *c* > 1 and utilizing multiple directions *d* per cone (see monotonically increasing power in Fig. S1 in Supporting Information). While the other two parameters (*θ* and *l*) do not have monotonic properties, their effects on SINATRA’s performance still have natural interpretations. Selection of *θ* ∈ [0.1, 0.8] supports previous theoretical results that cones should be defined by directions in close proximity to each other [13, 15]; but not so close that they explain the same local information with little variation. Note that our sensitivity analyses suggest that the power of SINATRA Pro is relatively robust to the choice of *θ*. Optimal choice of *l* depends on the size of the protein molecules that are being analyzed. Intuitively, for rigid proteins, coarse filtrations with too few sublevel sets cause SINATRA Pro to miss or “step over” structural shifts that occur locally during the course of a molecular dynamic (MD) trajectory. In practice, we recommend choosing the angle between directions within cones *θ* and the number of sublevel sets *l* via cross validation or some grid-based search.

| Free Parameters in SINATRA Pro Software |  |  |  |
| --- | --- | --- | --- |
| Notation | Description | Range | General Guidelines |
| $r$ | Radius cutoff ($\text{\AA}$) for simplicial reconstruction | $[0, \infty)$ | Use smaller $r \leq 2.0 \text{\AA}$ for rigid proteins and $r \in [2.0 \text{\AA}, 6.0 \text{\AA}]$ for flexible proteins. |
| $c$ | Number of cones of directions | $[1, \infty)$ | Set much greater than 1 as more power is generally achieved by taking filtrations over multiple directions |
| $d$ | Number of directions per cone | $[1, \infty)$ | Set much greater than 1 as more power is generally achieved by taking filtrations over multiple directions |
| $\theta$ | Cap radius used to generate directions within a cone | $(0, 2\pi]$ | Set between $[0.1, 0.8]$ since cones should be defined by directions in close proximity |
| $l$ | Number of sublevel sets (filtration steps) | $[1, \infty)$ | Optimal choice depends on the size of protein molecule being analyzed so use grid search |

### Performance of SINATRA Pro on Benchmark Simulations

We implemented a controlled simulation study designed to assess SINATRA Pro’s performance at identifying structurally-perturbed regions in protein dynamics relative to other methods. Here, the premise behind “controlled simulations” is that topological artifacts (i.e., perturbations of atomic positions in a certain region) are manually introduced to a set of protein structures to establish a ground truth and statistically evaluate the concept of power. The original and perturbed structures represent two phenotypic classes and are fed into SINATRA Pro to assess whether it can reliably identify the perturbed regions of interest.

To generate data for these controlled simulations, we use real structural data of wild-type *β*-lactamase (TEM), an enzyme widely implicated in microbial resistance that has evolved numerous mutations of clinical relevance. In the first phenotypic group (set *A*), original structures are drawn at 1 nanosecond (ns) intervals over a 100 ns MD trajectory (e.g., *t*_MD_ = [0, 1, 2, 3, …, 99] ns + *δ*, where *δ* is a time offset parameter). Next, a comparable set of perturbed structures (set *B*) are drawn at 1 ns intervals but shifted by 0.5 ns with respect to the set *A* structures (e.g., *t*_MD_ = [0.5, 1.5, 2.5, 3.5, …, 99.5] ns + *δ*) to allow for thermal noise to be introduced. Here, we displace the atomic positions of each atom in the Ω-loop (i.e., the region of interest or ROI) in each perturbed structure within set *B* by

- a constant Cartesian vector set to *(i)* 0.5 Å, *(ii)* 1.0 Å, and *(iii)* 2.0 Å in each (*x, y, z*) direction;
- a spherically uniform random vector where each (*x, y, z*) direction is first drawn from a standard Gaussian distribution 𝒩 (0, 1) and then the vector is normalized to be of length *(iv)* 0.5 Å, *(v)* 1.0 Å, and *(vi)* 2.0 Å.

These simple, artificial control cases are designed to represent two different forms of structural changes that can happen within protein dynamics. Namely, scenarios *(i)*-*(iii)* involve a displacement of atoms by a constant amount in a constant direction, which emulates a static structural change; while, scenarios *(iv)*-*(vi)* displace the atoms by a constant amount in a (spherically uniform) random direction, which emulates a dynamic or stochastic structural change. Altogether, we use datasets of *N* = 1000 protein structures per simulation scenario: 100 ns intervals *×* 5 different choices of *δ* = {0.0, 0.1, 0.2, 0.3, 0.4} ns *×* 2 phenotypic classes (wild-type versus perturbed). We evaluate all competing methods’ abilities to correctly identify perturbed atoms located within the Omega-loop region (Material and Methods). Here, we use receiver operating characteristic (ROC) curves that plot true positive rates (TPR) against false positive rates (FPR) (Fig. 2). This is further quantified by assessing the area under the curve (AUC). The results presented in the main text reflect using SINATRA Pro with parameters set to {*r* = 1.0 Å, *c* = 20, *d* = 8, *θ* = 0.80, *l* = 120} chosen via a grid search. Note that additional figures assessing how robust SINATRA Pro is to different free parameter value settings can be found in the Supporting Information (see sensitivity analysis in Fig. S1).

**Figure 2.**
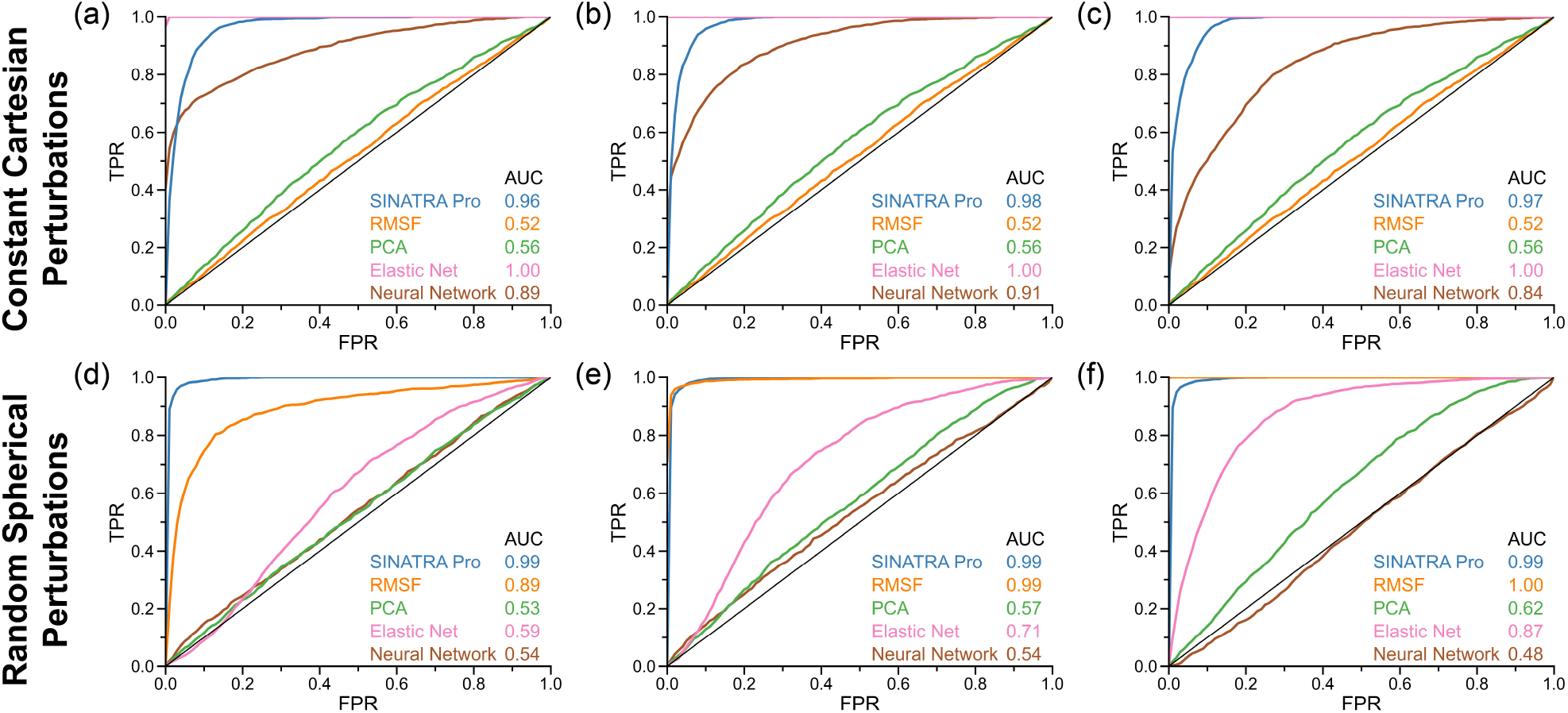
Receiver operating characteristic (ROC) curves comparing the power and robustness of SINATRA Pro to competing 3D mapping approaches in controlled molecular dynamic (MD) simulations. To generate data for these simulations, we consider two phenotypic classes using the real structural data of wild-type *β*-lactamase (TEM). In the first phenotypic class, structural protein data are drawn from equally spaced intervals over a 100 ns MD trajectory (e.g., *t*_MD_ = [0, 1, 2, 3, …, 99] ns + *δ*, where *δ* is a time offset parameter). In the second phenotypic class, proteins are drawn at 1 ns intervals shifted 0.5 ns with respect to the first set (e.g., *t*_MD_ = [0.5, 1.5, 2.5, 3.5, …, 99.5] ns + *δ*) to introduce thermal noise, and then we displace the atomic positions of each atom in the Ω-loop region by **(top row)** a constant Cartesian vector of **(a)** 0.5 Ångströms (Å), **(b)** 1.0 Å, and **(c)** 2.0 Å, or **(bottom row)** by a spherically uniform random vector of **(d)** 0.5 Å, **(e)** 1.0 Å, and **(f)** 2.0 Å. Altogether, we have a dataset of *N* = 1000 proteins per simulation scenario: 100 ns interval *×* 5 different choices *δ* = {0.0, 0.1, 0.2, 0.3, 0.4} ns *×* 2 phenotypic classes (original wild-type versus perturbed). The ROC curves and corresponding area under the curves (AUC) depict the ability of SINATRA Pro to identify “true class defining” atoms located within the Ω-loop region using parameters {*r* = 1.0 Å, *c* = 20, *d* = 8, *θ* = 0.80, *l* = 120} chosen via a grid search. We compare SINATRA Pro to four methods: root mean square fluctuation (RMSF) (orange); principal component analysis (PCA) (green); Elastic Net classification (pink); and a Neural Network (brown). For details on these approaches, see Materials and Methods. A sensitivity analysis exploring the optimal parameter configurations for PCA and Neural Network can be found in Supporting Information (see Figs. S3 and S4, respectively).

Lastly, we want to point out that we do not realign the proteins after structural perturbation has occurred. Realigning the structures after introducing a perturbation poses a slightly different and notably less controlled simulation study. For example, in the constant displacement case, realigning the structures will shift the whole structure against the perturbing vector and result in an unintentional displacement on the opposite side of the structure. A true positive in this case will not be as well-defined as when we keep the unperturbed structure in place and define the perturbed structure as the ground truth for the positive signal. An example of this can be seen in Fig. S2.

#### Overview of Competing Baselines

In this section, we compare SINATRA Pro to four competing approaches: root mean square fluctuation (RMSF) calculations, principal component analysis (PCA), Elastic Net classification, and Neural Network classification. The first baseline is RMSF which computes 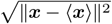, where ***x*** = (*x, y, z*) denotes the positions of the protein’s atoms for each frame and ***x*** is the average position of that corresponding atom over the entire MD simulation. The difference in the RMSF values between the original and perturbed structures is taken as the score for feature selection. The second baseline performs PCA (based on singular value decomposition) over the Cartesian (*x, y, z*)-coordinates for the atoms using `scikit-learn` [18], which reduces the sample space into principal components. We sum the components (weighed by their singular values) for the original wild-type and perturbed data separately, and then determine the magnitudes of the change in the component sum between the two protein classes as the score for feature selection. The last two baselines concatenate the coordinates of all atoms within each protein and treats them as features in a dataframe. The Elastic Net uses a regularized linear classification model via stochastic gradient descent in `scikit-learn` to assign sparse individual coefficients to each coordinate of every atom, where the free regularization parameter is chosen with 90% training and 10% validation set splits. We assess the power of the Elastic Net by taking the sum of the coefficient values corresponding to each atomic position. The Neural Network uses the following architecture with Rectified Linear Unit (ReLU) nonlinear activation functions [19]: *(1)* an input layer of Cartesian coordinates of all of the atoms; *(2)* a hidden layer with *H* = 2048 neurons; *(3)* a second hidden layer with *H* = 512 neurons; *(4)* a third hidden layer with *H* = 128 neurons; and *(5)* an outer layer with a single node which uses a sigmoid link function for protein classification. Batch normalization was implemented between each layer and a normalized saliency map to rank the importance of each atom [20]. The simplest saliency map attributes the partial derivatives *∂y*_*i*_*/∂****x***_*ij*_ as the importance of the coordinates for the *j*-th atom in the *i*-th protein structure; here, *y*_*i*_ denotes the neural network output after the sigmoid link function for the *i*-th protein structure. We then assign global importance to each atom by 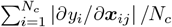, where *N*_*c*_ denotes the number of protein structures in a given class. For the Neural Network, we assess power by taking the sum of the saliency map values corresponding to each atomic position. To ensure that both PCA and the Neural Network are evaluated with their optimal parameter configurations, we also performed variants of these approaches in their own sensitivity analysis. The results for these variants of PCA and the Neural Network can be found in Supporting Information (see Figs. S3 and S4, respectively).

It is important to note that, while SINATRA Pro is implemented over the entire protein structure, the four baselines that we consider are limited to only assessing structural differences between atoms with correspondences between the two datasets — that is, for the competing baselines, we use all “pairable” atoms and omit mutated residues which be considered “unpairable” between structures. The main reason for this is that atomic features can be gained or lost due to mutations or phylogenetic variations that introduce heterogeneity in protein sequences, thus creating a lack of a one-to-one correspondence between any two given 3D structures. Without this explicit mapping between structures, none of the four coordinate-based competing approaches are able to be fully implemented as they all rely on (in some way or another) equal dimensionality across all proteins. Therefore, when assessing MD trajectories, the corresponding or “pairable” atoms represent consistent “landmarks” that summarize the global geometry of the protein structure. Ultimately, we recognize that these method comparisons with SINATRA Pro are not equivalent; however, they do highlight a key and practical advantage of the topological data analytic approach used in SINATRA Pro which maintains its utility even when such atom-by-atom correspondences between protein structures are not available.

#### Method Comparisons

The overall performance of each competing method to identify is dependent on two factors: *(1)* whether the structural changes are reproduced by static or stochastic conformations, and *(2)* the underlying statistical assumptions of the methods. For example, RMSF had the most difficulty identifying constant displacements in the protein structures (Figs. 2(a)-(c)). In these scenarios, RMSF was effectively a random classifier with an average AUC ≈ 0.5 and diagonal ROC curves showing no signal detected. These results are explained by that fact that RMSF effectively measures how much each atomic coordinate ***x*** deviates from the average atomic position in the ensemble ***x***. When conformations are constantly shifted, ***x*** and ***x*** are scaled by the same factor and, as a result, their differences remain unchanged. Therefore, static structural changes are essentially undetectable by RMSF. On the other hand, RMSF is perfectly well suited for stochastic structural changes because, when atomic displacements are caused by a random spherical vector, the scaling factors between each ***x*** and ***x*** are noticeably different (AUC ≥ 0.89 in Figs. 2(d)-(f)). Note that PCA follows a similar trend, but with much less power likely due to the fact that we only consider the top 10 PCs of each protein trajectory in isolation for these analyses. One potential improvement for the PCA approach (at least for the larger perturbation scenarios) could be to first treat the original and perturbed structures as a single dataset and compute the top PCs over the concatenated protein trajectories. We leave this exploration to the reader.

A slightly different intuition can be followed when looking at the results for the Elastic Net and Neural Network classifiers. When atomic positions are shifted equally by a constant Cartesian vector, the atoms in the ROI for the perturbed proteins become (in some cases) completely separable from those in the original structures. Therefore, an Elastic Net and Neural Network have no trouble assigning the true causal atoms non-zero effect sizes (AUC ≥ 0.85 for both approaches in Figs. 2(a)-(c)). This observation is similar to previous works which show coordinate-based regularization to be most effective when variation between 3D structures occurs on a global scale and in the same direction on the unit sphere [13]. In the cases of random spherical perturbations, the variance of the distribution of atoms in the ROI widens; hence, the Elastic Net and Neural Network have a more difficult time identifying features that differentiate two protein classes, unless those variations happen on a global scale (again see Figs. 2(d)-(f)).

Most notably, SINATRA Pro performs consistently well in all simulation scenarios, identifying both static and dynamic differences better than most of the competing baselines that we considered (AUC ≥ 0.96 in Fig. 2). Although SINATRA Pro is not as adept as the Elastic Net (AUC = 1.00 in Figs. 2(a)-(c)) at detecting static changes, it is able to robustly select significant features that are ignored by RMSF. In addition, SINATRA Pro is much better than the Elastic Net and Neural Network at identifying significant spherical perturbations that arise dynamically between protein structures. We hypothesize that summarizing atomic positions with Euler statistics is what enables SINATRA Pro to robustly capture both varying topology and geometry, unlike its coordinate-based counterparts, regardless of whether those differences occur in a constant or stochastic way.

### Detecting Conformational Changes in Real Protein Systems

To examine SINATRA Pro’s ability to identify known structural changes of biological significance in real data, we consider the following five protein systems (Table 2): *(1)* the wild-type and Arg164Ser mutant of TEM *β*-lactamase; *(2)* the wild-type and Ile50Val mutant of HIV-1 protease; *(3)* the guanosine triphosphate (GTP) and guanosine diphosphate (GDP) bound states of EF-Tu; *(4)* the wild-type and Met290Ala mutant of the Abl1 tyrosine protein kinase; and *(5)* unbound and IBB-bound states of importin-*β*. We choose to analyze these particular systems because they undergo varying degrees of conformational changes that have been well-studied in the literature (again see Table 2). Here, we will treat these previously identified features as ROIs, where the assumed “difficulty” for SINATRA Pro to statistically resolve structural signatures will be based on the stochasticity observed within each protein system. Namely, it will be more difficult to perform feature selection on structural ensembles that are highly dynamic as spurious fluctuations can interfere with detecting signal from the ROI. For each protein system, three MD trajectories were generated. From each of these trajectories, ten different series of structures are sampled, where each series is separated by 0.1 ns and each structure in a series is separated by 1 ns. SINATRA Pro is then implemented on the ten different series of structures extracted from the same trajectory data to mitigate the impact of variability between MD simulations (see Material and Methods).

**Table 2.** Detailed overview of the different protein systems analyzed in this study. The columns of this table are arranged as follows: (1) the name of each protein studied; (2) the corresponding Protein Data Bank (PDB) ID for each molecule [94]; (3) the known chemical change or mutation type that is considered; (4) the specific structural signatures that are known to be associated with each chemical change or mutation type; (5) the presumed difficulty level for SINATRA Pro to detect each structural signature based on the homogeneity in shape variation between the wild-type and mutant proteins; and (6) references that have previously suggested some level of association or enrichment between each structural change and the mutation of interest.

| Protein | PDB ID | Chemical Change | Structural Signature | Difficulty | Ref(s) |
| --- | --- | --- | --- | --- | --- |
| β-lactamase (TEM) | 1BTL | Arg164Ser | Increased dynamics of Ω-Loop (Residues 163-178) | Easy | [21, 22] |
| HIV-1 Protease | 3NU3 | Ile50Val | Reduced stability in the flaps (Residues 47-55) | Medium | [29, 31, 32] |
| EF-Tu | 1TTT | GTP Hydrolysis | Increased flexibility of Domain 2 (Residues 208-308) | Easy | [37, 89, 90] |
| Abl1 | 3KFA | Met290Ala | Fluctuations in the DFG motif and displacement of helix αC | Hard | [2, 43, 91–93] |
| Importin-β | 2P8Q | IBB Release | Uncoiling in the conformation of the superhelix | Hard | [45–47] |

Atomic enrichments are illustrated in Figs. S5 and S6, while residue-level structural enrichments are shown in Figs. 3-5 and Figs. S7-S21. To quantitatively assess the probability that SINATRA Pro is identifying any given ROI by chance, we implement a null region hypothesis test to estimate a *P*-value and an approximate Bayes factor (BF) corresponding to our power to reliably and robustly select certain features (Material and Methods). Reported results for the *P*-values and BF calculations are based on all MD simulated structures and can be found in Table 3. For comparison, we again implement the RMSF (Figs. 3-5 and Figs. S12, S18, and S20-S21) and Elastic Net (Figs. S8, S11, S14, S16, and S19) baselines on all atomic positions with correspondences within these same protein systems. Here, we use scatter plots to illustrate the correlation between how each of these methods and SINATRA Pro rank the variable importance of the “pairable” atoms. All results presented in the main text reflect using SINATRA Pro with parameters set to {*r* = 6.0 Å, *c* = 20, *d* = 8, *θ* = 0.80, *l* = 120} chosen via a grid search.

**Figure 3.**
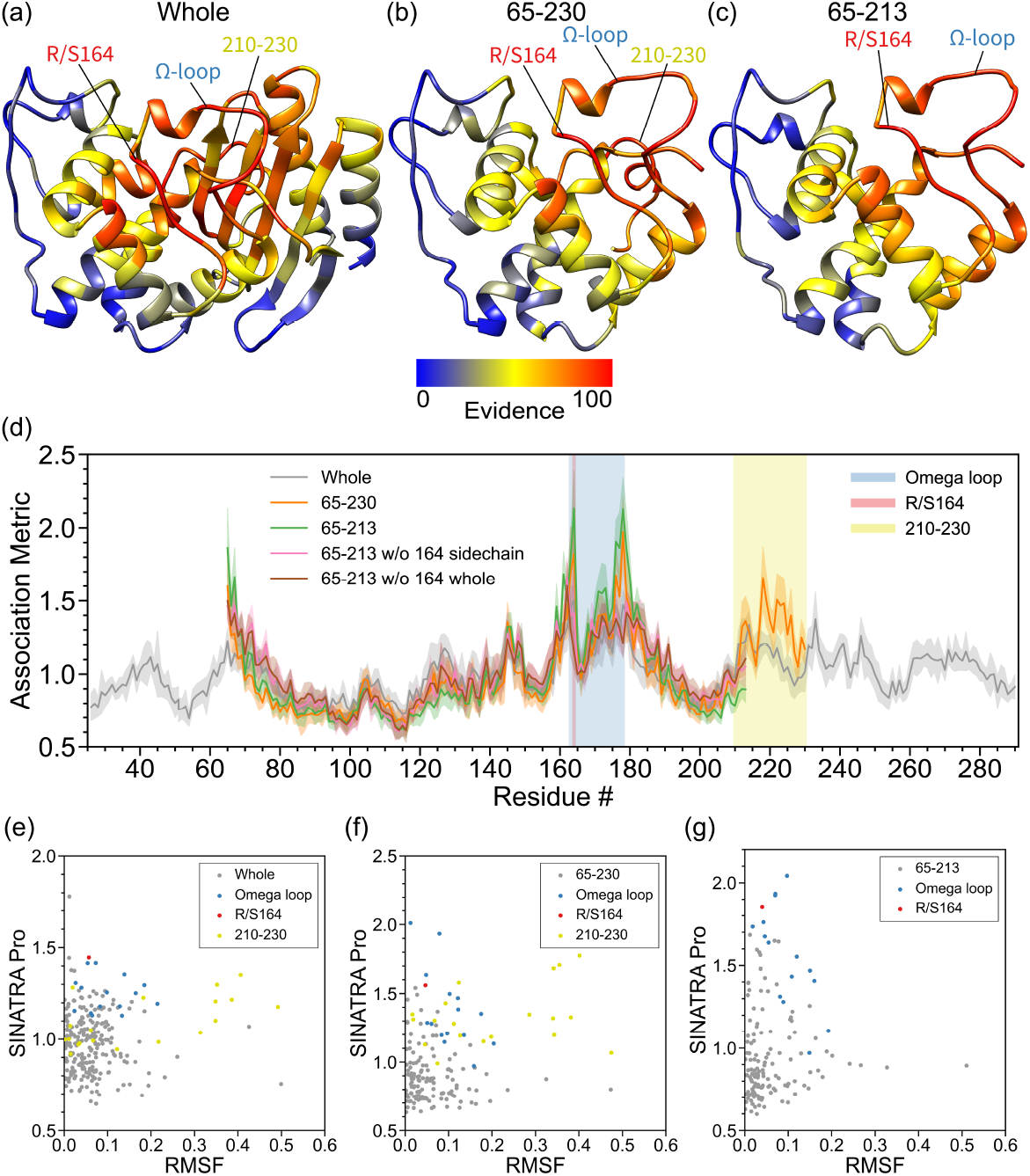
Real data analyses aimed at detecting structural changes in the Ω-loop of *β*-lactamase (TEM) induced by an R164S mutation. In this analysis, we compare the molecular dynamic (MD) trajectories of wild-type *β*-lactamase (TEM) versus R164S mutants [21, 22]. For both phenotypic classes, structural data are drawn from equally spaced intervals over a 100 ns MD trajectory (e.g., *t*_MD_ = [0, 1, 2, 3, …, 99] ns + *δ*, where *δ* is a time offset parameter). Altogether, we have a final dataset of *N* = 2000 protein structures in the study: 100 ns long interval × 10 different choices *δ* = {0.0, 0.1, 0.2, …, 0.9} ns × 2 phenotypic classes (wild-type versus mutant). This figure depicts results after applying SINATRA Pro using parameters {*r* = 6.0 Å, *c* = 20, *d* = 8, *θ* = 0.80, *l* = 120} chosen via a grid search. The heatmaps in panels **(a)**-**(c)** highlight residue evidence potential on a scale from [0 100]. A maximum of 100 represents the threshold at which the first residue of the protein is reconstructed, while 0 denotes the threshold when the last residue is reconstructed. Panel **(a)** shows the residue-level evidence potential when applying SINATRA Pro to the whole protein, while panels **(b)** and **(c)** illustrate results when strictly applying the SINATRA Pro pipeline to atoms in residues 65-230 and 65-213, respectively. Annotated regions of interest are color coded and correspond to the shaded residue windows in panel **(d)**. Panel **(d)** shows the mean association metrics (and their corresponding standard errors) computed for each residue within each analysis (see Material and Methods). Here, the overlap shows the robustness of SINATRA Pro to identify the same signal even when it does not have access to the full structure of the protein. The final row plots the correlation between the SINATRA Pro association metrics and the root mean square fluctuation (RMSF) for all atoms with correspondences in the **(e)** whole protein, **(f)** fragment 65-230, and **(g)** fragment 65-213.

**Figure 4.**
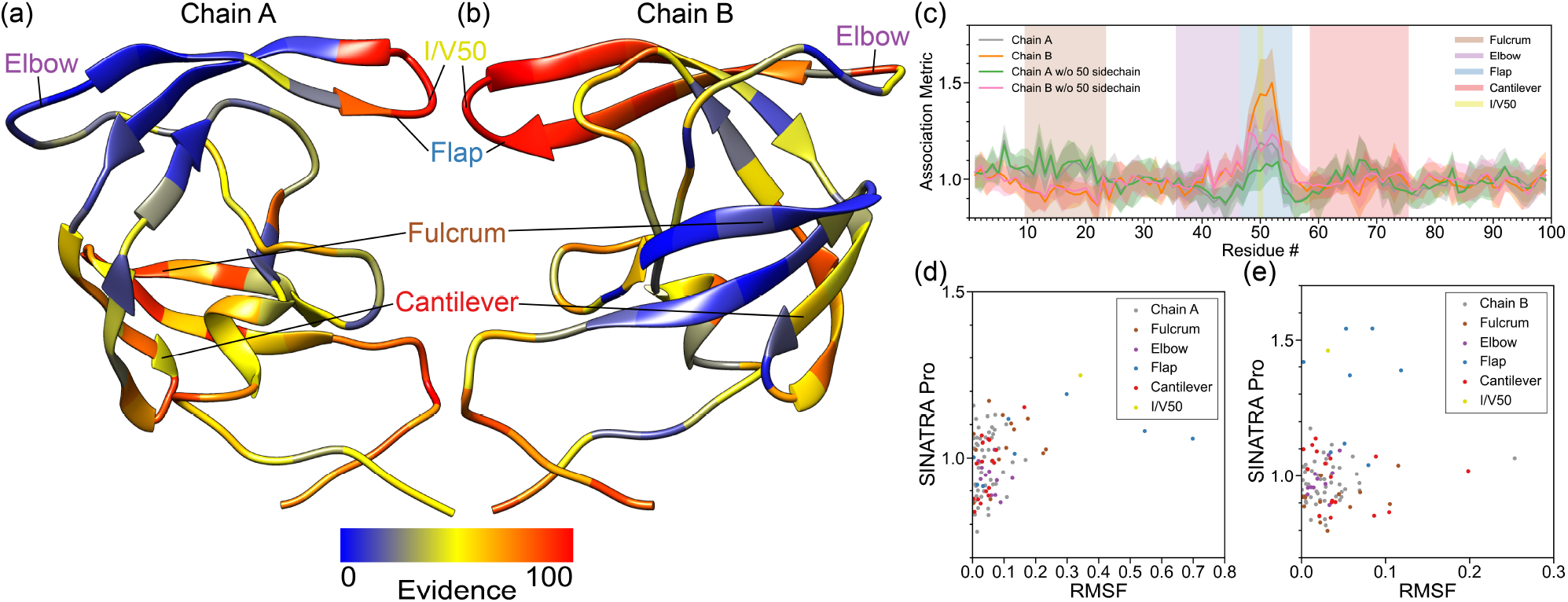
Real data analyses recover structural changes in the flap region of HIV-1 protease driven by a Ile50Val mutation. In this analysis, we compare the molecular dynamic (MD) trajectories of wild-type HIV-1 protease versus Ile50Val mutants (i.e., within residues 47-55). For both phenotypic classes, structural data are drawn from from equally spaced intervals over a 100 ns MD trajectory (e.g., *t*_MD_ = [0, 1, 2, 3, …, 99] ns + *δ*, where *δ* is a time offset parameter). Altogether, we have a final dataset of *N* = 2000 proteins in the study: 100 ns long interval × 10 different choices *δ* = {0.0, 0.1, 0.2, …, 0.9} ns × 2 phenotypic classes (wild-type versus mutant). This figure depicts results after applying SINATRA Pro using parameters {*r* = 6.0 Å, *c* = 20, *d* = 8, *θ* = 0.80, *l* = 120} chosen via a grid search. The heatmaps in panels **(a)** and **(b)** highlight residue evidence potential on a scale from [0 100]. A maximum of 100 represents the threshold at which the first residue of the protein is reconstructed, while 0 denotes the threshold when the last residue is reconstructed. Panel **(a)** shows residue-level evidence potential when applying SINATRA Pro to chain A, while panel **(b)** depicts results for chain B. Annotated regions of interest are color coded and correspond to the shaded residue windows in panel **(c)**. Panel **(c)** shows the association metrics (and their corresponding standard errors) computed for each residue in chains A and B, with and without the 50th residue’s side chain being included in the analysis (see Material and Methods). Here, the overlap shows the robustness of SINATRA Pro for identifying the same signal even when it does not have access to the full structure of the protein. The final row plots the correlation between the SINATRA Pro association metrics and the root mean square fluctuation (RMSF) for all atoms with correspondences in **(d)** chain A and **(e)** chain B, respectively. Highlighted are all atoms with correspondences found in regions of the protein corresponding to the fulcrum (brown), elbow (purple), flap (blue), cantilever (red), and I/V50 (yellow) [29, 31, 32].

**Figure 5.**
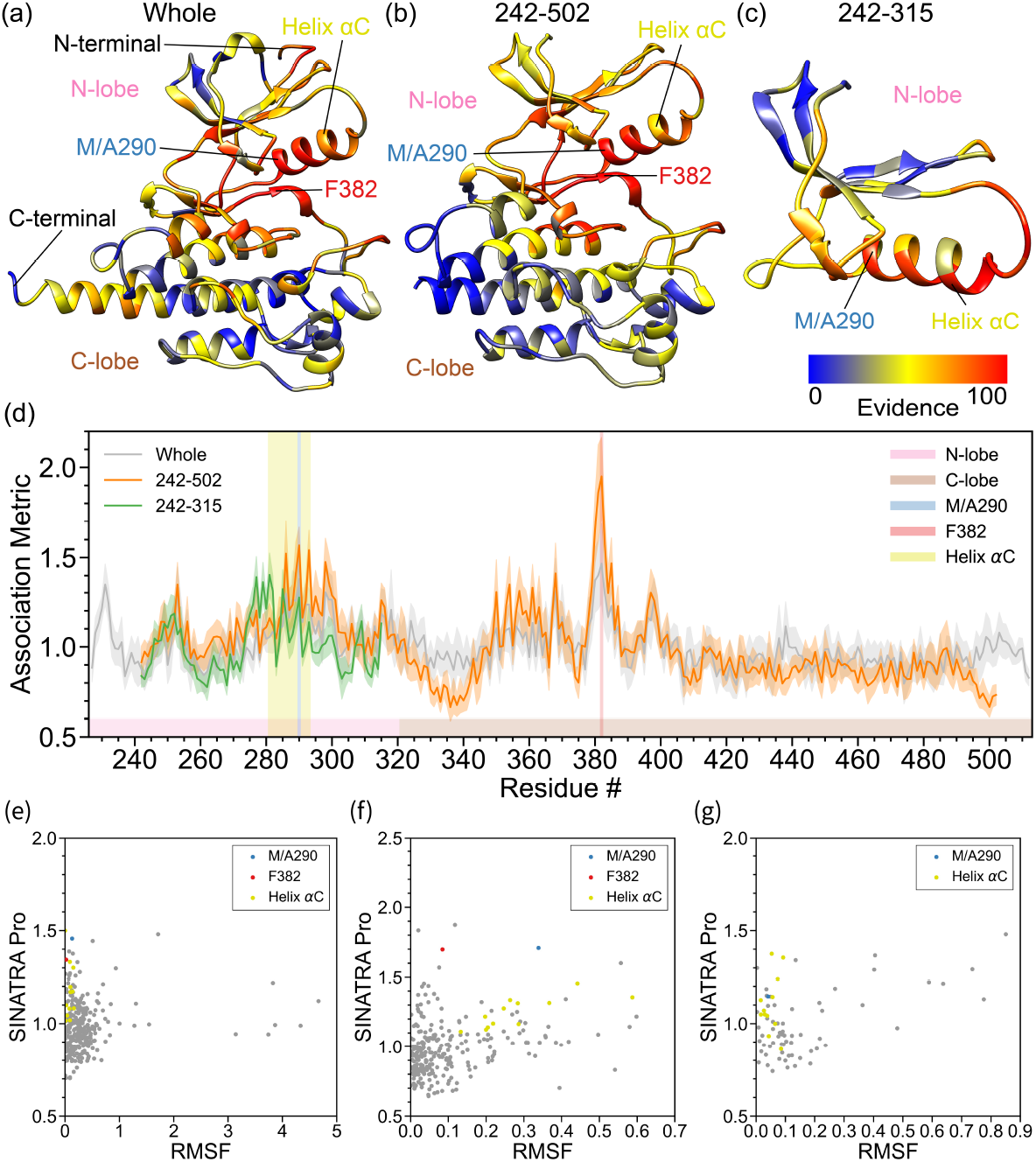
Real data analyses identify enrichment in the N-terminal pocket of the Abl1 Tyrosine protein kinase due to a M290A mutation in the *α*C helix. In this analysis, we compare the molecular dynamic (MD) trajectories of wild-type Abl1 versus M290A mutants [2, 43, 91–93]. For both phenotypic classes, structural data are drawn from equally spaced intervals over a 150 ns MD trajectory (e.g., *t*_MD_ = [0, 1, 2, 3, …, 99] × 1.5 ns + *δ*, where *δ* is a time offset parameter). Altogether, we have a final dataset of *N* = 3000 proteins in the study: 150 ns long interval × 15 different choices *δ* = {0.0, 0.1, 0.2, …, 1.4} ns × 2 phenotypic classes (wild-type versus mutant). This figure depicts results after applying SINATRA Pro using parameters {*r* = 6.0 Å, *c* = 20, *d* = 8, *θ* = 0.80, *l* = 120} hosen via a grid search. The heatmaps in panels **(a)**-**(c)** highlight residue evidence potential on a scale from [0 − 100]. A maximum of 100 represents the threshold at which the first residue of the protein is reconstructed, while 0 denotes the threshold when the last residue is reconstructed. Panel **(a)** shows residue-level evidence potential when applying SINATRA Pro to the whole protein, while panels **(b)** and **(c)** illustrate results when strictly applying the SINATRA Pro pipeline to atoms in residues 242-502 and 242-315, respectively. Annotated regions of interest are color coded and correspond to the shaded residue windows in panel **(d)**. Panel **(d)** shows the association metrics (and their corresponding standard errors) computed for each residue within each analysis (see Material and Methods). Here, the overlap shows the robustness of SINATRA Pro to identify the same signal even when it does not have access to the full structure of the protein. The final row plots the correlation between the SINATRA Pro association metrics and the root mean square fluctuation (RMSF) for all atoms with correspondences in the **(e)** whole protein, **(f)** fragment 242-502, and **(g)** fragment 242-315.

**Table 3.** Null hypothesis experiment to evaluate SINATRA Pro’s ability to find regions of interest (ROI) in each of the proteins analyzed in this study. Here, we assess how likely it is that SINATRA Pro finds the region of interest (ROI) by chance. These ROIs include: *(i)* the Ω-loop (residues 163-178) in TEM; *(ii)* the flap region (residues 47-55) in HIV-1 protease; *(iii)* Domain 2 (residues 208-308) in EF-Tu; and *(iv)* the DFG motif (residues 381-383) in Abl1. Note that protein structures were only analyzed if they contained an entire ROI. For example, in the context of Importin-*β*, the superhelix includes the entire structure and so we do conduct a null analysis. In this experiment, to produce the results above, we generate “null” regions on each protein using a *K*-nearest neighbors (KNN) algorithm on different atoms as random seeds [84], and exclude any generated regions that overlap with the ROI. Next, for each region, we sum the association metrics of all its atoms. We compare how many times the aggregate scores for the ROI are higher than those for the null regions. These “*P*-values,” and their corresponding calibrated Bayes factors (BF) when the computed *P <* 1*/e*, are provided above. Note that *P*-values less than the nominal size 0.05 and BFs greater than 2.456 are in bold. Results above are based on SINATRA Pro using parameters {*c* = 20, *d* = 8, *θ* = 0.80, *l* = 120} while varying the radius cutoff parameter *r* for mesh construction on each protein structure.

| | | | $r = 2 \text{ \AA}$ | | $r = 4 \text{ \AA}$ | | $r = 6 \text{ \AA}$ | |
| --- | --- | --- | --- | --- | --- | --- | --- | --- |
| Protein | ROI | Fragment | $P$-value | Bayes Factor | $P$-value | Bayes Factor | $P$-value | Bayes Factor |
| TEM | Ω-Loop | Whole | $5.95 \times 10^{-1}$ | — | $3.35 \times 10^{-4}$ | **137.121** | $5.63 \times 10^{-2}$ | 2.270 |
| | | 65-230 | $1.20 \times 10^{-1}$ | 1.447 | $4.16 \times 10^{-2}$ | **2.783** | $6.85 \times 10^{-2}$ | 2.004 |
|  |  | 65-213 | **$7.22 \times 10^{-4}$** | **70.438** | **$7.22 \times 10^{-4}$** | **70.438** | **$7.22 \times 10^{-4}$** | **70.438** |
| HIV-1 | Flap | Chain A | $2.33 \times 10^{-1}$ | 1.084 | $4.03 \times 10^{-2}$ | **2.841** | $2.95 \times 10^{-1}$ | 1.022 |
|  |  | Chain B | **$8.14 \times 10^{-4}$** | **63.554** | **$8.14 \times 10^{-4}$** | **63.554** | **$8.14 \times 10^{-4}$** | **63.554** |
| EF-Tu | Domain 2 | Whole | **$9.30 \times 10^{-4}$** | **56.657** | **$9.30 \times 10^{-4}$** | **56.657** | **$9.30 \times 10^{-4}$** | **56.657** |
| Abl1 | DFG Motif | Whole | $1.94 \times 10^{-1}$ | 1.157 | $5.38 \times 10^{-1}$ | — | **$8.86 \times 10^{-3}$** | **8.783** |
| | | 242-502 | $1.54 \times 10^{-1}$ | 1.279 | **$2.50 \times 10^{-4}$** | **177.614** | **$2.50 \times 10^{-4}$** | **177.614** |

Note that we also provide additional figures assessing how robust SINATRA Pro is to different configurations of protein meshes for these data in the Supporting Information. For example, results analyzing SINATRA Pro with protein meshes constructed under different radius cutoffs *r* values can be found in Figs. S22-S26. To ensure sampling consistency of the 100 ns results, we performed sensitivity analyses based on protein structures drawn from different lengths up to 200 ns MD simulation trajectories (Figs. S7, S10, S13, S15, and S17). Lastly, we test the robustness of SINATRA Pro when the protein structures have been aligned in two different ways (Material and Methods). In the first, we align protein structures by minimizing the root-mean-square distance (RMSD) between their atoms (which we focus on in the main text); while, in the second, we perform sequence-independent structural alignment where we *implicitly* normalize the 3D protein structures by rotationally aligning their topological summary statistics (i.e., alignment without taking into correspondence between structures). Figures comparing the correlation between SINATRA Pro results on the RMSD pre-aligned protein structures versus using the topological rotational alignment can be found in Figs. S27 and S28.

### Conformational Changes in the Active Site and Regulatory Ω-Loop of Arg164Ser TEM *β*-lactamase

Previous studies suggest that the Arg164Ser mutation in *β*-lactamase (TEM) induces structural changes in a highly plastic region known as the Ω-loop (residues 163-178), which plays a major role in the regulation of enzymatic activity [21, 22]. In wild-type *β*-lactamase, Arg164 makes a salt bridge with Asp179 that “pins down” the Ω-loop. Mutating Arg164 to serine breaks this salt bridge and disrupts a vast network of electrostatic and hydrogen interactions, dramatically affecting the dynamical behavior of the area surrounding the loop, parts of the active site, and potentially other protein domains [23]. These dynamical rearrangements confer multi-drug resistance to bacteria expressing TEM Arg164Ser, allowing them to hydrolyze a large number of cephalosporin antibiotics such as ceftazidime, cefixime, and cefazolin, in lieu of hydrolyzing ampicillin [24]. Given the enormous burden of multiresistant bacteria on public health, it is important that we understand the molecular mechanisms behind the structural rearrangements responsible for the transition to the cephalosporinase phenotype in order to orient future antibiotic design. Although previous studies have probed these rearrangements with varying approaches [23], the full mechanism remains elusive, highlighting the need for novel sampling and analytical methods that can detect the very slight changes in TEM’s active site topology that lead to drug resistance in the Arg164Ser and similar mutants. To help bridge this gap in understanding, we ran all-atom MD simulations of unbound TEM-1 and its Arg164Ser mutant, generated by homology modeling, and analyzed the results using SINATRA Pro, RMSF, and the Elastic Net baselines. Here, we expect SINATRA Pro to reveal new insights about the molecular mechanisms underlying the specificity shift precipitated by the Arg164Ser mutation, due its ability to detect both minute static and stochastic changes in topology that elude traditional methods.

We compare the MD trajectories of wild-type and mutant TEM using aligned structures of the whole protein (Fig. 3(a) and Fig. S5(a)), residues 65-230 (Fig. 3(b) and Fig. S5(b)), and residues 65-213 (Fig. 3(c) and Fig. S5(c). In all three cases, statistical association measures from SINATRA Pro suggest that there are indeed significant structural changes in the Ω-loop (residues 163-178) relative to the rest of the regions in the protein (Fig. 3(d)), especially on residues 164 and 176-179, which are involved in the electrostatic interaction networks disrupted by the arginine to serine substitution. This ROI is not as prominently identified by the RMSF (see scatter plots in Fig. 3(e)-(g)) or the Elastic Net baselines (Fig. S8(b)-(d)). Alternatively, all three approaches were able to identify the region harboring residues 213-230, which undergoes a noticeable dynamic shift over the course the MD trajectory. These results are consistent with our controlled simulations, which showed that the only time that RMSF and the Elastic Net both exhibit relatively decent power for stochastic changes is when large structural deviations are introduced (e.g., see power comparisons in Fig. 2(f)).

To more thoroughly assess if the Arg164Ser mutation contributes to the detected changes, we removed the Arg/Ser164 sidechain, as well as the whole residue (backbone and side chain), from our analyses. With the Arg/Ser164 atoms removed, association metrics of Arg/Ser164 and residues 176-179 diminished, which implies that signals pertaining to the dynamical contributions from the electrostatic interaction networks mediated by the side-chains of Arg164 and Glu179 are lost due to the missing topology. However, enrichment in the Ω-loop persisted, affirming that the identified topological differences are not just due to changes in these atoms. The null region test showed that the Ω-loop is indeed a robust significant structural feature in TEM, with *P* = 5.63 × 10^*−*2^ and BF = 2.27 when the whole TEM protein is analyzed (Table 3 with *r* = 6.0 Å), *P* = 6.85 × 10^*−*2^ and BF = 2.00 when residues 65-230 are analyzed, and *P* = 7.22 × 10^*−*4^ and BF = 70.4 when residues 65-213 are analyzed. We hypothesize that the ROI *P*-value is larger than the nominal 0.05 level for analyses with the whole structure and residues 65-230 because movement in the Ω-loop occurs jointly with moderate fluctuations in the region harboring residues 210-230. Overall, when we limit our scope to just residues 65-213, the region test robustly rejects the null hypothesis of the Ω-loop being identified by chance. It is worth noting that these results also highlight that there can be variations in the strength of signal detected by SINATRA Pro depending on if one carries out analyses on whole protein structures versus on different fragments. These variations are most likely due to bias caused by different structural segments having different centers of mass and reference points during alignment. This will cause differences in the topological summary statistics that are computed for each structure and will lead to slightly varying signals in the biophysical signatures detected in downstream SINATRA Pro analyses.

Our results are particularly interesting for the TEM *β*-lactamase example because they highlight the importance of codon positions 164 and 179 in controlling Ω-loop dynamics, which contributes to modulating activity. Moreover, SINATRA Pro correctly captures the topological effects of the disruption of the electrostatic network formed by Arg164, Arg178, and Asp179 due to the Arg164Ser mutation. In addition to reaffirming previously observed phenomena, SINATRA Pro also identified meaningful shifts in the 210-230 segment in response to the resistance-granting Arg164Ser mutation. This suggests that the topology of the 210-230 segment, which forms the upper boundaries of the active site, is tightly correlated with shifts in the Ω-loop. Our results suggest an additional potential mechanism for activity modulation by Ω-loop fluctuations, where topological changes propagate from regulatory loops to parts of the active site, suggesting potential allosteric couplings between the Ω-loop and the 210-230 segment. These results function as a testament to SINATRA Pro’s capacity for distinguishing meaningful topological differences from the random fluctuations introduced by disorder-inducing mutations such as Arg164Ser, which obfuscates traditional analyses pipelines.

### Changes in the Flap Region of HIV-1 Protease Driven by the Ile50Val Mutation

Our next analysis focuses on the HIV-1 protease, an enzyme that is essential for viral reproduction and is a well-established target for controlling HIV infections [25]. *In vivo*, the protease cleaves the HIV polyproteins Gag and Gag-Pol at multiple sites, creating the mature protein components of an HIV virion [26]. Over the past 25 years, ten HIV protease inhibitors have been approved for human use by the Food and Drug Administration (FDA), with many more undergoing clinical trials [27]. Similar to TEM, point mutations in the protease gene lead to products that are considerably less susceptible to inhibition by current drugs, generating drug-resistant HIV variants that pose a considerable risk [28]. Many hypotheses have been proposed for the molecular mechanisms underlying the most common resistance-granting mutations, and recent studies have used sophisticated geometric analyses to classify conformational ensembles of mutant structures based on their influences on the dynamics [29]. Structurally, the HIV protease forms a homodimer with highly ordered domains [30]. Most resistance-granting mutations, such as the Ile50Val substitution, are thought to mainly affect the cross-correlated fluctuations of the flaps (residues 47-55), imparting minute changes to the fulcrum and lateral topology [29, 31, 32]. These findings suggest that mutations such as Ile50Val effectively rewire residue communication networks, significantly reducing its affinity for binding the inhibitor. These structural rearrangements lead to surprisingly nuanced changes to the topology, which as discussed previously, require refined quantitative methods to be detected. To test SINATRA Pro’s performance in detecting these small changes, we ran all-atom molecular dynamics simulations of “protein and ligand complex in water” systems containing HIV Protease or its Ile50Val mutant complexed with the antiviral drug Amprenavir [33]. We then followed that with analysis using either SINATRA Pro, RMSF, or the Elastic Net, with the objective of measuring each routine’s capacity for detecting and reporting the topological changes induced by the mutation.

Even though MD simulations are performed on the protein’s native dimeric form, chains A and B were separately selected and aligned before being input into each statistical method to avoid alignment bias due to inter-chain orientation. This focuses SINATRA, RMSF, and the Elastic Net on identifying the structural differences within each chain (e.g., Fig. 4(a)-(b) and Fig. S5(d)-(e)). However, note that results for the analyses on the dimeric structures can be found in the Supporting Information (Fig. S9). Overall, our analyses reveal that chains A and B seem to respond asymmetrically to the backbone effects of the mutation within the timeframe of the simulations (Table 3). This is not unexpected, as during the course of the simulations, the inhibitor Amprenavir affects dynamics asymmetrically by interacting more significantly with residues of Chain A. The change in RMSF for most of the residues in the flap are shown to be greater than 0.2 Å for chain A and smaller than 0.2 Å for chain B, indicating that the flap became more dynamic in the MD simulations when the Ile50Val mutation was introduced into chain A (Fig. 4(d)-(e) and Fig. S20(b)). Meanwhile, the Elastic Net shows larger nonzero coefficients in the fulcrum for chain A than in chain B (Fig. S11(a)).

While association metrics from SINATRA Pro identify structural changes in the flap for both chains (Fig. 4(c)), they also capture the geometric shifts within the fulcrum for chain A. We hypothesize that the coexistence of the two changes (flap and fulcrum) in chain A contributes to a smaller peak (i.e., a weaker signal) in the association metrics produced by SINATRA Pro in the flap for chain A than in chain B. This asymmetry is confirmed by the null test, as topological changes in the flap appear to be less statistically significant in chain A (*P* = 2.95 × 10^*−*1^ and BF = 1.022) than in chain B (*P* = 8.14 × 10^*−*4^ and BF = 63.554) for this MD simulation data. Similar to *β*-lactamase, we assess if the Ile50Val mutation contributes to these detected topological changes. Upon removing the Ile/Val50 side-chain, the signal observed by SINATRA Pro in the flap drops with the missing topology, but still displays a significant peak relative to the rest of the protein, which implies that the change in association scores is not solely due to the structural differences upon introducing the Ile50Val mutation (Fig. 4(c)). Although SINATRA Pro clearly identified the effects of the isoleucine to valine substitution in flap topology and fulcrum dynamics, SINATRA Pro did not detect other previously elucidated structural signatures of the mutation, such as lateral extension [29]. As the baseline approaches also failed to identify these features, it is likely that their absence stems from sampling limitations inherent to the brute-force and relatively short production dynamics used to generate the conformational datasets.

The HIV protease system presents a good test case for SINATRA Pro due to its relative structural simplicity and the symmetry of the dimer. Encouragingly, SINATRA Pro’s results closely match those observed in previous studies that sought to characterize deltas in the backbone dynamics of the HIV protease in response to resistance-granting mutations [29].

### Domain 2 in EF-Tu Undergoes Structural Changes upon GTP Hydrolysis

In our third analysis, we focus on EF-Tu (elongation factor thermo unstable), which is a G-protein that is responsible for catalyzing the binding of aminoacyl-tRNAs to the ribosome in prokaryotes. After binding GTP and a given aa-tRNA, EF-Tu strongly interacts with the ribosomal A site [34]. Following productive aa-tRNA binding, EF-Tu is released upon GTP hydrolysis [35]. The resultant GDP molecule is exchanged for GTP with EF-Ts (elongation factor thermo stable), allowing elongation to continue. Structurally, EF-Tu is composed of a Ras-like catalytic domain (RasD), common to G-proteins, and two beta-barrel domains (D2 and D3) [36]. Previous studies probing dynamic fluctuations of GTPases have identified that, after hydrolysis (in the GDP-bound state), EF-Tu shows considerably increased flexibility of backbone atoms belonging to Domains 2 and 3, which are downstream of the nucleotide binding site in RasD [37]. These fluctuations are thought to be correlated with conformational rearrangements required for the exchange of GDP for GTP [37]. The conformational rearrangements are thought to occur on multiple millisecond timescales [37], presenting an obstacle for their study using all-atom molecular dynamics simulations. While the full relaxations associated with the change in ligand chemistry are challenging to sample effectively, we expect SINATRA Pro to have the ability to detect subtle topological differences arising from rearrangements of residue interacting networks in response to the chemical change in the ligand.

To compare our method’s performance to that of alternative techniques (Figs. S12 and S14), we run SINATRA Pro, RMSF, and the Elastic Net on the whole structure (Figs. S5(f) and S12(a)) and fragment windows limited to residues 208-308 (Figs. S5(g) and S12(b)) and 311-405 (Figs. S5(h) and S12(c)). Note that all figures displaying the enrichment of structural features are projected onto the GTP-bound structures. The evidence scores from SINATRA Pro reveal significant structural changes at Domain 2, with minimal structural changes in the majority of the Ras-like Domain, which agrees with findings in previous studies [37]. The null region test shows that Domain 2 is indeed an important structural feature in EF-Tu identified by SINATRA Pro with *P* = 9.30 × 10^*−*4^ and BF = 56.657, which robustly rejects the null hypothesis of the ROI being identified by chance (Table 3).

The chemical changes associated with the substitution of GTP with GDP in the EF-Tu system are thought to have significant impacts on backbone topology, making this a particularly interesting use case for SINATRA Pro. Despite the challenges associated with the considerable noise inherent to the complex EF-Tu system, SINATRA Pro succeeded in identifying the meaningful topological deltas that are thought to be important for function and that were elucidated in previous studies.

### N-pocket Enlargement and *α*C Helix Displacement in Met290Ala Abl1

Protein tyrosine kinases (TKs) such as Abl1 and Src play significant roles in eukaryotic life, as phosphorylation of tyrosine residues in key proteins act as on/off switches that regulate a plethora of cellular processes and allow for efficient message passing [38]. Deregulation of the activity of these enzymes due to mutations is usually associated with severe forms of cancer and other chronic diseases, posing a grave public health problem [39]. Due to their physiological importance, the enzymatic activity of tyrosine kinases is tightly regulated by a series of structural elements that fluctuate among metastable conformations between the active and inactive states [40]. This highly dynamic behavior has been exploited for the development of TK inhibitors, such as the widely-known anticancer drug Imatinib, which exclusively targets the “DFG out” state of Abl1 [40–42]. In this conformation, the phenylalanine residue of the region known as the DFG motif (comprised of Asp381, Phe382, and Gly383) occupies Abl1’s ATP binding site, preventing substrate binding and inactivating the enzyme [43]. Other TK inhibitors such as Dasatinib are capable of binding to Abl1’s “DFG-in” conformation, in which the positions of the aspartic acid and phenylalanine side-chains are inverted with respect to their positions in DFG-out conformations, activating the enzyme (i.e., making it capable of productive phosphorylation) [42]. The transition from the DFG-in to the DFG-out state is thought to happen on the multi-millisecond timescale, which presents a challenge for capturing it with unbiased atomistic MD simulations [43]. As a workaround, previous studies have used an engineered Abl1 mutant, Met290Ala, in which the energy barrier for the DFG flip is considerably reduced, as the steric effect presented by the bulky methionine is removed [2]. Although the sampling of the entire DFG flip is a rare event that happens in the milisecond timescale and thus outside of the scope of this work, we hypothesize that the Met290Ala mutation should induce minute topological changes around the DFG motif even in shorter simulations, due to the removal of the steric hindrances associated with the bulky methionine and that SINATRA Pro is capable of detecting these changes in contrast to the wild-type dynamics. To test this hypothesis and further measure our method’s capacity for detecting localized topological changes, we ran molecular dynamics simulations on the TK domain of the unbound state of Abl1 and its Met290Ala mutant. Specifically, we run SINATRA Pro, RMSF, and the Elastic Net on the whole structure (Fig. 5(a) and Fig. S6(a)), fragments limited to residues 242-502 (Fig. 5(b) and Fig. S6(b)), and the N-lobe spanning residues 242-315 (Fig. 5(c) and Fig. S6(c)).

Since we only simulated the kinase domain of Abl1, both the N-terminal and C-terminal domains are shown to be highly dynamic as they are no longer stabilized by the mass of the entire protein. As a result, whole structural changes are overshadowed by large and noisy fluctuations, and competing methods (Elastic Net and RMSF) have a difficult time identifying enrichment in the DFG motif (Figs. S16(a) and S21(a)). Nonetheless, SINATRA Pro is able to identify the enrichment in the DFG motif regardless of the inclusion or exclusion of the N- and C-terminal (Figs. 5). The signal in the DFG motif ROI becomes better statistically resolved when we remove some of the structural noise and concentrate on regions spanning residue fragments 242-502 and 242-315 (i.e., the N-lobe). The null region test results for SINATRA Pro show that the DFG motif is indeed an important structural feature in Abl1: *P* = 8.86 × 10^*−*3^ and BF= 8.783 for the whole structure analysis (i.e., including the termini) and *P* = 2.50 × 10^*−*4^ and BF= 178 for the analysis on residues 242-502 (i.e., excluding the termini), both of which reject the null hypothesis of the ROI being identified by chance (Table 3). In these analyses, the structural differences around the DFG motif between unbound Abl1 and its Met290Ala mutant were large enough for both RMSF and the Elastic Net to have power. As a comparison, SINATRA Pro not only robustly identifies residues associated with the greater N-pocket cleft as being statistically significant (i.e., the DFG motif), but also the *α*C helix spanning residues 281-293 as a moderately enriched region (Fig. 5).

From the SINATRA Pro output, we can postulate hypotheses regarding the involvement of specific codon positions outside of the DFG motif in the concerted motions that culminate in the flip, such as the two peaks of signal surrounding it (residues 350-360 and 390-400). These interesting results show that even short simulations can prove useful for gaining mechanistic insights regarding long-timescale macromolecular relaxations, as long as the heuristics employed to analyze the resulting trajectories are capable of detecting the often minute signals associated with these topological shifts.

### Opening of Superhelix Differentiates Unbound and IBB-bound Importin-*β*

Our last analysis focused on the karyopherin Importin-*β*, an essential member of the nuclear import complex in eukaryotes, as it mediates the transportation of cargo from the cytosol to the nucleus [44]. Molecular recognition by Importin-*β* often requires the cooperative binding of molecular adaptors that recognize and bind to nuclear localization sequences (NLS)—structural motifs present in cargo destined for the nucleus [44]. Structurally, Importin-*β* is organized as a superhelix composed of up to 20 tandem HEAT repeats, each of which contain two antiparallel alpha helices linked by a turn [45]. This highly ordered structure is further stabilized by interactions with Importin-*β*-binding (IBB) domains of transport adaptors such Importin-*α* or Snurportin 1, which attach very strongly to Importin-*β* [46]. The release of IBB peptides after successful transport across the nuclear pore leads to large structural rearrangements and fluctuations that are propagated across most of Importin-*β*’s backbone [47]. Although not difficult to detect with traditional analysis pipelines, such as calculating per-residue root-mean-square fluctuations or the backbone’s radius of gyration, the pseudo-global nature of these rearrangements is diametrically opposite to most of the previously explored examples, presenting an important test for SINATRA Pro. Considering this, we ran MD simulations of unbound and IBB-bound Importin-*β* and, as with the previous examples, analyzed the resulting trajectories with SINATRA Pro to compare against standard methods. The structural features identified are projected onto the IBB-bound form (Figs. S6(d) and S18(a)). Association metrics from SINATRA Pro, RMSF, and the Elastic Net all indicate large-scale conformational changes occur upon IBB release that involve the majority of the importin-*β* structure (Figs. S18(b)-(c) and S19).

Since Importin-*β* functions as a molecular spring due to its supercoiled structure and extensive interactions with targets for transport, the sudden removal of the bound IBB domain to generate the unbound structure leads to extensive and drastic fluctuations across most of the backbone during production dynamics, originating from multiple highly-correlated nodes in each HEAT repeat. These drastic rearrangements translate to significant deltas in the topology and per-residue fluctuations that are readily detected by all tested methods. Importantly, the SINATRA Pro output replicates the expected results for the IBB bound/unbound Importin-*β* system, demonstrating the method’s capacity for picking up relevant structural determinants not only for localized changes, but also for backbone-wide large-scale fluctuations.

## Discussion

There is a growing library of computational methods that leverage geometry and topology to study aspects of both protein structure and function. For example, recent work has used shape retrieval techniques from computer vision to identify distant protein homologs [48, 49]; while others are actively studying the utility of topology-based and tessellation-based protein representations in various deep learning tasks for accurate protein structure prediction [50]. In this paper, we introduced SINATRA Pro: a topological data analytic approach designed to extract biologically-relevant structural differences between two protein ensembles. Through an extensive benchmark simulation study, we assessed the utility and statistical properties of SINATRA Pro against commonly used methods in the field. Here, we showed that our proposed framework can robustly identify both static and dynamic structural changes that occur between protein ensembles. We also highlighted that, unlike other standard approaches in the field, SINATRA Pro does not require atom-by-atom correspondences between structures and thus can be implemented using all atomic information that is available, rather than being limited to atomic features that are conserved over side-chain substitutions. With real MD data, we used SINATRA Pro to analyze five different protein systems and demonstrated its ability to identify known regions of interest that have been validated by previous experimental and computational studies, as well as reveal novel structural features that are also likely to be biologically-relevant according to evidence in the literature. Overall, these results show the promise of SINATRA Pro as a hypothesis generation tool that practitioners can use to design more informed experiments for answering downstream scientific questions (e.g., whether a mutation or chemical change “induces” a specific structural change).

There are many potential extensions to the SINATRA Pro pipeline. First, in its current form, SINATRA Pro treats all atomic features as being equally important *a priori* to the phenotype of interest. One particularly interesting extension of the method would be to up- or down-weight the contributions of different types of atomic features (e.g., carbons, hydrogens, or oxygens) or residues (e.g., serine versus arginine) to more accurately represent the topology of specific inter-atomic connections such as hydrogen and covalent bonds. In practice, this would require making such annotations and deriving topological summary statistics of protein structures based on a weighted Euler characteristic transform [51]. Another natural extension would be to apply the SINATRA Pro pipeline to other data types used to study variation in 3D protein structures such as cryogenic electron microscopy (cryo-EM), nuclear magnetic resonance (NMR) ensembles, and X-ray crystallography (i.e., electron density) data. Previous work has already shown that topological characteristics computed on tumors from magnetic resonance images (MRIs) have the potential to be powerful predictors of survival times for patients with glioblastoma multiforme (GBM) [17, 51] and other cancer subtypes [52–54]; however, it has also been noted that the efficacy of current topological summaries decreases when heterogeneity between two phenotypic classes is driven by minute differences [13]. For example, cryo-EM images can look quick similar even for two proteins harboring different mutations. SINATRA Pro’s improved ability to capture inter-class variation is driven by local fluctuations in shape morphology, so it would be interesting to see if our proposed pipeline could offer more resolved insights for these types of applications.

## URLs

SINATRA Pro software, https://github.com/lcrawlab/SINATRA-Pro; Schrödinger Desmond software, https://www.schrodinger.com/products/desmond; GROMACS software, https://www.gromacs.org; Visual Molecular Dynamics (VMD) software, https://www.ks.uiuc.edu/Research/vmd/; MDAnalysis software, https://www.mdanalysis.org; UCSF Chimera software, https://www.cgl.ucsf.edu/chimera/.

## Acknowledgements

This research was supported in part by an Alfred P. Sloan Research Fellowship and a David & Lucile Packard Fellowship for Science and Engineering awarded to LC. GM and BR were funded by National Science Foundation EPSCoR Track-II award number OIA1736253. A majority of this research was conducted using computational resources and services at the Center for Computation and Visualization (CCV), Brown University. SM would like to acknowledge partial funding from HFSP RGP005, NSF DMS 17-13012, NSF BCS 1552848, NSF DBI 1661386, NSF IIS 15-46331, NSF DMS 16-13261, as well as high-performance computing partially supported by grant 2016-IDG-1013 from the North Carolina Biotechnology Center. Any opinions, findings, and conclusions or recommendations expressed in this material are those of the author(s) and do not necessarily reflect the views of any of the funders.

## Author Contributions

WST and LC conceived the study. WST, HK, TS, SM, and LC developed the theoretical aspects of the framework. WST developed the software and carried out the statistical analyses. GM, ES, BF, and BR performed the MD simulations. GM and BR designed the strategy for the protein analysis, conducted RMSF and normal mode baseline comparisons, provided expertise about the underlying biophysics for results. WST, TS, KY, and LC conducted Elastic Net and Neural Network baseline comparisons. All authors wrote and revised the manuscript.

## Material and Methods

### Molecular Dynamics Simulations

The protein structure data used in the current study are a result of molecular dynamic (MD) simulations. For large systems (i.e., IBB-bound Importin-*β*, unbound Importin-*β*) and those containing small-molecule ligands (i.e., wild-type HIV Protease, Ile50Val HIV protease, GTP-bound EF-Tu, and GDP-bound EF-Tu), we used Schrödinger’s `Desmond` (release 2020-1) [55] to run three independent 100 nanosecond (ns) simulations for each system. This decision is rooted in Desmond’s high performance when dealing with hundreds of thousands of atoms, and the extensive validation of the small-molecule parameters contained in the OPLS3e force-field [56]. The systems were built within a dodecahedron box extending 1 nanometer (nm) beyond the solute in all three dimensions and solvated with water molecules using the SPC model [57]. Charges were neutralized by replacing a varying number of solvent molecules with sodium and chloride ions. Before production dynamics, all systems were relaxed and equilibrated with Desmond’s standard relaxation protocol, which first performs energy minimization with 50 kcal/mol/Å2 restraints on the protein’s heavy atoms, followed by an extensive equilibration protocol. This protocol is detailed below:

1. NVT equilibration at 10 K for 12 ps
2. NPT equilibration at 10 K for 12 ps
3. NPT equilibration at 300K with harmonic restraints on the protein’s heavy atoms for 120 ps
4. NPT equilibration at 300 K, unrestrained, for 240 ps,

where NVT denotes constant temperature and volume and NPT denotes constant temperature and pressure. After equilibration, unrestrained NPT production simulations were conducted at 300 K and 1 atm for 100 ns for each system, to ensure consistency among the trajectories. Time steps for all simulations were set to their default values: 2:2:6 fs (bonded:near:far).

For comparatively small systems without ligands (i.e., wild-type TEM, Arg164Ser TEM, wild-type Abl1, and Met290Val Abl1), we used `GROMACS` (release 2018-2) [58] to run three independent 100 ns simulations for each system (150 ns for Abl1). Simulations were conducted with a 2 fs time step using the Amberff14SB force-field [59] and the TIP3P water model [57]. As with the `Desmond` simulations, the systems were built within a dodecahedron box and charges neutralized by replacing a number of solvent atoms with sodium and chloride ions. For each system, energy was minimized using a steepest-descent algorithm until the maximum force on any given atom was less than 1000 kJ/mol/min. Solvent atoms were equilibrated in sequential 0.5 ns NVT and NPT simulations with solute heavy atoms restrained by a spring constant of 1,000 kJ/mol/nm^2^ using the LINCS algorithm [60]. After equilibration, production dynamics were conducted sans the position restraints. All simulations were conducted at 300 K and 1 atm. Lastly, using `Visual Molecular Dynamics (VMD)` (version 1.9.3) [61], we converted all trajectories employed in this study to a DCD file format and stripped solvent atoms to facilitate downstream computational analyses.

### Protein Structure Alignment

In the current study, protein structures are aligned in one of two ways. In the first procedure, we align protein structures by minimizing the root-mean-square distance (RMSD) between the atoms on their backbone alpha-carbons (which we denote in shorthand by C_*α*_). The first frame of the MD simulation is chosen as the reference structure. Next, all other frames (i.e., the mobile and fluctuating structures) in the dataset are aligned to this reference frame by *(i)* first superposing the center of mass of the C_*α*_ atoms to the same origin and then *(ii)* minimizing the RMSD rotation matrix. This calculation is performed using the `MDAnalysis` software package in `Python` (see Data and Software Availability) [6, 62–64]. For interclass alignment when comparing protein class *A* to class *B* (e.g., mutants versus wild-type), all frames in the trajectory of class *A* and *B* are aligned to the first frame in the trajectory of class *A*. Note that, in our current study, there is only one point mutation in each protein. In this case, determining correspondences between sequences in the trajectories of class *A* and *B* is trivial because mutated residues in the mutant structures just correspond to the un-mutated residue in the wild-type (e.g., Arg164 in WT versus Ser164 in Arg164Ser mutant in *β*-lactamase). In the controlled simulation experiments, perturbed structures were obtained by directly modifying the atomic coordinates of the pre-aligned proteins; therefore, the perturbed structures do not need further alignment since their unperturbed regions remain aligned after the controlled modifications.

In the second alignment procedure, we perform sequence-independent structural alignment where we *implicitly* normalize the 3D protein structures by rotationally aligning their topological summary statistics. To carry out this alignment procedure, we first take each pair of protein structures and superpose the center of mass of the C_*α*_ atoms to the same origin. Next, we compute topological summary statistics over the mesh representation of each structure in *m* = 500 spherically uniformly distributed directions (see the remaining sections of Material and Methods). We take the squared Euclidean distance between any two directions to be the cost needed to align structures via their topological summaries; and we determine the “optimal” direction alignment by finding the rotation that minimizes the cumulative cost of aligning all directional pairs between proteins. Here, we use the random sample consensus (RANSAC) method to determine the rotational matrix that aligns the angle between any two directions to be within an error threshold of 0.9 [65]. More specifically, we require that the dot product between two directions has to be larger than 0.9 to be considered aligned in RANSAC. Figures comparing the correlation between SINATRA Pro results on the five real protein systems using the RMSD pre-alignment versus using the topological rotational alignment can be found in the Supporting Information (Figs. S27 and S28). There are large agreements between both alignment procedures in almost all protein systems. The one exception being HIV-1 protease where we believe that the difference in results is driven by the fact that the small dynamic topology of protein makes it difficult to identify optimal stable reference signatures for alignment in summary statistic space.

### Converting Protein Structure Data to 3D Mesh Representations

To convert protein structures into a mesh representation, in the first step of the SINATRA Pro pipeline, we make use of a technique which we refer to as a “simplicial construction” (Fig. 1(b)). In this procedure, we treat the atomic positions for the protein as vertices on a 3D shape or surface. First, we draw an edge between any two atoms if their Euclidean distance is smaller than some radius cutoff *r*, namely dist | (*x*_1_, *y*_1_, *z*_1_), (*x*_2_, *y*_2_, *z*_2_)| *< r*. Next, we fill in all of the triangles (or faces) formed by these connected edges. The resulting triangulated meshes are then normalized to the unit sphere, which means that the coordinates for all atoms are scaled with respect to the mesh with the largest radius. We treat the normalized meshes as simplicial complexes which we then use to compute topological summary statistics.

### Topological Summary Statistics for Protein Mesh Representations

Adopted from its predecessor [13], the second step of the SINATRA Pro pipeline uses a tool from integral geometry and differential topology called the Euler characteristic (EC) transform [14–17]. As a brief overview of this approach, given the mesh representation ℳ of a protein structure, the Euler characteristic is an accessible topological invariant defined as

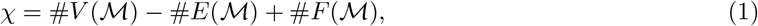

where the collection #*V* (*ℳ*), #*E*(*ℳ*), #*F* (*ℳ*) denotes the number of vertices (atoms), edges (connections between atoms), and faces (triangles enclosed by edges) of the mesh, respectively. An EC curve *χ*_*ν*_ (*ℳ*) tracks the change in the Euler characteristic with respect to a given filtration of length *l* in some direction *ν*. Theoretically, this is done by first specifying a height function *h*_*ν*_ (***x***) = ***x***^r^*ν* for some atomic position ***x*** *∈ ℳ* in direction *ν*. This height function is then used to define sublevel sets (or subparts) of the mesh 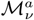 in direction *ν*, where *h*_*ν*_ (***x***) *≤ a*. In practice, the EC curve is 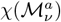 computed over a range of *l* filtration steps in direction *ν*. The corresponding EC transform is defined as the collection of EC curves across a set of *ν* = 1, …, *m* directions, and maps onto a 3D protein structure as a concatenated *J* = (*l × m*)-dimensional feature vector to be used for statistical analyses.

In previous studies, it has been observed that the Euler characteristic can be a less-than-optimal shape summary statistic when inter-class variation between 3D objects is high and driven by local fluctuations in morphology [13, 16, 17, 52, 66]. Given that this situation can be quite common in molecular dynamics, we introduce a new topological invariant which we refer to as the differential Euler characteristic (DEC) (see Fig. 1(c)). As an alternative to Eq. (1), the DEC is computed as the following

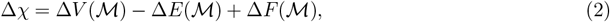

where, for some lag parameter *t*, we define Δ*V* (*ℳ*) = #*V*_*l*_(*ℳ*) *−* #*V*_*l−t*_(*ℳ*), Δ*E*(*ℳ*) = #*E*_*l*_(*ℳ*) *−* #*E*_*l−t*_(*ℳ*) and Δ*F* (*ℳ*) = #*F*_*l*_(*ℳ*) − #*F*_*l−t*_(*ℳ*). In this study, we set *t* = 1 such that, intuitively, the DEC tracks the changes (i.e., the local appearance or disappearance of topological features) in the number of vertices, edges, and faces from one sublevel set to the next. Much like with the original Euler characteristic, the DEC curve is 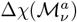 computed over a range of *l* filtration steps in a given direction *ν* and the DEC transform is similarly defined as the collection of DEC curves across a set of *ν* = 1, …, *m* directions. Overall, for each dataset with *N* total proteins, an *N × J* design matrix **X** is statistically analyzed, where the columns denote the differential Euler characteristic computed at a given filtration step and direction. Each sublevel set value, direction, and set of atomic positions used to compute a DEC curve are stored by the algorithm for the association mapping and projection phases of the pipeline.

#### Choosing the Number of Directions and Filtration Steps

In this paper, we use a series of simulations and sensitivity analyses to develop an intuition as to how to set the granularity of sublevel filtrations *l* and choose the number of directions *m* for real protein structure data (Fig. 1, Fig. S1, and Table 1). Since the structural changes that a protein class exhibits can occur on both a global and local scale, depending on its biophysical and chemical properties, we recommend choosing the former parameter *l* via cross validation or a grid-based search. For the latter, the SINATRA Pro software defines the total number of directions *m* as the union of *c* sets of cones of directions 𝒟 = ∪ 𝒞_*k*_(*θ*), where each cone 𝒞_*k*_(*θ*) = {*ν*_*k*,1_, …, *ν*_*k,d*_ | *θ*} for *k* = 1, …, *c* is parameterized by a cap radius *θ* from which equidistant vectors are generated over the unit sphere. We use cones because local shape information matters most when determining reconstructed manifolds and it has been shown that topological invariants that are measured in directions of close proximity contain similar local information [13, 15, 67, 68]. This naturally leads to the construction of sets 𝒞_*k*_(*θ*) where the angle *θ* between them is relatively small (again see Table 1). In general, we use sufficiency results for topological transforms (see Theorem 7.14 in Curry et al. [15]) to motivate the notion that considering larger numbers of *m* = *c* × *d* directions will lead to a more robust summary of 3D shapes and surfaces. Hence, ideally, one would select an effectively large number of *c* cones (and *d* directions within these cones) to ensure that SINATRA Pro is summarizing all relevant structural information about the variance between phenotypic classes (e.g., mutants versus wild-type).

### Probabilistic Model for Protein Structure Classification

In the third step of the SINATRA Pro pipeline, we use (weight-space) Gaussian process probit regression model to classify protein structures based on their topological summaries generated by the DEC transformation via Eq. (2). Here, we specify the following probabilistic hierarchical model (Fig. 1(d)) [69–73]

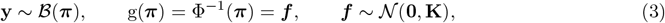

where **y** is an *N*-dimensional vector of Bernoulli distributed phenotypic class labels (e.g., mutants versus wild-type), ***π*** is an *N*-dimensional vector representing the underlying probability that a shape is classified as a “class” (e.g., *y* = 1 if “mutant”), g(**·**) is a probit link function with Φ(**·**) the cumulative distribution function (CDF) of the standard normal distribution, and ***f*** is an *N*-dimensional vector estimated from the data. We take a classic kernel regression approach [72, 74–76] where we posit that ***f*** lives within a reproducing kernel Hilbert space (RKHS) defined by some (nonlinear) covariance function, which implicitly accounts for higher-order interactions between features, leading to more complete classifications of structural data [77–79]. To this end, we assume ***f*** is normally distributed with mean vector **0** and covariance matrix **K** with elements defined by the radial basis function **K**_*ii*′_ = exp{−*ϑ*‖**x**_*i*_ *−* **x**_*i*′_‖^2^} with bandwidth *ϑ* set using the “median criterion” approach to maintain numerical stability and avoid additional computational costs [80]. Here, **x**_*i*_ and **x**_*i*_*I* denote the collection of topological features for the *i*-th and *i*^*/*^-th observation in **X**, respectively. The full model specified in Equation (3) is commonly referred to as “Gaussian process classification” (GPC).

Given the complete specification of the GPC, we use Bayesian inference to draw samples from the posterior distribution of the latent variables, which is proportional to *p*(***f*** | **y**) ∝ *p*(**y** | ***f***) × *p*(***f***). Here, *p*(**y** | ***f***) denotes the likelihood of the observed binary labels given the functions (i.e., the Bernoulli distribution), and *p*(***f***) is the prior distribution for the latent variables (i.e., the multivariate normal distribution). The probit likelihood in Eq. (3) makes it intractable to estimate the posterior distribution *p*(***f*** | **y**) via a closed-form solution. We instead use a Markov chain Monte Carlo (MCMC) method called “elliptical slice sampling” to conduct posterior inference (see Data and Software Dependencies) [81].

### Feature Selection of Topological Summary Statistics

After implementing the elliptical slice sampling algorithm to estimate the posterior distribution of the latent variables ***f*** in Eq. (3), we define a nonparametric effect size for each topological summary statistic via the following standard projection [78, 82]

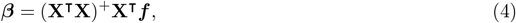

where **M**^+^ is used to denote the generalized inverse of a matrix **M**, and each element in ***β*** details the nonlinear relationship between the DEC topological summary statistics and the variance between protein structures. In order to determine a statistical rank ordering for these effect sizes, we assign an information theoretic-based measure of relative centrality to each *j*-th topological feature using Kullback-Leibler divergence (KLD) [79]

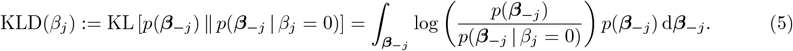

for *j* = 1, …, *J* topological features. Finally, we normalize to obtain an association metric (Fig. 1(d)),

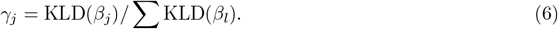

There are two key takeaways from this scaled formulation. First, the KLD is non-negative, and it equals zero if and only if the posterior distribution of ***β***_*−j*_ is independent of the effect *β*_*j*_. Intuitively, this is equivalent to saying that removing an unimportant topological feature has no impact on explaining the variance between different protein structure. Second, ***γ*** = (*γ*_1_, …, *γ*_*J*_) is bounded on the unit interval [0, 1] with the natural interpretation of providing relative evidence of association for each DEC statistic (where values close to 1 suggest greater importance). From a classical hypothesis testing point-of-view, the null hypothesis for Eq. (6) assumes that every DEC feature equally contribute to the total variance between proteins, while the alternative hypothesis proposes that some DEC features are better associated with biophysical changes in protein structures than others [13, 79].

#### Closed Form Solution for Atomic-Level Association Measures

For simplicity, we assume that the implied posterior distribution of ***β*** (deterministically given in Eq. (4)) is approximately multivariate normal with an empirical mean vector ***µ*** and positive semi-definite covariance/precision matrix **Σ** = **Λ**^*−*1^ [13, 79]. Given these values, we iteratively partition such that, for each *j*-th topological feature:

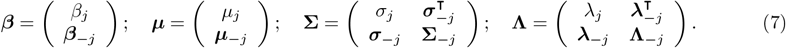

Under normality assumptions, Eq. (5) has the following closed form solution

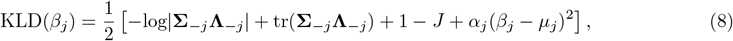

where log| **·** | represents the matrix log-determinant function, and tr(**·**) is the matrix trace function. Importantly, the term 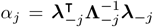 characterizes the linear (and non-negative) rate of change of information when the effect of any topological feature is absent from the analysis [79]. By symmetry in the notation for elements of the sub-vectors and sub-matrices, we simply permute the order of the variables in ***β*** and iteratively compute the KLD to measure the centrality of each DEC transform.

#### Approximate Computation

In practice, we use a few approximations to scale the otherwise computationally expensive steps in Eq. (8). The first approximation involves computing the log determinant. With a dataset of reasonably dense meshes, the number of topological features is expected to be large (i.e., *J* ≫ 0). In this setting, the term −log(|**Σ**_*−j*_**Λ**_*−j*_|) + tr(**Σ**_*−j*_**Λ**_*−j*_) + (1 − *p*) remains relatively equal for each feature *j* and makes a negligible contribution to the entire sum. Thus, we simplify Eq. (8) to

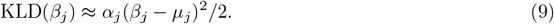

This approximation of the KLD still relies on the full precision matrix **Λ**. For a large number of topological features *J*, this calculation is expensive; however, it is only done once and and can be done with efficient matrix decomposition. The rate of change parameter 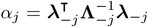, on the other hand, depends onthe partitioned matrix 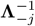 for every *j*-th topological feature. This requires inverting a (*J* − 1) × (*J* − 1) matrix *J* times. Fortunately, we can reduce this computational burden by taking advantage of the fact that any 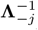 is formed by removing the *j*-th row and column from the precision matrix **Λ**. Therefore, given the partition in Eq. (7), we can use the Sherman-Morrison formula [83] to efficiently approximate these quantities using the following rank-1 update for each topological feature

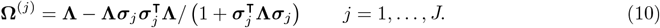

Here, ***σ***_*j*_ is the *j*-th column from the posterior covariance matrix **Σ**, and each 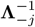 is approximated by removing the *j*-th row and column from **Ω**^(*j*)^. Ultimately, this reduces the computational complexity of Equation (9) to just *J*-independent *O*(*J*^2^) operations which can be parallelized.

### Reconstruction and Visualization of Biophysical Signatures

After obtaining association measures ***γ*** for each topological feature computed via Eq. (6), in the fourth step of the SINATRA Pro pipeline, we map this information back onto the original structures to visualize topological differences between the protein classes. The main idea is that we want to select or prioritize atoms that correspond to the topological features with the greatest association measures. To do this, we perform a criterion-based *reconstruction* algorithm [13]. In each direction, each atom (i.e., vertex) lies along a filtration step that corresponds to a *γ*_*j*_ value. Therefore, each atom corresponds to *m* = (*c × d*) values in ***γ***. To perform the reconstruction, we sort the values in ***γ*** from smallest to largest and continuously increase a threshold. If all of the *γ* values corresponding to an atom are larger than the threshold, the atom is considered “alive”. As the threshold is increased, when the criterion is no longer satisfied, the atom is considered “dead” and that minimum value below the threshold (which we will denote by 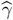) is assigned the atom as its evidence score. This calculation is repeated for each frame in the dataset. For atomic-level evidence scores (e.g., Figs. S5 and S6), the 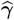 values are ranked among all atoms and scaled from 0 (lowest) to 100 (highest) to facilitate the visualization and interpretation of structural and biophysical enrichment. To compute residue-level evidence scores (e.g., Fig. 3-5, and Figs. S7, S9, S10, S12, S13, S15, S17, and S18), we take the average of the 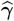 values for all atoms within a residue which are then also ranked and scaled from 0 to 100.

### Performance Assessment for Controlled Simulation Study

We demonstrate the power of the SINATRA Pro pipeline for identifying biophysical signatures in protein dynamics via multiple controlled simulations studies using the sequential procedure:

1. Fit the Gaussian process classification (GPC) model using elliptical slice sampling and compute relative centrality association measures *γ*_*j*_ for each *j*-th topological feature (i.e., differential Euler characteristic or DEC per sublevel set filtration). Recall, the total number of features *J* = *c × d × l* is a product of *(i) c*, the number of cones of directions; *(ii) d*, the number of directions within each cone; and *(iii) l*, the number of sublevel sets (i.e., steps in the filtration) used to compute the DEC along a given direction.
2. Sort the topological features from largest to smallest according to their association measures *γ*_1_ ≥ *γ*_2_ ≥ ⋯ ≥ *γ*_*p*_.
3. By iteratively moving through the sorted measures *T*_*k*_ = *γ*_*k*_ (starting with *k* = 1), we reconstruct the atoms corresponding to the topological features with {*j* : *γ*_*j*_ *≥ T*_*k*_}.

An atom is “detected” when the sublevel set in which it resides is selected across all of the directions within a particular cone. We form a union of the set of detected atoms across all cones to construct the set of reconstructed vertices at a given level *T*_*k*_. Using this set of vertices, we compute the true positive rate (TPR) and false positive rate (FPR) by assessing overlap with the set of truly associated (i.e., perturbed) atoms used to generate the protein classes:

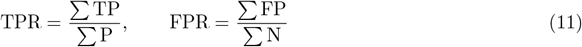

where TP is the number of correctly detected true atoms, P is the total number of causal atoms, TN stands for the true negatives detected by the SINATRA Pro pipeline, and N stands for the total number of non-causal atoms. In this manner, we obtain a receiver operating characteristic (ROC) curve for the simulation studies (see Fig. 2 and Fig. S1).

### ROI Null Experiment and Statistical Assessment

To statistically assess whether SINATRA Pro is identifying the known regions of interest (ROI) in proteins by chance (see Table 2), we use a previously developed null-based scoring method [13]. The goal of this analysis is to estimate the probability of obtaining a result from SINATRA Pro under the assumption that the null hypothesis *H*_0_ of there being no structural differences between mutant and wild-type proteins is true. Here, we treat the *K* atoms located within each ROI of every mutant protein as a landmark. We construct a test statistic *τ*^*∗*^ for each ROI by summing the association metric scores of every atom it contains. To construct a “null” distribution and assess the strength of any score *τ*^*∗*^, we randomly select *T* “seed” atoms across the mesh outside the ROI for each mutant protein and uniformly generate *T*-”null” regions that are also *K*-atoms wide. We then compute similar (null) scores *τ*_1_, …, *τ*_*T*_ for each randomly generated region. A “*P*-value”-like quantity (for the *i*-th mutant protein) is then generated by:

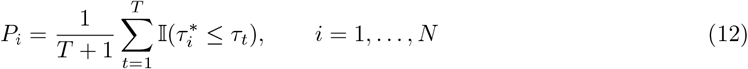

where 𝕀(**·**) is an indicator function, and a smaller *P*_*i*_ means more confidence in either method’s ability to find the desired paraconid landmark. To ensure the robustness of this analysis, we generate the *N*-random null regions using a *K*-nearest neighbors (KNN) algorithm on each of the *T*-random seed vertices [84]. We also use a calibration formula to transform each *P*-value to an approximate Bayes factor (BF) [85], which is defined as the ratio of the marginal likelihood under the alternative hypothesis *H*_1_ (i.e., that there is indeed a structural difference between phenotypic classes) versus the null hypothesis *H*_0_:

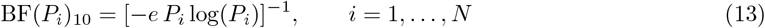

for *P*_*i*_ *<* 1*/e* and BF(*P*_*i*_)_10_ is an estimate of Pr(*H*_1_ | *M*)*/*Pr(*H*_0_ | *M*), where *M* is again used to denote the protein meshes. We take the median of the *P*_*i*_ and *BF*(*P*_*i*_)_10_ values in Eqs. (12) and (13) across all mutant proteins, respectively, and report them in Table 3.

## Data and Software Dependencies

Code for implementing the SINATRA Pro pipeline is freely available at https://github.com/lcrawlab/SINATRA-Pro, and is written in `Python` (version 3.6.9). As part of this procedure: *(i)* inference for the Gaussian process classification (GPC) model is based on an elliptical slice sampling algorithm adapted from the `R` package `FastGP` (version 1.2) [86] and *(ii)* the computation of nonlinear effect sizes and association measures for the differentiated Euler characteristic (DEC) curves was done by adapting the “RelATive cEntrality (RATE)” source code originally written in `R` (version 1.0.0; https://github.com/lorinanthony/RATE) [79]. Visualizing the reconstructed protein regions outputted by SINATRA Pro was done using the extensive molecular modeling system software `Chimera` (version 1.14) [87]. Molecular dynamic simulations were performed using Schrödinger’s `Desmond` (release 2020-1) [55] and `GROMACS` (release 2018-2) [58]. Furthermore, preprocessing steps for the protein structures resulting from MD simulations examined in the study were performed using `Visual Molecular Dynamics (VMD)` (version 1.9.3) [61] and the `Python` library `MDAnalysis` (version 1.1.1) [6,62–64]. Data generated from the MD simulations can be downloaded at https://www.dropbox.com/sh/l4fj3paagyrpu2f/AAA65_NbNaX5IUllrazScZo9a?dl=0. Scripts to reproduce the results in this paper are also publicly available and can be found at https://github.com/lcrawlab/SINATRA_Pro_Paper_Results.

## References

1. Orengo CA, Todd AE, Thornton JM. From protein structure to function. Current Opinion in Structural Biology. 1999;9(3):374–382. Available from: https://www.sciencedirect.com/science/article/pii/S0959440X99800517.

2. Shan Y, Seeliger MA, Eastwood MP, Frank F, Xu H, Jensen MØ, et al. A conserved protonation-dependent switch controls drug binding in the Abl kinase. Proceedings of the National Academy of Sciences of the United States of America. 2009;106(1):139–144. Available from: https://www.ncbi.nlm.nih.gov/pmc/articles/PMC2610013/.

3. Wilson C, Agafonov RV, Hoemberger M, Kutter S, Zorba A, Halpin J, et al. Using ancient protein kinases to unravel a modern cancer drug’s mechanism. Science. 2015;347(6224):882–886. Available from: https://science.sciencemag.org/content/347/6224/882.

4. Hollingsworth SA, Dror RO. Molecular Dynamics Simulation for All. Neuron. 2018;99(6):1129–1143. Publisher: Elsevier. Available from: https://www.cell.com/neuron/abstract/S0896-6273(18)30684-6.

5. Grossfield A, Patrone PN, Roe DR, Schultz AJ, Siderius DW, Zuckerman DM. Best Practices for Quantification of Uncertainty and Sampling Quality in Molecular Simulations [Article v1.0]. Living journal of computational molecular science. 2018;1(1):5067. Available from: https://www.ncbi.nlm.nih.gov/pmc/articles/PMC6286151/.

6. Michaud-Agrawal N, Denning EJ, Woolf TB, Beckstein O. MDAnalysis: A toolkit for the analysis of molecular dynamics simulations. Journal of Computational Chemistry. 2011;32(10):2319–2327.

7. Grant BJ, Skjaerven L, Yao XQ. The Bio3D packages for structural bioinformatics. Protein Science: A Publication of the Protein Society. 2021;30(1):20–30.

8. Maisuradze GG, Liwo A, Scheraga HA. Principal Component Analysis for Protein Folding Dynamics. Journal of Molecular Biology. 2009;385(1):312–329. Available from: https://www.sciencedirect.com/science/article/pii/S0022283608012886.

9. Sittel F, Jain A, Stock G. Principal component analysis of molecular dynamics: On the use of Cartesian vs. internal coordinates. The Journal of Chemical Physics. 2014;141(1):014111. Publisher: American Institute of Physics. Available from: https://aip.scitation.org/doi/10.1063/1.4885338.

10. Pál C, Papp B, Lercher MJ. An integrated view of protein evolution. Nature Reviews Genetics. 2006;7(5):337–348. Bandiera abtest: a Cg type: Nature Research Journals Number: 5 Primary atype: Reviews Publisher: Nature Publishing Group. Available from: https://www.nature.com/articles/nrg1838.

11. Rustamov RM, Ovsjanikov M, Azencot O, Ben-Chen M, Chazal F, Guibas L. Map-based exploration of intrinsic shape differences and variability. ACM Trans Graph. 2013;32(4):1–12.

12. Huang R, Achlioptas P, Guibas L, Ovsjanikov M. Limit Shapes–A Tool for Understanding Shape Differences and Variability in 3D Model Collections. Comput Graph Forum. 2019;38(5):187–202.

13. Wang B, Sudijono T, Kirveslahti H, Gao T, Boyer DM, Mukherjee S, et al. A Statistical Pipeline for Identifying Physical Features that Differentiate Classes of 3D Shapes. Ann Appl Stat. 2021;15(2):638–661.

14. Turner K, Mukherjee S, Boyer DM. Persistent homology transform for modeling shapes and surfaces. Inf Inference. 2014;3(4):310–344. Available from: https://academic.oup.com/imaiai/article-abstract/3/4/310/724811?redirectedFrom=fulltext.

15. Curry J, Mukherjee S, Turner K. How many directions determine a shape and other sufficiency results for two topological transforms. arXiv. 2019;p. 1805.09782. Available from: https://arxiv.org/abs/1805.09782.

16. Ghrist R, Levanger R, Mai H. Persistent homology and Euler integral transforms. J Appl and Comput Topology. 2018;2(1-2):55–60. Available from: https://link.springer.com/article/10.1007/s41468-018-0017-1.

17. Crawford L, Monod A, Chen AX, Mukherjee S, Rabadán A. Predicting clinical outcomes in glioblastoma: an application of topological and functional data analysis. J Am Stat Assoc. 2020;115(531):1139–1150.

18. Pedregosa F, Varoquaux G, Gramfort A, Michel V, Thirion B, Grisel O, et al. Scikit-learn: Machine Learning in Python. Journal of Machine Learning Research. 2011;12:2825–2830.

19. Xu B, Wang N, Chen T, Li M. Empirical evaluation of rectified activations in convolutional network; 2015. ArXiv.

20. Simonyan K, Vedaldi A, Zisserman A. Deep inside convolutional networks: Visualising image classification models and saliency maps. arXiv preprint 13126034. 2013;.

21. Stojanoski V, Chow DC, Hu L, Sankaran B, Gilbert HF, Prasad BVV, et al. A triple mutant in the ?-loop of TEM-1 β-lactamase changes the substrate profile via a large conformational change and an altered general base for catalysis. Journal of Biological Chemistry. 2015;290(16):10382–10394. Available from: https://pubmed.ncbi.nlm.nih.gov/25713062.

22. Egorov A, Rubtsova M, Grigorenko V, Uporov I, Veselovsky A. The Role of the ?-Loop in Regulation of the Catalytic Activity of TEM-Type β-Lactamases. Biomolecules. 2019;9(12). Available from: https://www.ncbi.nlm.nih.gov/pmc/articles/PMC6995641/.

23. Knox JR. Extended-spectrum and inhibitor-resistant TEM-type beta-lactamases: mutations, specificity, and three-dimensional structure. Antimicrobial Agents and Chemotherapy. 1995;39(12):2593–2601. Available from: https://www.ncbi.nlm.nih.gov/pmc/articles/PMC162995/.

24. Gniadkowski M. Evolution of extended-spectrum β-lactamases by mutation. Clinical Microbiology and Infection. 2008;14:11–32. Available from: https://www.sciencedirect.com/science/article/pii/S1198743X14604729.

25. Brik A, Wong CH. HIV-1 protease: mechanism and drug discovery. Organic & Biomolecular Chemistry. 2003;1(1):5–14. Publisher: Royal Society of Chemistry. Available from: https://pubs.rsc.org/en/content/articlelanding/2003/ob/b208248a.

26. Karacostas V, Wolffe EJ, Nagashima K, Gonda MA, Moss B. Overexpression of the HIV-1 gag-pol polyprotein results in intracellular activation of HIV-1 protease and inhibition of assembly and budding of virus-like particles. Virology. 1993;193(2):661–671.

27. Lv Z, Chu Y, Wang Y. HIV protease inhibitors: a review of molecular selectivity and toxicity. HIV/AIDS (Auckland, NZ). 2015;7:95–104. Available from: https://www.ncbi.nlm.nih.gov/pmc/articles/PMC4396582/.

28. Rhee SY, Taylor J, Fessel WJ, Kaufman D, Towner W, Troia P, et al. HIV-1 protease mutations and protease inhibitor cross-resistance. Antimicrobial Agents and Chemotherapy. 2010;54(10):4253–4261.

29. Sheik Amamuddy O, Bishop NT, Tastan Bishop Ö. Characterizing early drug resistance-related events using geometric ensembles from HIV protease dynamics. Scientific Reports. 2018;8(1):17938. Number: 1 Publisher: Nature Publishing Group. Available from: https://www.nature.com/articles/s41598-018-36041-8.

30. Palese LL. Conformations of the HIV-1 protease: A crystal structure data set analysis. Biochimica Et Biophysica Acta Proteins and Proteomics. 2017;1865(11 Pt A):1416–1422.

31. Hornak V, Okur A, Rizzo RC, Simmerling C. HIV-1 protease flaps spontaneously open and reclose in molecular dynamics simulations. Proceedings of the National Academy of Sciences of the United States of America. 2006;103(4):915–920. Available from: http://www.pnas.org/content/103/4/915.abstract.

32. Liu F, Kovalevsky AY, Tie Y, Ghosh AK, Harrison RW, Weber IT. Effect of flap mutations on structure of HIV-1 protease and inhibition by saquinavir and darunavir. Journal of Molecular Biology. 2008;381(1):102–115. Available from: https://pubmed.ncbi.nlm.nih.gov/18597780.

33. Adkins JC, Faulds D. Amprenavir. Drugs. 1998 Jun;55(6):837–842. Available from: https://doi.org/10.2165/00003495-199855060-00015.

34. Harvey KL, Jarocki VM, Charles IG, Djordjevic SP. The Diverse Functional Roles of Elongation Factor Tu (EF-Tu) in Microbial Pathogenesis. Frontiers in Microbiology. 2019;10. Publisher: Frontiers. Available from: https://www.frontiersin.org/articles/10.3389/fmicb.2019.02351/full.

35. Warias M, Grubmüller H, Bock LV. tRNA Dissociation from EF-Tu after GTP Hydrolysis: Primary Steps and Antibiotic Inhibition. Biophysical Journal. 2020;118(1):151–161. Available from: https://www.sciencedirect.com/science/article/pii/S000634951930877X.

36. Schmeing TM, Voorhees RM, Kelley AC, Gao YG, Murphy FV, Weir JR, et al. The Crystal Structure of the Ribosome Bound to EF-Tu and Aminoacyl-tRNA. Science. 2009;326(5953):688–694. Publisher: American Association for the Advancement of Science Section: Research Article. Available from: https://science.sciencemag.org/content/326/5953/688.

37. Li H, Yao XQ, Grant BJ. Comparative structural dynamic analysis of GTPases. PLOS Computational Biology. 2018;14(11):e1006364. Publisher: Public Library of Science. Available from: https://journals.plos.org/ploscompbiol/article?id=10.1371/journal.pcbi.1006364.

38. Hubbard SR, Till JH. Protein tyrosine kinase structure and function. Annual Review of Biochemistry. 2000;69:373–398.

39. Greuber EK, Smith-Pearson P, Wang J, Pendergast AM. Role of ABL Family Kinases in Cancer: from Leukemia to Solid Tumors. Nature Reviews Cancer. 2013;13(8):559–571. Available from: https://www.ncbi.nlm.nih.gov/pmc/articles/PMC3935732/.

40. Reddy EP, Aggarwal AK. The Ins and Outs of Bcr-Abl Inhibition. Genes & Cancer. 2012;3(5-6):447–454. Available from: https://www.ncbi.nlm.nih.gov/pmc/articles/PMC3513788/.

41. Sacha T. Imatinib in Chronic Myeloid Leukemia: an Overview. Mediterranean Journal of Hematology and Infectious Diseases. 2014;6(1):e2014007. Available from: https://www.ncbi.nlm.nih.gov/pmc/articles/PMC3894842/.

42. Aguilera DG, Tsimberidou AM. Dasatinib in chronic myeloid leukemia: a review. Therapeutics and Clinical Risk Management. 2009;5:281–289. Available from: https://www.ncbi.nlm.nih.gov/pmc/articles/PMC2697539/.

43. Xie T, Saleh T, Rossi P, Kalodimos CG. Conformational states dynamically populated by a kinase determine its function. Science. 2020;370(6513):eabc2754. Publisher: American Association for the Advancement of Science Section: Research Article. Available from: https://science.sciencemag.org/content/early/2020/09/30/science.abc2754.

44. Harel A, Forbes DJ. Importin Beta: Conducting a Much Larger Cellular Symphony. Molecular Cell. 2004;16(3):319–330. Publisher: Elsevier. Available from: https://www.cell.com/molecular-cell/abstract/S1097-2765(04)00647-1.

45. Zachariae U, Grubmüller H. Importin-β: Structural and Dynamic Determinants of a Molecular Spring. Structure. 2008;16(6):906–915. Available from: https://www.sciencedirect.com/science/article/pii/S0969212608001445.

46. Cingolani G, Petosa C, Weis K, Müller CW. Structure of importin-β bound to the IBB domain of importin-α. Nature. 1999;399(6733):221–229. Number: 6733 Publisher: Nature Publishing Group. Available from: https://www.nature.com/articles/20367.

47. Halder K, Dölker N, Van Q, Gregor I, Dickmanns A, Baade I, et al. MD Simulations and FRET Reveal an Environment-Sensitive Conformational Plasticity of Importin-β. Biophysical Journal. 2015;109(2):277–286. Available from: https://www.ncbi.nlm.nih.gov/pmc/articles/PMC4621615/.

48. Langenfeld F, Peng Y, Lai YK, Rosin PL, Aderinwale T, Terashi G, et al. SHREC 2020: Multidomain protein shape retrieval challenge. Computers & Graphics. 2020;91:189–198. Available from: https://www.sciencedirect.com/science/article/pii/S0097849320301151.

49. Machat M, Langenfeld F, Craciun D, Sirugue L, Labib T, Lagarde N, et al. Comparative evaluation of shape retrieval methods on macromolecular surfaces: an application of computer vision methods in structural bioinformatics. Bioinformatics. 2021 07;Btab511. Available from: https://doi.org/10.1093/bioinformatics/btab511.

50. Kinch LN, Pei J, Kryshtafovych A, Schaeffer RD, Grishin NV. Topology evaluation of models for difficult targets in the 14th round of the critical assessment of protein structure prediction (CASP14). Proteins. 2021;89(12):1673–1686.

51. Jiang Q, Kurtek S, Needham T. The Weighted Euler Curve Transform for Shape and Image Analysis. CoRR. 2020;abs/2004.11128. Available from: https://arxiv.org/abs/2004.11128.

52. Moon C, Li Q, Xiao G. Predicting survival outcomes using topological features of tumor pathology images. arXiv. 2020;p. 2012.12102.

53. Somasundaram E, Litzler A, Wadhwa R, Owen S, Scott J. Persistent homology of tumor CT scans is associated with survival in lung cancer. Medical Physics. 2021;48(11):7043–7051.

54. Vipond O, Bull JA, Macklin PS, Tillmann U, Pugh CW, Byrne HM, et al. Multiparameter persistent homology landscapes identify immune cell spatial patterns in tumors. Proc Natl Acad Sci U S A. 2021;118(41).

55. SC ‘06: Proceedings of the 2006 ACM/IEEE Conference on Supercomputing. New York, NY, USA: Association for Computing Machinery; 2006.

56. Harder E, Damm W, Maple J, Wu C, Reboul M, Xiang JY, et al. OPLS3: A Force Field Providing Broad Coverage of Drug-like Small Molecules and Proteins. Journal of Chemical Theory and Computation. 2016;12(1):281–296. Publisher: American Chemical Society. Available from: https://doi.org/10.1021/acs.jctc.5b00864.

57. Mark P, Nilsson L. Structure and Dynamics of the TIP3P, SPC, and SPC/E Water Models at 298 K. The Journal of Physical Chemistry A. 2001;105(43):9954–9960. Available from: https://doi.org/10.1021/jp003020w.

58. Van Der Spoel D, Lindahl E, Hess B, Groenhof G, Mark AE, Berendsen HJC. GROMACS: Fast, flexible, and free. Journal of Computational Chemistry. 2005;26(16):1701–1718.

59. Maier JA, Martinez C, Kasavajhala K, Wickstrom L, Hauser KE, Simmerling C. ff14SB: Improving the Accuracy of Protein Side Chain and Backbone Parameters from ff99SB. Journal of Chemical Theory and Computation. 2015;11(8):3696–3713. Publisher: American Chemical Society. Available from: https://doi.org/10.1021/acs.jctc.5b00255.

60. Hess B, Bekker H, Berendsen HJC, Fraaije JGEM. LINCS: A linear constraint solver for molecular simulations. Journal of Computational Chemistry. 1997;18(12):1463–1472.

61. Humphrey W, Dalke A, Schulten K. VMD – Visual Molecular Dynamics. Journal of Molecular Graphics. 1996;14:33–38.

62. Theobald DL. Rapid calculation of RMSDs using a quaternion-based characteristic polynomial. Acta Crystallographica Section A. 2005;61(4):478–480.

63. Liu P, Agrafiotis DK, Theobald DL. Fast determination of the optimal rotational matrix for macromolecular superpositions. Journal of Computational Chemistry. 2010;31(7):1561–1563. PMC2958452[pmcid].

64. Gowers RJ, Linke M, Barnoud J, Reddy TJE, Melo MN, Seyler SL, et al. MDAnalysis: A Python Package for the Rapid Analysis of Molecular Dynamics Simulations. In: Proceedings of the 15th Python in Science Conference; 2016. p. 98–105.

65. Fischler MA, Bolles RC. Random Sample Consensus: A Paradigm for Model Fitting with Applications to Image Analysis and Automated Cartography. Commun ACM. 1981;24(6):381–395. Available from: https://doi.org/10.1145/358669.358692.

66. Jiang Q, Kurtek S, Needham T. The Weighted Euler Curve Transform for Shape and Image Analysis. In: Proceedings of the IEEE/CVF Conference on Computer Vision and Pattern Recognition (CVPR) Workshops; 2020..

67. Fasy BT, Micka S, Millman DL, Schenfisch A, Williams L. Challenges in reconstructing shapes from Euler characteristic curves. arXiv. 2018;p. 1811.11337.

68. Oudot S, Solomon E. Inverse Problems in Topological Persistence. In: Baas NA, Carlsson GE, Quick G, Szymik M, Thaule M, editors. Topological Data Analysis. Cham: Springer International Publishing; 2020. p. 405–433.

69. Neal RM. Monte Carlo implementation of Gaussian process models for Bayesian regression and-Monte Carlo implementation of Gaussian process models for Bayesian regression and classification. Dept. of Statistics, University of Toronto; 1997. 9702.

70. Neal RM. Regression and classification using Gaussian process priors. Bayesian Anal. 1998;6:475.

71. Williams CKI, Barber D. Bayesian classification with Gaussian processes. IEEE Trans Pattern Anal Mach Intell. 1998;20(12):1342–1351. Available from: https://ieeexplore.ieee.org/document/735807/.

72. Rasmussen CE, Williams CKI. Gaussian processes for machine learning. Cambridge, MA: MIT Press; 2006.

73. Nickisch H, Rasmussen CE. Approximations for binary Gaussian process classification. J Mach Learn Res. 2008;9(10):2035–2078.

74. Schölkopf B, Herbrich R, Smola AJ. A generalized representer theorem. In: Proceedings of the 14th Annual Conference on Computational Learning Theory and and 5th European Conference on Computational Learning Theory. London, UK, UK: Springer-Verlag; 2001. p. 416–426. Available from: http://dl.acm.org/citation.cfm?id=648300.755324.

75. Pillai NS, Wu Q, Liang F, Mukherjee S, Wolpert R. Characterizing the function space for Bayesian kernel models. J Mach Learn Res. 2007;8:1769–1797.

76. Zhang Z, Dai G, Jordan MI. Bayesian generalized kernel mixed models. J Mach Learn Res. 2011;12:111–139.

77. Jiang Y, Reif JC. Modeling epistasis in genomic selection. Genetics. 2015;201:759–768.

78. Crawford L, Wood KC, Zhou X, Mukherjee S. Bayesian approximate kernel regression with variable selection. J Am Stat Assoc. 2018;113(524):1710–1721. Available from: https://doi.org/10.1080/01621459.2017.1361830.

79. Crawford L, Flaxman SR, Runcie DE, West M. Variable prioritization in nonlinear black box methods: a genetic association case study. Ann Appl Stat. 2019;13(2):958–989. Available from: https://projecteuclid.org/euclid.aoas/1560758434.

80. Chaudhuri A, Kakde D, Sadek C, Gonzalez L, Kong S. The mean and median criteria for kernel bandwidth selection for support vector data description. Data Mining Workshops (ICDMW), 2017 IEEE International Conference on. 2017;p. 842–849. Available from: https://ieeexplore.ieee.org/abstract/document/8215749/.

81. Murray I, Prescott Adams R, MacKay DJ. Elliptical slice sampling. Proceedings of the Thirteenth International Conference on Artificial Intelligence and Statistics. 2010;p. 541–548.

82. Singleton KR, Crawford L, Tsui E, Manchester HE, Maertens O, Liu X, et al. Melanoma therapeutic strategies that select against resistance by exploiting MYC-driven evolutionary convergence. Cell Rep. 2017;21(10):2796–2812.

83. Hager WW. Updating the inverse of a matrix. SIAM Review. 1989;31(2):221–239.

84. Cover T, Hart P. Nearest neighbor pattern classification. IEEE Trans Inf Theor. 2006;13(1):21–27. Available from: https://doi.org/10.1109/TIT.1967.1053964.

85. Sellke T, Bayarri MJ, Berger JO. Calibration of p-values for testing precise null hypotheses. Am Stat. 2001;55(1):62–71.

86. Gopalan G, Bornn L. FastGP: An R package for Gaussian processes. arXiv. 2015;p. 1507.06055. Available from: https://arxiv.org/abs/1507.06055.

87. Pettersen EF, Goddard TD, Huang CC, Couch GS, Greenblatt DM, Meng EC, et al. UCSF Chimera–a visualization system for exploratory research and analysis. J Comput Chem. 2004;25(13):1605–1612.

88. Belongie S. Rodrigues’ rotation formula. From MathWorld–A Wolfram Web Resource, created by Eric W Weisstein http://mathworld wolfram com/RodriguesRotationFormula html. 1999;.

89. Wallin G, Kamerlin SCL, Åqvist J. Energetics of activation of GTP hydrolysis on the ribosome. Nature Communications. 2013;4(1):1733. Available from: https://doi.org/10.1038/ncomms2741.

90. Mondal D, Warshel A. EF-Tu and EF-G are activated by allosteric effects. Proceedings of the National Academy of Sciences of the United States of America. 2018;115(13):3386. Available from: http://www.pnas.org/content/115/13/3386.abstract.

91. Kornev AP, Haste NM, Taylor SS, Eyck LFT. Surface comparison of active and inactive protein kinases identifies a conserved activation mechanism. Proc Natl Acad Sci U S A. 2006;103(47):17783–17788.

92. Azam M, Seeliger MA, Gray NS, Kuriyan J, Daley GQ. Activation of tyrosine kinases by mutation of the gatekeeper threonine. Nat Struct Mol Biol. 2008;15(10):1109–1118.

93. Kornev AP, Taylor SS. Defining the conserved internal architecture of a protein kinase. Biochim Biophys Acta. 2010;1804(3):440–444.

94. Berman HM, Westbrook J, Feng Z, Gilliland G, Bhat TN, Weissig H, et al. The Protein Data Bank. Nucleic Acids Research. 2000;28(1):235–242. Available from: https://doi.org/10.1093/nar/28.1.235.95.463540v2

