## Supplementary Figures and Tables for "A Topological Data Analytic Approach for Discovering Biophysical Signatures in Protein Dynamics"

### 1 Supplementary Figures

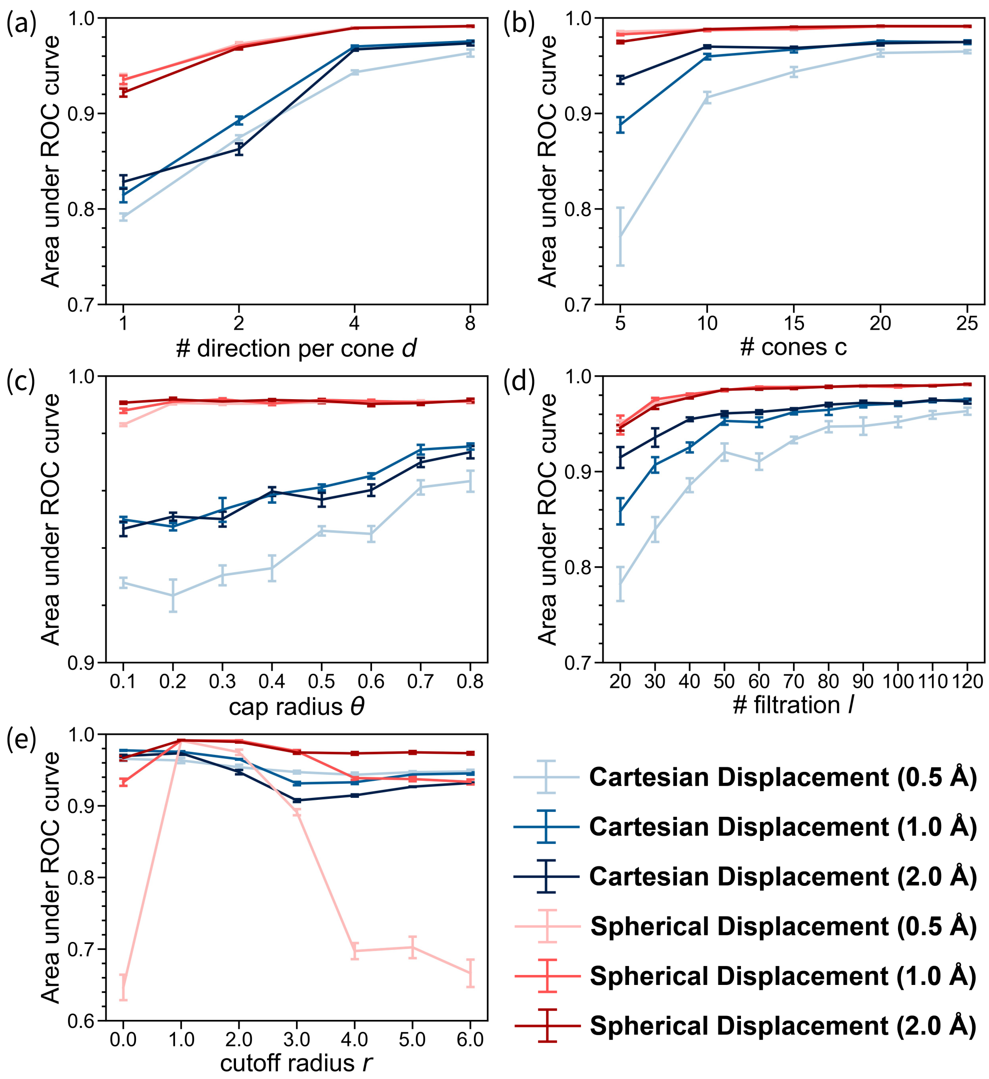

**Figure S1. Power and sensitivity analysis assessing the robustness of SINATRA Pro to different free parameter settings in controlled molecular dynamic (MD) simulations.** To generate data for these simulations, we consider two phenotypic classes using real structural data of wild-type $\beta$-lactamase (TEM). In the first phenotypic class, structural protein data are drawn from equally spaced intervals over a 100 ns MD trajectory (e.g., $t_{\text{MD}} = [0, 1, 2, 3, \ldots, 99]$ ns + $\delta$, where $\delta$ is a time offset parameter). In the second phenotypic class, proteins are drawn from 0.5 ns intervals later relative to the first group (e.g., $t_{\text{MD}} = [0.5, 1.5, 2.5, 3.5, \ldots, 99.5]$ ns + $\delta$) to introduce physical thermal noise, and then we displace the atomic positions of each atom in the $\Omega$-loop region by (1) a constant cartesian vector of (light blue) 0.5 ångströms (Å), (blue) 1.0 Å, and (dark blue) 2.0 Å, or (2) by a spherically uniform random vector of (pink) 0.5 Å, (red) 1.0 Å, and (dark red) 2.0 Å. Altogether, we have a dataset of $N = 1000$ proteins per simulation scenario: 100 ns interval $\times$ 5 different choices $\delta = \{0.0, 0.1, 0.2, 0.3, 0.4\}$ ns $\times$ 2 phenotypic classes (original wild-type versus perturbed). The area under the curve (AUC) details the ability of SINATRA Pro to identify "true class defining" atoms located within the $\Omega$-loop region as a function of changing the different free parameters used in the algorithm. Here, we assess the robustness of the algorithm to **(a)** $d$ number of directions per cone, **(b)** $c$ number of cones, **(c)** $\theta$ cap radius used to generate directions within each cone, **(d)** $l$ number of sublevel sets (or filtration steps) used to compute the topological summary statistics, and **(e)** the radius cutoff $r$ in Å used to construct the simplicial complex. While varying each parameter, the other parameters are fixed at $\{r = 1.0\,\text{Å}, c = 20, d = 8, \theta = 0.80, l = 120\}$. Guidelines for how to choose the free parameters are given in Table 1 in the main text.

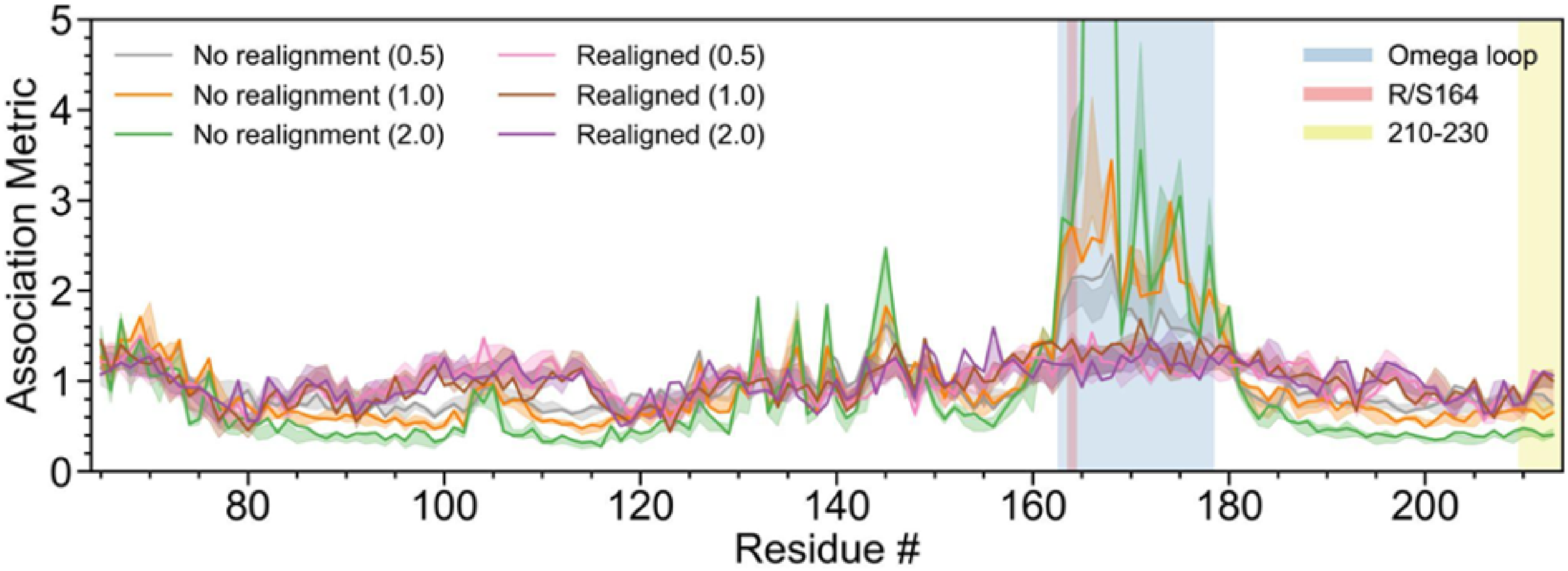

**Figure S2. Effect of realignment after displacement in the controlled experiments aimed at assessing the ability of SINATRA Pro to detect artificial changes in the Ω-loop of $\beta$-lactamase (blue region).** In the main text, we conduct a controlled simulation study where consider two phenotypic classes using real structural data of wild-type $\beta$-lactamase (TEM). In the first phenotypic class, structural protein data are drawn from equally spaced intervals over a 100 ns MD trajectory (e.g., $t_{\mathrm{MD}} = [0, 1, 2, 3, \ldots, 99]$ ns $+ \delta$, where $\delta$ is a time offset parameter). In the second phenotypic class, proteins are drawn from 0.5 ns intervals later relative to the first group (e.g., $t_{\mathrm{MD}} = [0.5, 1.5, 2.5, 3.5, \ldots, 99.5]$ ns $+ \delta$) to introduce physical thermal noise. Here, we displace the atomic positions of each atom in the Ω-loop region by a constant cartesian vector of 0.5, 1.0, and 2.0 ångströms (Å), respectively. The purpose of this figure is to show why we do not realign the proteins after structural perturbation has occurred. Realigning the structures after introducing a perturbation poses a slightly different and notably less controlled simulation study. For example, in this constant displacement case, realigning the structures will shift the whole structure against the perturbing vector and result in an unintentional displacement on the opposite side of the structure (as depicted by the pink, brown, and purple lines). A true positive in this case is not as well-defined as when we keep the unperturbed structure in place and define the perturbed structure as the ground truth for the positive signal (as shown by the grey, orange, and green lines).

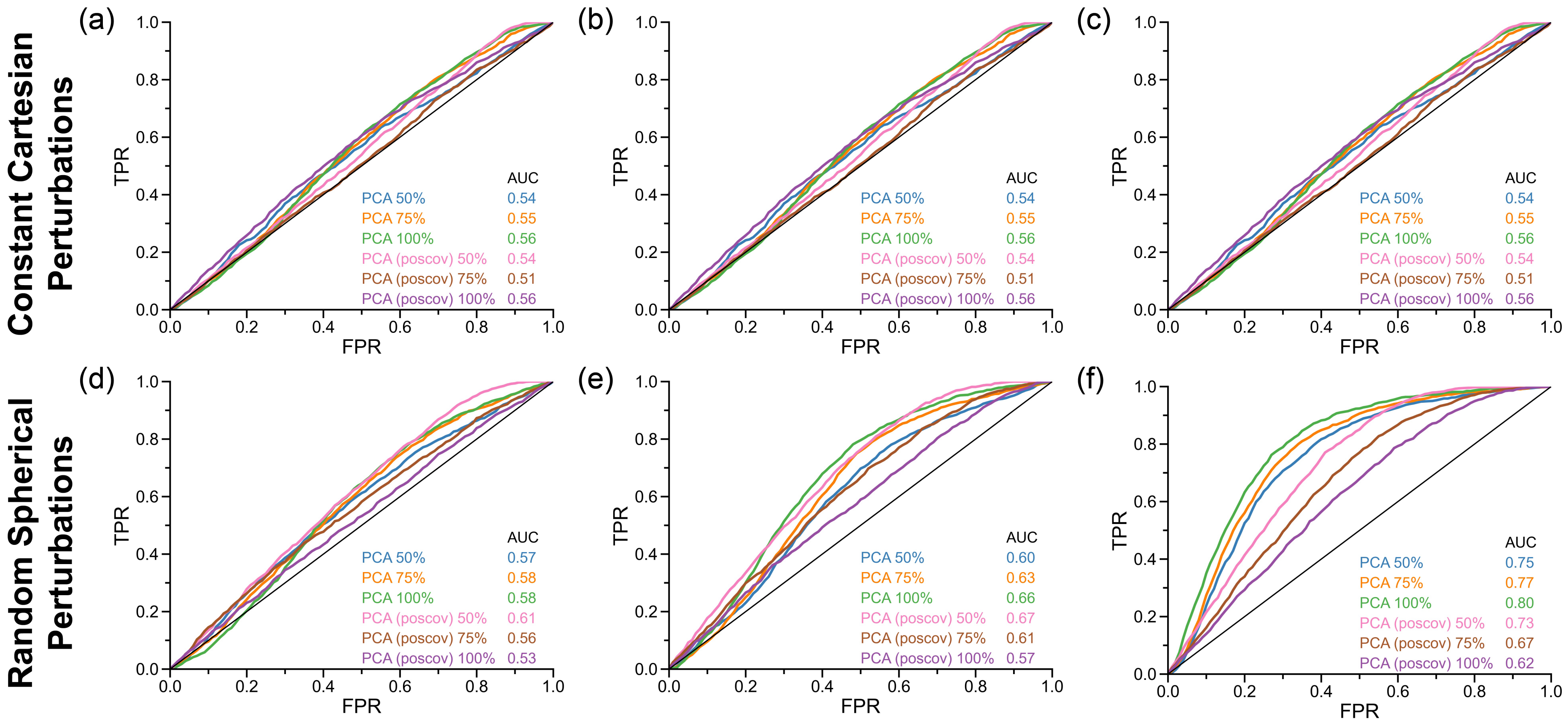

**Figure S3. Receiver operating characteristic (ROC) assessing the differentiating power of PCA under different model parameter configurations in controlled molecular dynamic (MD) simulations.** Here, in addition to the Cartesian-based PCA approach described in the main text, we also perform an additional PCA strategy on the covariance matrix between atomic positions. This is done by first taking the atomic positions of all frames, centering their mean to be zero, and normalizing them to have unit variance equal to one. Next, the position covariance matrix is generated between the two datasets using the function $\mathbb{V}[\mathbf{x}, \mathbf{y}] = \mathbb{E}[(\mathbf{x} - \bar{\mathbf{x}})(\mathbf{y} - \bar{\mathbf{y}})^\intercal]$ which takes on positive values if two variables are correlated and negative if two variables are anti-correlated. We then run PCA on the covariance matrix and choose the number of principal components that explain at least some percentage of its cumulative variance. Note that the output from PCA produces vectors that have dimensionality equal to the total number of atoms in the protein structures, and these can be interpreted as a measure of how explanatory each atomic position is in determining the variation between two sets of data (e.g., class $A$ and $B$, respectively). In this analysis, the principal components are taken as feature vectors to generate the ROC curves, where the least correlated (or anti-correlated) variables with the class labels are considered to be the differentiating features. Above, we are ranking atoms according to the least absolute correlation. In the legend, the approach used to in the main text is simply listed as PCA, while the method run on the positional covariance matrix is indicated by `poscov`. Each method is evaluated using a different number of principal components based on the cumulative variance explained: 50%, 75%, and 100%. We then use area under the curves (AUC) to summarize this performance. Both strategies show similar performance across all simulation scenarios in the constant perturbations, while the original PCA approach that explains 100% of the variation over the Cartesian $(x, y, z)$-coordinates for the atoms is best in the simulations with random spherical perturbations. The latter is what is displayed in the main text.

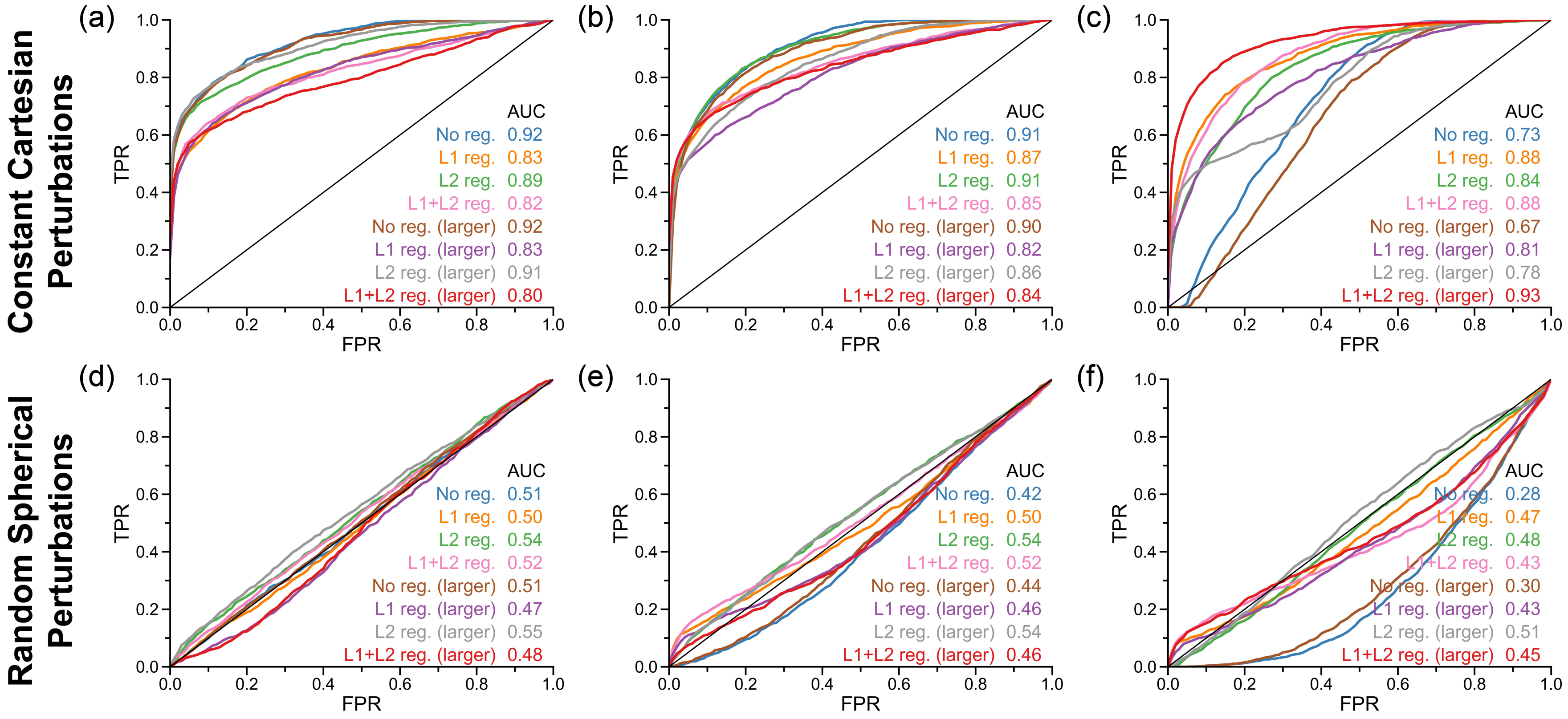

**Figure S4. Receiver operating characteristic (ROC) assessing the differentiating power of the Neural Network under different model parameter and architecture configurations in controlled molecular dynamic (MD) simulations.** Here, in addition to the Neural Network described in the main text (depicted in blue), we also perform an additional search over different architectures and model training procedures. For the former, we try deepening the architecture with Rectified Linear Unit (ReLU) nonlinear activation functions [1] to the following: *(1)* an input layer of Cartesian coordinates of all of the atoms; *(2)* a hidden layer with $H = 2048$ neurons; *(3)* a second hidden layer with $H = 2048$ neurons; *(4)* a third hidden layer with $H = 512$ neurons; *(5)* a third hidden layer with $H = 128$ neurons; and *(6)* an outer layer with a single node which uses a sigmoid link function for protein classification. For the latter modification, we try regularizing the network weights using a combination of $L_1$, $L_2$, and $L_1 + L_2$ penalties. These correspond to the "Least Absolute Shrinkage and Selection Operator" or LASSO solution [2], Ridge Regression [3], and the Elastic Net [4], respectively. Once again, batch normalization was implemented between each layer and a normalized saliency map to rank the importance of each atom [5]. We assess power by taking the sum of the saliency map values corresponding to each atomic position which is summarized by the area under the curves (AUC).

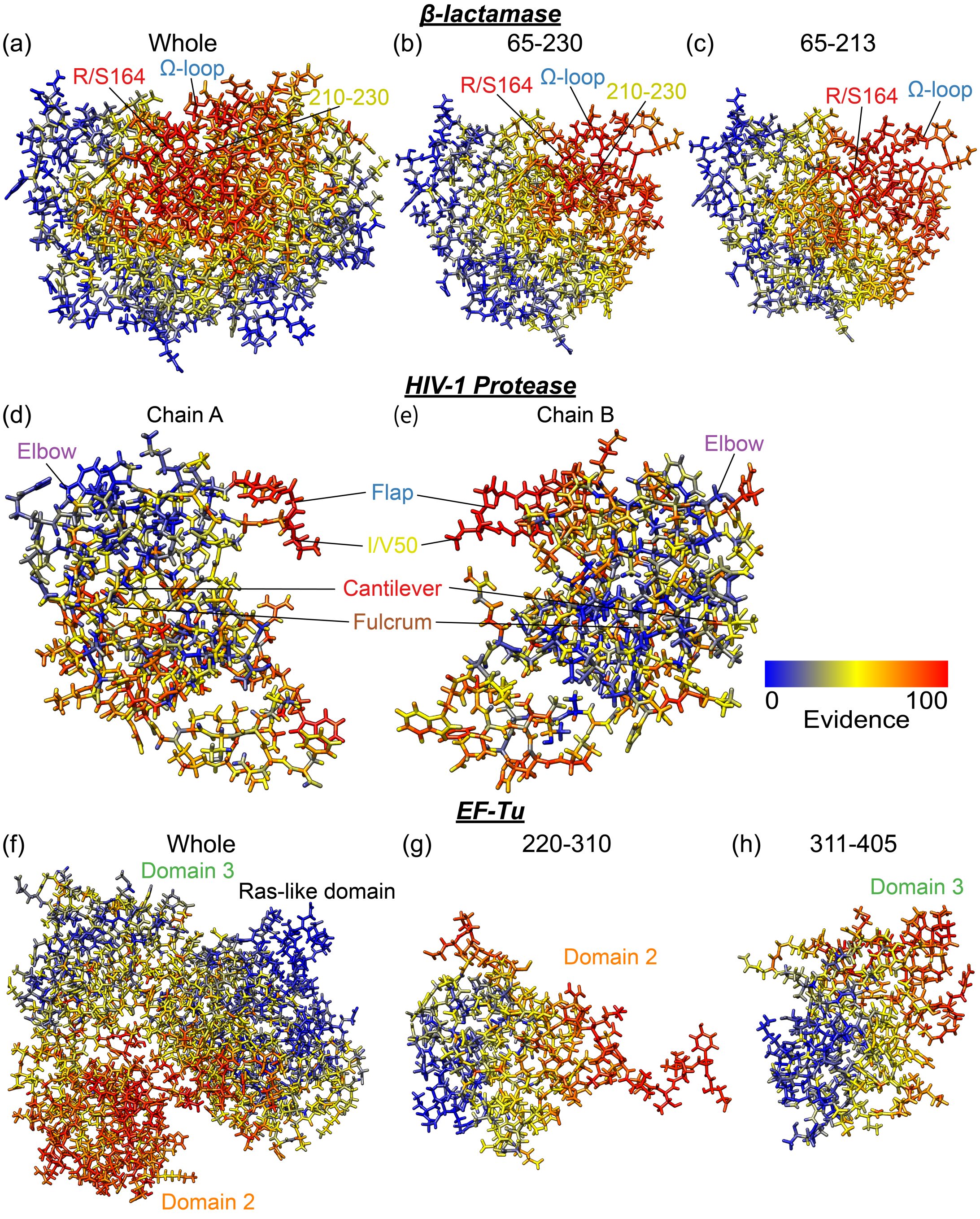

**Figure S5. Atomic-level results for detecting biophysical signatures in (top row) TEM$\beta$-lactamase, (middle row) HIV-1 protease bound to Amprenavir, and (bottom row) GTP-bound EF-Tu.** In these analyses, we compare the molecular dynamics (MD) trajectories of alternative states for each protein to the corresponding trajectories of *(i)* Arg164Ser TEM $\beta$-lactamase, *(ii)* Ile50Val HIV-1 protease bound to Amprenavir, and *(iii)* GDP-bound EF-Tu, respectively. We analyze datasets based on different fragments of each protein. Specifically, in the case of TEM $\beta$-lactamase, we analyze **(a)** the whole protein structure, **(b)** residues 65-230, and **(c)** residues 65-213; in HIV-1 protease, we analyze **(d)** chain A and **(e)** chain B; and, in EF-Tu, we analyze **(f)** the whole protein structure, **(g)** residues 220-310, and **(h)** residues 311-405. Here, consistency between fragments within a protein type shows the robustness of SINATRA Pro to identify the same signal even when it does not have access to the full structure. The heatmaps highlight the atomic evidence potential on a scale from $[0-100]$. A maximum of 100 represents the threshold at which the first atom of the protein is reconstructed, while 0 denotes the threshold when the last atom is reconstructed. Annotated are regions of interest (ROIs) according to literature sources that have previously suggested some level of structural association for each chemical change of interest, including: *(i)* the $\Omega$-loop (residues 163-178) in TEM; *(ii)* the flap region (residues 47-55) in HIV-1 protease; and *(iii)* Domain 2 (residues 208-308) in EF-Tu.

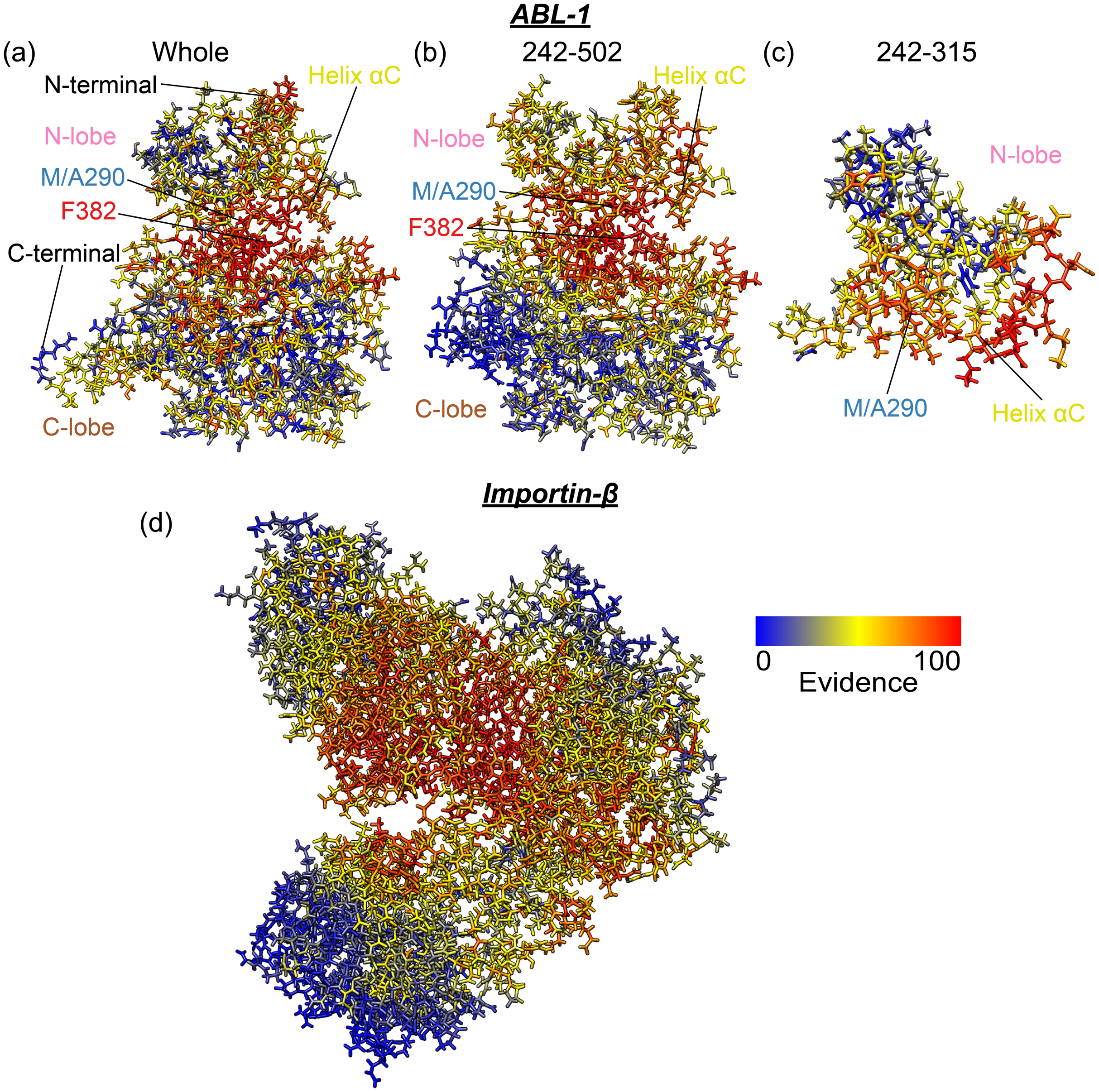

**Figure S6. Atomic-level results for detecting biophysical signatures in (top row) Abl1 and (bottom row) IBB-bound Importin-$\beta$.** In these analyses, we compare the molecular dynamics (MD) trajectories of the alternative states for each protein to the corresponding trajectories of *(i)* Met290Val Abl1 and *(ii)* unbound Importin-$\beta$, respectively. We analyze Abl1 based on different fragments the protein. Specifically, we analyze **(a)** the whole protein structure, **(b)** residues 242-502, and **(c)** residues 242-315. Here, consistency between fragments within a protein type shows the robustness of SINATRA Pro to identify the same signal even when it does not have access to the full structure. The heatmaps highlight the atomic evidence potential on a scale from $[0 - 100]$. A maximum of 100 represents the threshold at which the first atom of the protein is reconstructed, while 0 denotes the threshold when the last atom is reconstructed. Annotated are regions of interest (ROIs) according to literature sources that have previously suggested some level of structural association for each chemical change of interest, including the DFG motif in Abl1. Note that, in the context of Importin-$\beta$, the superhelix includes the entire structure and so we do not include any additional annotations.

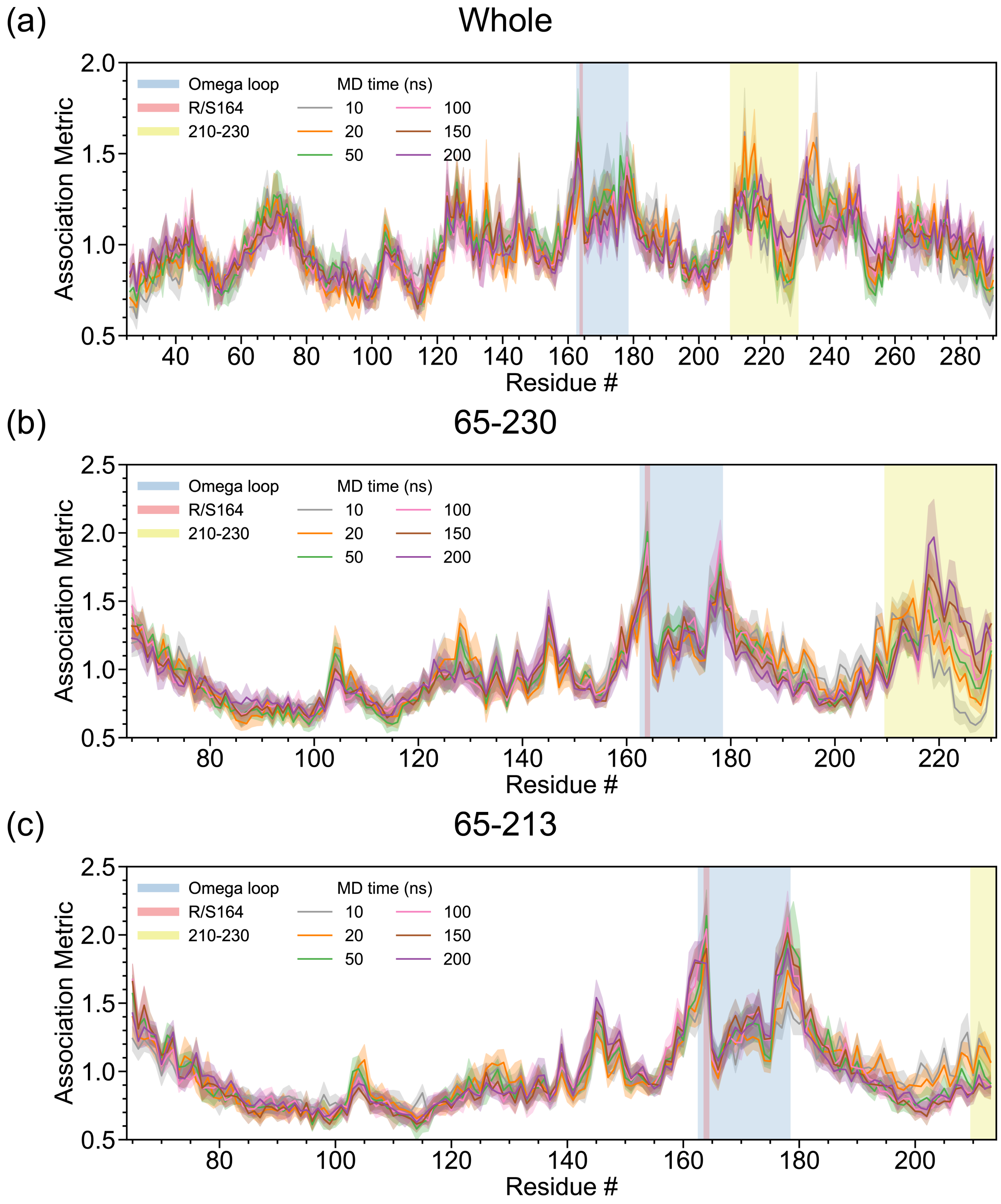

**Figure S7. Sensitivity analyses on different lengths of MD simulations aimed at detecting consistent structural changes in the Ω-loop of TEM $\beta$-lactamase induced by the Arg164Ser mutation using SINATRA Pro.** In this analysis, we compare the molecular dynamics (MD) trajectories of wild-type TEM $\beta$-lactamase versus the Arg164Ser mutant [6, 7]. For both phenotypic classes, structural data are drawn from equally spaced intervals over a 10 ns (grey), 20 ns (orange), 50 ns (green), 100 ns (pink), 150 ns (brown), and 200 ns (purple) MD trajectory. As an example of how data are sampled, in the 150 ns simulation case, we have $t_{\mathrm{MD}} = [0, 1.5, 3, \ldots, 148.5]$ ns$+\delta$, where $\delta = \{0.0, 0.15, 0.3, \ldots, 1.35\}$ ns is a time offset parameter. Panels **(a)-(c)** show the mean association metrics (and their corresponding standard errors) computed for each residue within each analysis (see Material and Methods) with the **(a)** whole protein, **(b)** fragment 65-230, and **(c)** fragment 65-213. The overlap of lines shows the robustness of SINATRA Pro to identify the same signal regardless of trajectory length.

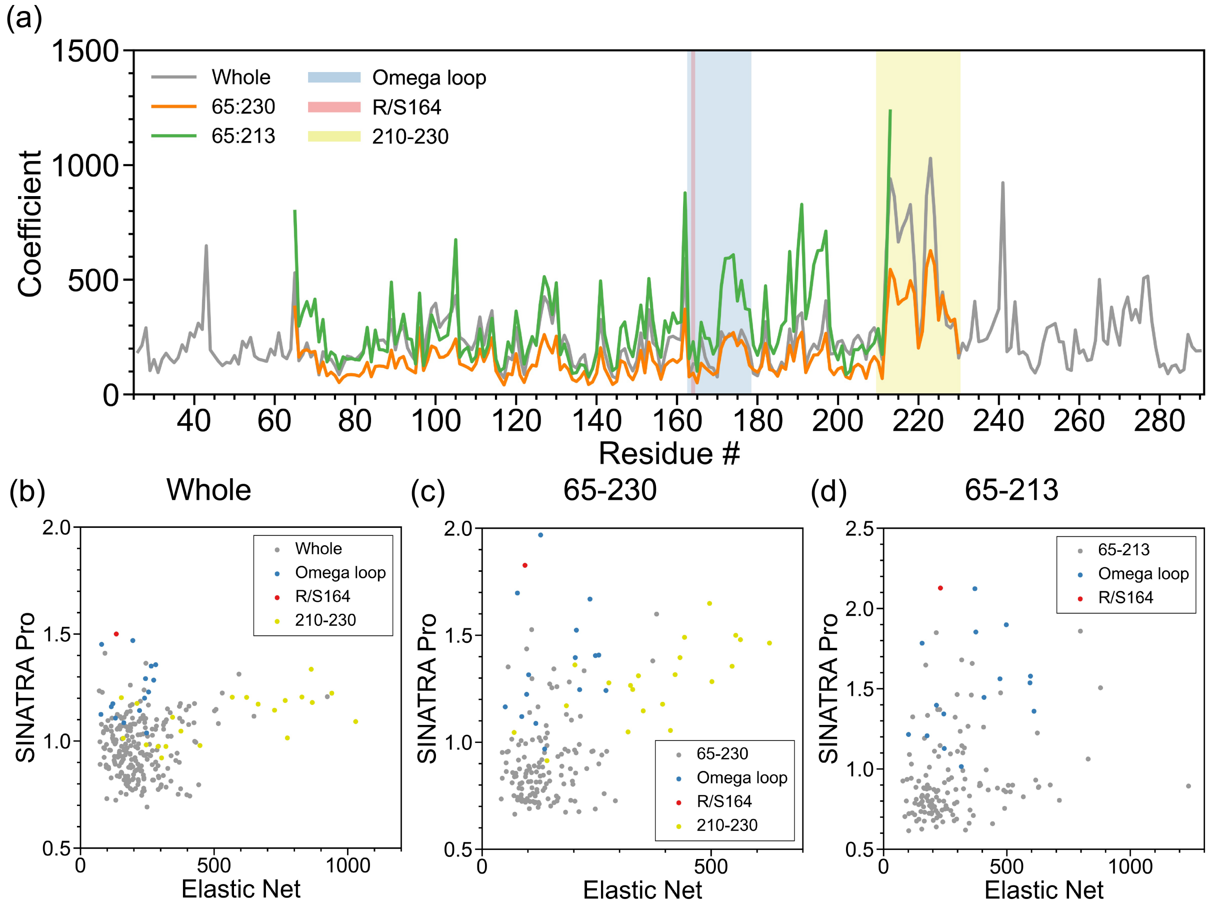

**Figure S8. Real data analyses aimed at detecting structural changes in the Ω-loop of TEM $\beta$-lactamase induced by the Arg164Ser mutation using atomic-level regularization with Elastic Net classification.** In this analysis, we compare the molecular dynamics (MD) trajectories of wild-type TEM $\beta$-lactamase versus the Arg164Ser mutant [6, 7]. For both phenotypic classes, structural data are drawn from equally spaced intervals over a 100 ns MD trajectory (e.g., $t_{\text{MD}} = [0, 1, 2, 3, \ldots, 99]$ ns $+ \delta$, where $\delta$ is a time offset parameter). Altogether, we have a final dataset of $N = 2000$ proteins in the study: 100 ns long interval $\times$ 10 different choices $\delta = \{0.0, 0.1, 0.2, \ldots, 0.9\}$ ns $\times$ 2 phenotypic classes (wild-type versus mutant). To generate these results, we first concatenate the $(x, y, z)$-coordinates of all atoms within each protein and treat them as features in a data frame. Next, we use Elastic Net regularization [4] to assign sparse regression coefficients to each coordinate of every atom (where the penalization term is chosen via cross-validation). Panel **(a)** shows the mean absolute coefficient of all atoms within each residue computed over each fragment-based analysis (see Material and Methods in the main text). The final row plots the correlation between the SINATRA Pro association metrics and the Elastic Net coefficients for all atoms with correspondences in the **(b)** whole protein, **(c)** fragment 65-230, and **(d)** fragment 65-213.

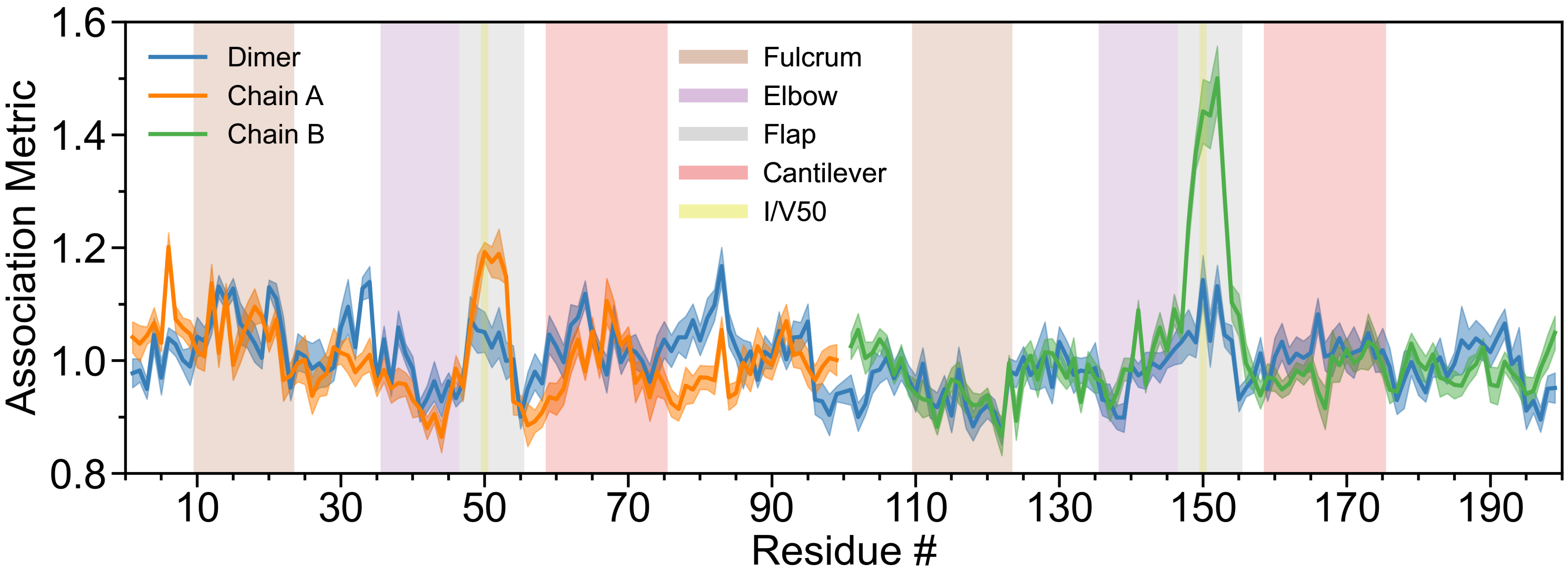

**Figure S9. SINATRA Pro analysis on the dimeric form of HIV-1 protease to detect structural change in the flap region driven by a Ile50Val mutation.** In this analysis, we compare the molecular dynamic (MD) trajectories of wild-type HIV-1 protease versus Ile50Val mutants (i.e., within residues 47-55). For both phenotypic classes, structural data are drawn from from equally spaced intervals over a 100 ns MD trajectory (e.g., $t_{\mathrm{MD}} = [0, 1, 2, 3, \ldots, 99]$ ns + $\delta$, where $\delta$ is a time offset parameter). Altogether, we have a final dataset of $N = 2000$ proteins in the study: 100 ns long interval $\times$ 10 different choices $\delta = \{0.0, 0.1, 0.2, \ldots, 0.9\}$ ns $\times$ 2 phenotypic classes (wild-type versus mutant). This figure depicts results after applying SINATRA Pro using parameters $\{r = 6.0\,\text{Å}, c = 20, d = 8, \theta = 0.80, l = 120\}$ chosen via a grid search. We compare these results to the analyses with chains A and B presented in the main text. Here, the signal in the region of interest (i.e., the flap of each chain) persisted in the dimeric form, but is overshadowed by the noise in the other parts of the protein since the relative orientation between the two monomers causes each chain to be misaligned with itself. Highlighted are residues for regions of the protein corresponding to the fulcrum (brown), elbow (purple), flap (blue), cantilever (red), and I/V50 (yellow) [8–10].

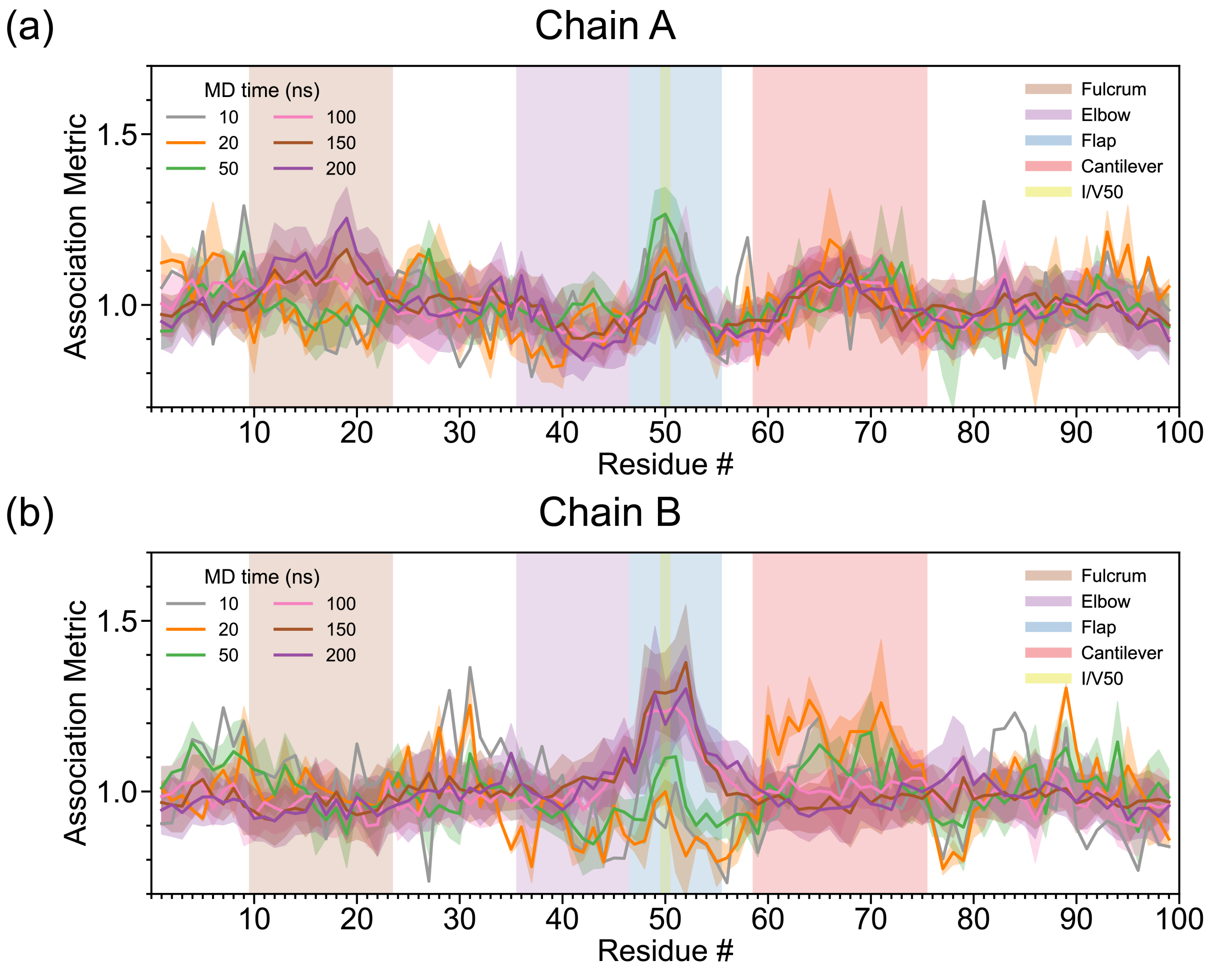

**Figure S10. Sensitivity analyses on different lengths of MD simulations aimed at detecting consistent structural changes in the flap region of HIV-1 protease driven by the Ile50Val mutation using SINATRA Pro.** In this analysis, we compare the molecular dynamics (MD) trajectories of wild-type HIV-1 protease versus the Ile50Val mutant (i.e., within residues 47-55) [8–10]. For both phenotypic classes, structural data are drawn from equally spaced intervals over a 10 ns (grey), 20 ns (orange), 50 ns (green), 100 ns (pink), 150 ns (brown), and 200 ns (purple) MD trajectory. As an example of how data are sampled, in the 150 ns simulation case, we have $t_{\mathrm{MD}} = [0, 1.5, 3, \ldots, 148.5]$ ns $+ \delta$, where $\delta = \{0.0, 0.15, 0.3, \ldots, 1.35\}$ ns is a time offset parameter. Panels **(a)-(c)** show the mean association metrics (and their corresponding standard errors) computed for each residue within each analysis (see Material and Methods) with **(a)** chain A and **(b)** chain B, respectively. The overlap of lines shows the robustness of SINATRA Pro to identify the same signal regardless of trajectory length.

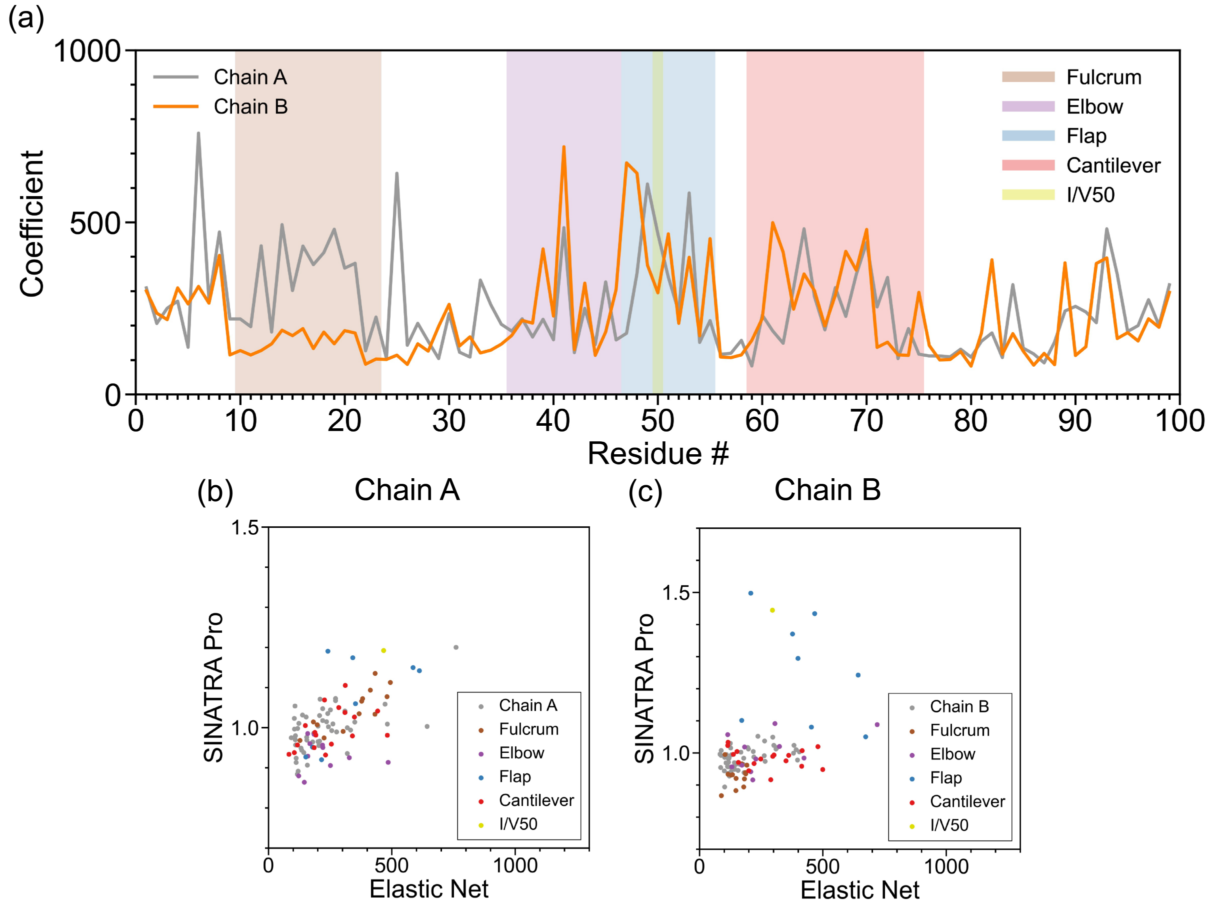

**Figure S11. Real data analyses aimed at detecting structural changes in the flap region of HIV-1 protease driven by the Ile50Val mutation using atomic-level regularization with Elastic Net classification.** In this analysis, we compare the molecular dynamics (MD) trajectories of wild-type HIV-1 protease versus the Ile50Val mutant (i.e., within residues 47-55) [8–10]. For both phenotypic classes, structural data are drawn from equally spaced intervals over a 100 ns MD trajectory (e.g., $t_{\text{MD}} = [0, 1, 2, 3, \ldots, 99]$ ns $+ \delta$, where $\delta$ is a time offset parameter). Altogether, we have a final dataset of $N = 2000$ proteins in the study: 100 ns long interval $\times$ 10 different choices $\delta = \{0.0, 0.1, 0.2, \ldots, 0.9\}$ ns $\times$ 2 phenotypic classes (wild-type versus mutant). To generate these results, we first concatenate the $(x, y, z)$-coordinates of all atoms within each protein and treat them as features in a data frame. Next, we use Elastic Net regularization [4] to assign sparse regression coefficients to each coordinate of every atom (where the penalization term is chosen via cross-validation). Panel **(a)** shows the mean absolute coefficient of all atoms within each residue computed over each fragment-based analysis (see Material and Methods in the main text). The final row plots the correlation between the SINATRA Pro association metrics and the Elastic Net coefficients for all atoms with correspondences in **(b)** chain A and **(c)** chain B, respectively.

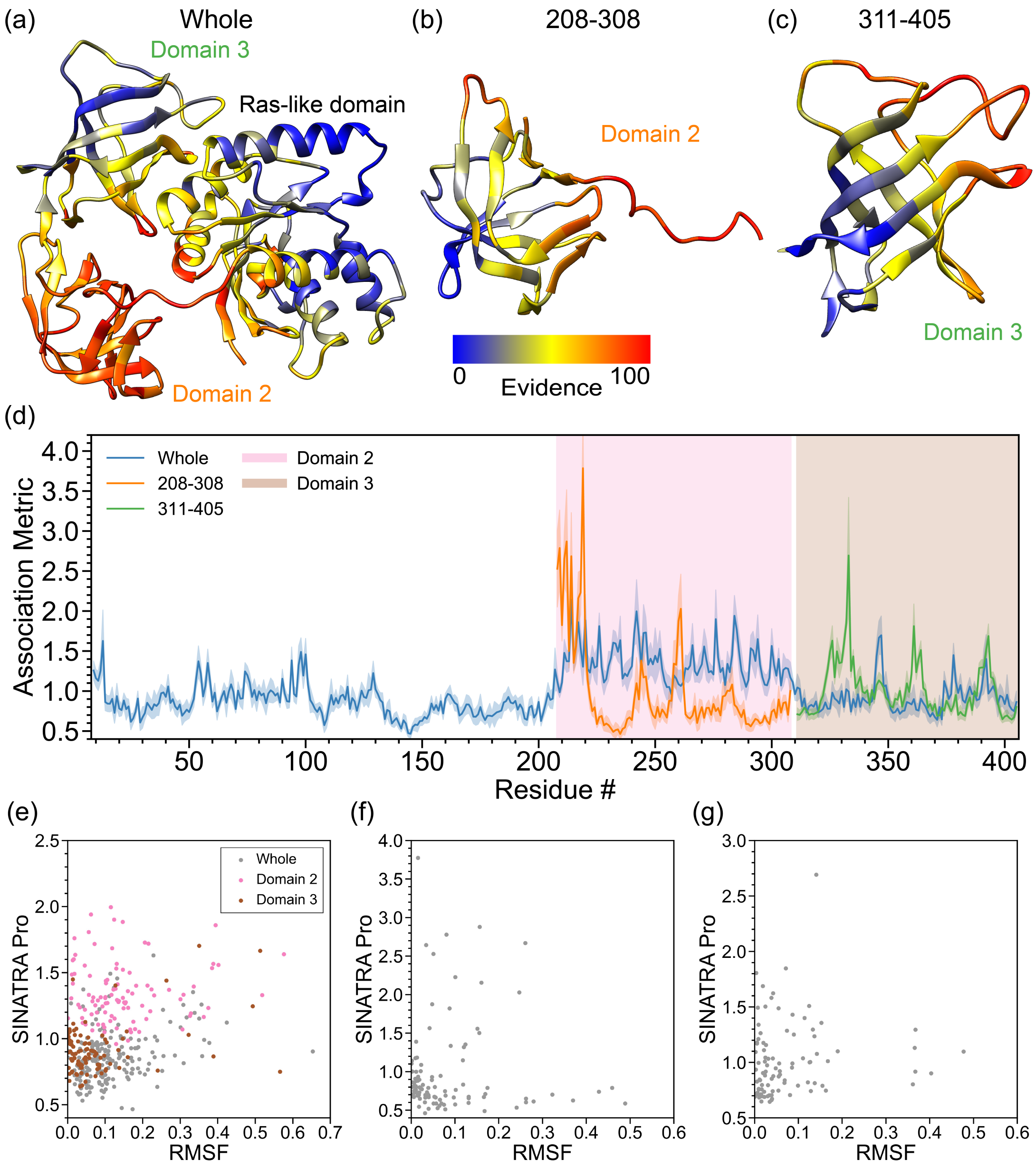

**Figure S12. Real data analyses aimed at detecting structural changes in Domain 2 of the elongation factor EF-Tu upon guanosine triphosphate (GTP) hydrolysis.** In this analysis, we compare the molecular dynamics (MD) trajectories of GTP-bound EF-Tu versus GDP-bound EF-Tu [11–13]. For both phenotypic classes, structural data are drawn from equally spaced intervals over a 100 ns MD trajectory (e.g., $t_{\text{MD}} = [0, 1, 2, 3, \ldots, 99]$ ns$+\delta$, where $\delta$ is a time offset parameter). Altogether, we have a final dataset of $N = 2000$ proteins in the study: 100 ns long interval $\times$ 10 different choices $\delta = \{0.0, 0.1, 0.2, \ldots, 0.9\}$ ns $\times$ 2 phenotypic classes (wild-type versus mutant). This figure depicts results after applying SINATRA Pro using parameters $\{r = 6.0\,\text{Å}, c = 20, d = 8, \theta = 0.80, l = 120\}$ chosen via a grid search. The heatmaps in panels **(a)**-**(c)** highlight residue evidence potential on a scale from $[0-100]$. A maximum of 100 represents the threshold at which the first residue of the protein is reconstructed, while 0 denotes the threshold when the last residue is reconstructed. Panel **(a)** shows residue-level evidence potential when applying SINATRA Pro to the whole protein, while panels **(b)** and **(c)** illustrate results when strictly applying the SINATRA Pro pipeline to atoms in residues 208-308 and 311-405, respectively. Panel **(d)** shows the association metrics (and their corresponding standard errors) computed for each residue within each analysis (see Material and Methods). Here, the overlap shows the robustness of SINATRA Pro to identify the same signal even when it does not have access to the full structure of the protein. The final row plots the correlation between the SINATRA Pro association metrics and the root mean square fluctuation (RMSF) for all atoms with correspondences in the **(e)** whole protein, **(f)** fragment 208-308, and **(g)** fragment 311-405.

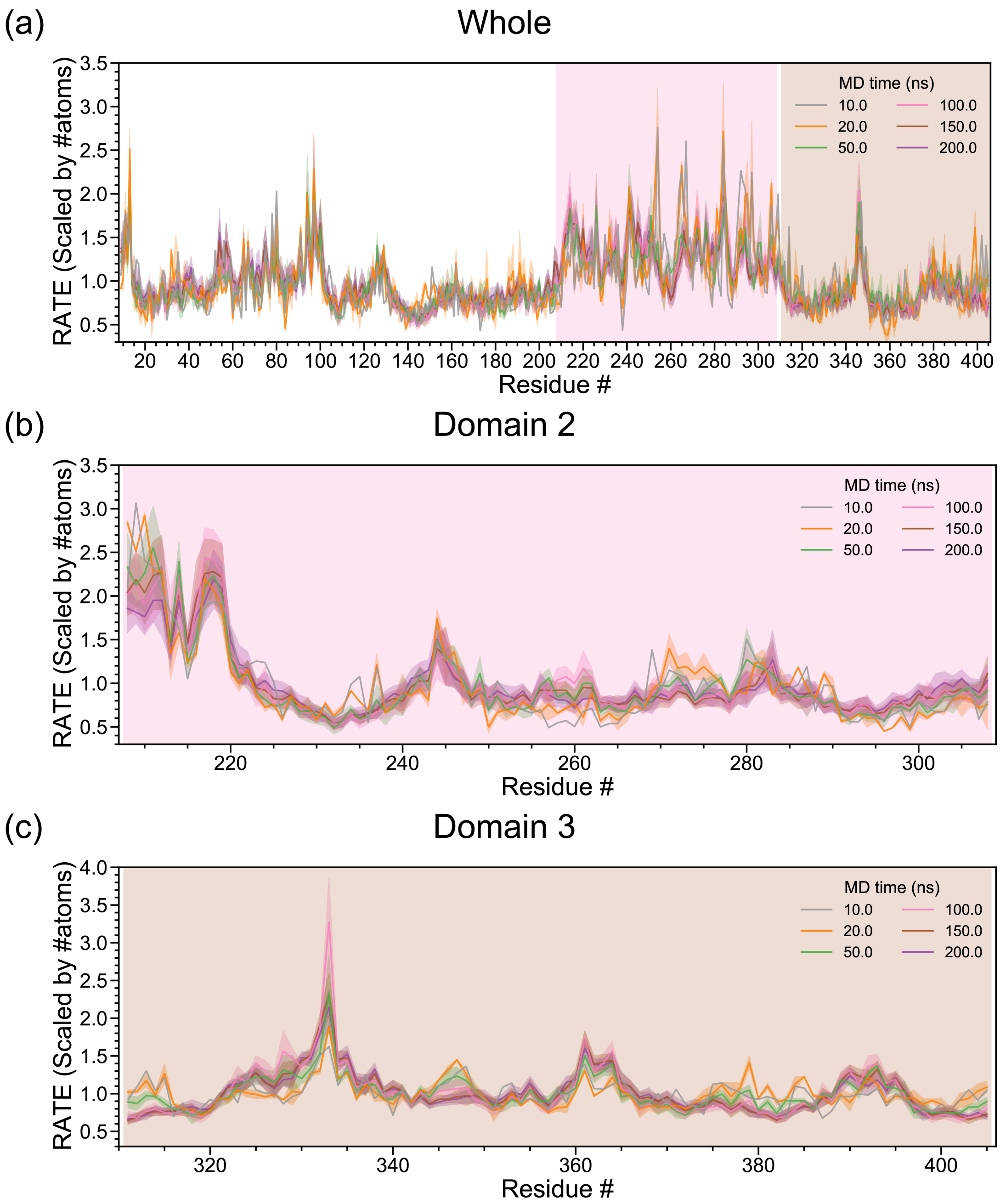

**Figure S13. Sensitivity analyses on different lengths of MD simulations aimed at detecting consistent structural changes in Domain 2 of the elongation factor EF-Tu upon guanosine triphosphate (GTP) hydrolysis using SINATRA Pro.** In this analysis, we compare the the molecular dynamics (MD) trajectories of GTP-bound EF-Tu versus GDP-bound EF-Tu [11–13]. For both phenotypic classes, structural data are drawn from equally spaced intervals over a 10 ns (grey), 20 ns (orange), 50 ns (green), 100 ns (pink), 150 ns (brown), and 200 ns (purple) MD trajectory. As an example of how data are sampled, in the 150 ns simulation case, we have $t_{\text{MD}} = [0, 1.5, 3, \ldots, 148.5]$ ns$+\delta$, where $\delta = \{0.0, 0.15, 0.3, \ldots, 1.35\}$ ns is a time offset parameter. Panels **(a)**-**(c)** show the mean association metrics (and their corresponding standard errors) computed for each residue within each analysis (see Material and Methods) with the **(a)** whole protein, **(b)** fragment 208-308, and **(c)** fragment 311-405. The overlap of lines shows the robustness of SINATRA Pro to identify the same signal regardless of trajectory length.

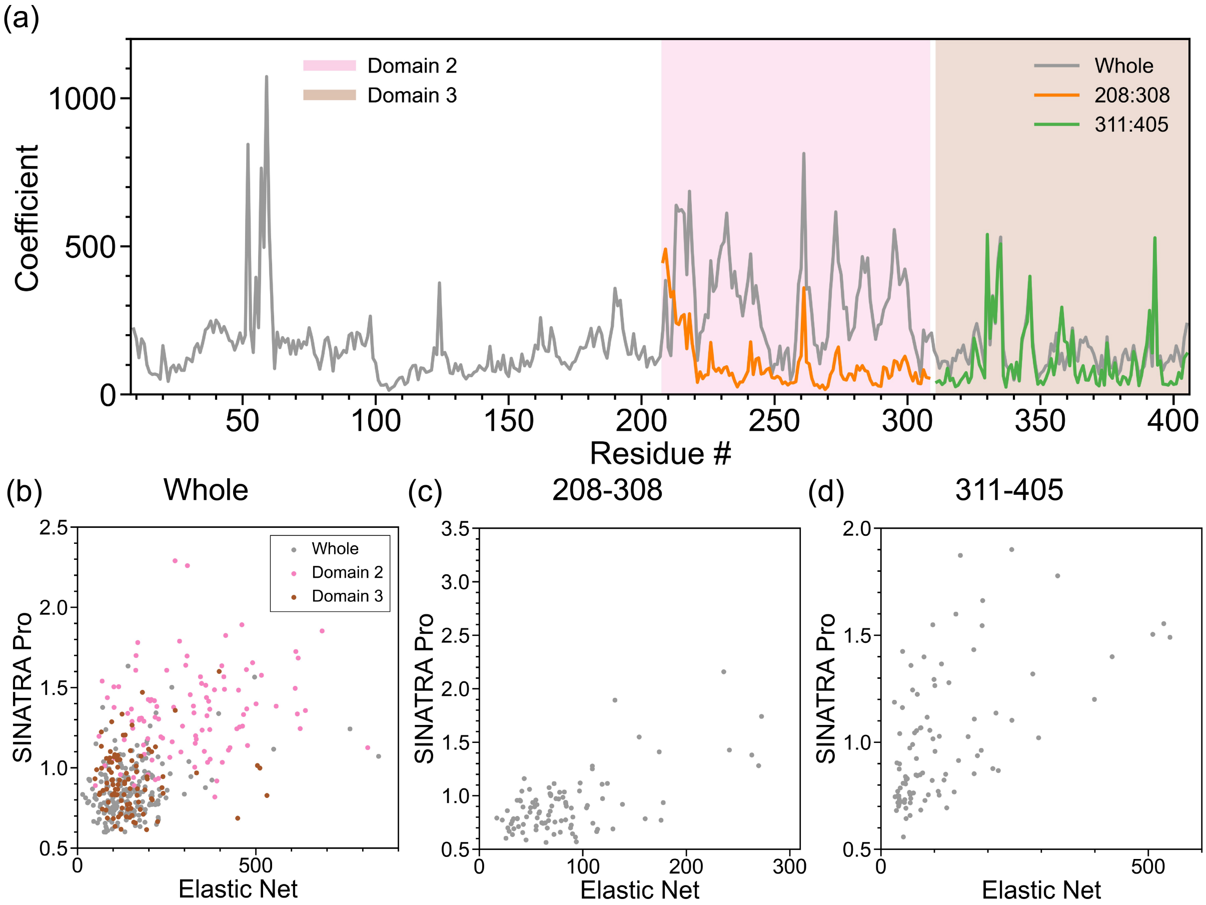

**Figure S14. Real data analyses aimed at detecting structural changes in Domain 2 of the elongation factor EF-Tu upon guanosine triphosphate (GTP) hydrolysis using atomic-level regularization with Elastic Net classification.** In this analysis, we compare the molecular dynamics (MD) trajectories of GTP-bound EF-Tu versus GDP-bound EF-Tu [11–13]. For both phenotypic classes, structural data are drawn from equally spaced intervals over a 100 ns MD trajectory (e.g., $t_{\text{MD}} = [0, 1, 2, 3, \ldots, 99]$ ns $+ \delta$, where $\delta$ is a time offset parameter). Altogether, we have a final dataset of $N = 2000$ proteins in the study: 100 ns long interval $\times$ 10 different choices $\delta = \{0.0, 0.1, 0.2, \ldots, 0.9\}$ ns $\times$ 2 phenotypic classes (wild-type versus mutant). To generate these results, we first concatenate the $(x, y, z)$-coordinates of all atoms within each protein and treat them as features in a data frame. Next, we use Elastic Net regularization [4] to assign sparse regression coefficients to each coordinate of every atom (where the penalization term is chosen via cross-validation). Panel **(a)** shows the mean absolute coefficient of all atoms within each residue computed over each fragment-based analysis (see Material and Methods in the main text). The final row plots the correlation between the SINATRA Pro association metrics and the Elastic Net coefficients for all atoms with correspondences in the **(b)** whole protein, **(c)** fragment 220-310, and **(d)** fragment 311-405.

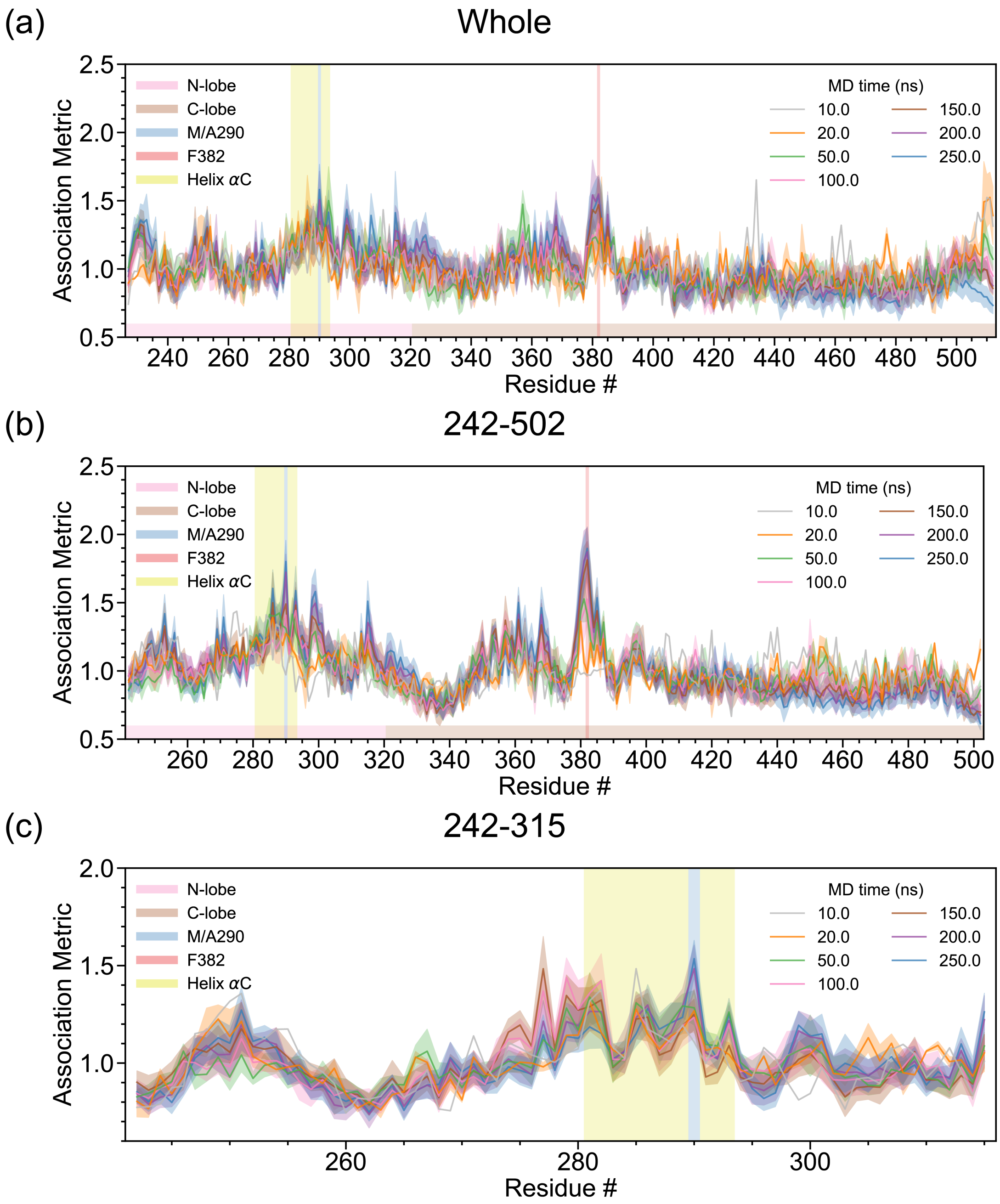

**Figure S15. Sensitivity analyses on different lengths of MD simulations aimed at detecting consistent structural changes in the N-terminal pocket of the Abl1 Tyrosine protein kinase due to the Met290Val mutation in the $\alpha$C helix using SINATRA Pro.** In this analysis, we compare the the molecular dynamics (MD) trajectories of wild-type Abl1 kinase domain versus the Met290Val mutant [14–18]. For both phenotypic classes, structural data are drawn from equally spaced intervals over a 10 ns (grey), 20 ns (orange), 50 ns (green), 100 ns (pink), 150 ns (brown), and 200 ns (purple) MD trajectory. As an example of how data are sampled, in the 150 ns simulation case, we have $t_{\text{MD}} = [0, 1.5, 3, \ldots, 148.5]$ ns $+\delta$, where $\delta = \{0.0, 0.15, 0.3, \ldots, 1.35\}$ ns is a time offset parameter. Panels **(a)-(c)** show the mean association metrics (and their corresponding standard errors) computed for each residue within each analysis (see Material and Methods) with the **(a)** whole protein, **(b)** fragment 242-502, and **(c)** fragment 242-315. The overlap of lines shows the robustness of SINATRA Pro to identify the same signal regardless of trajectory length.

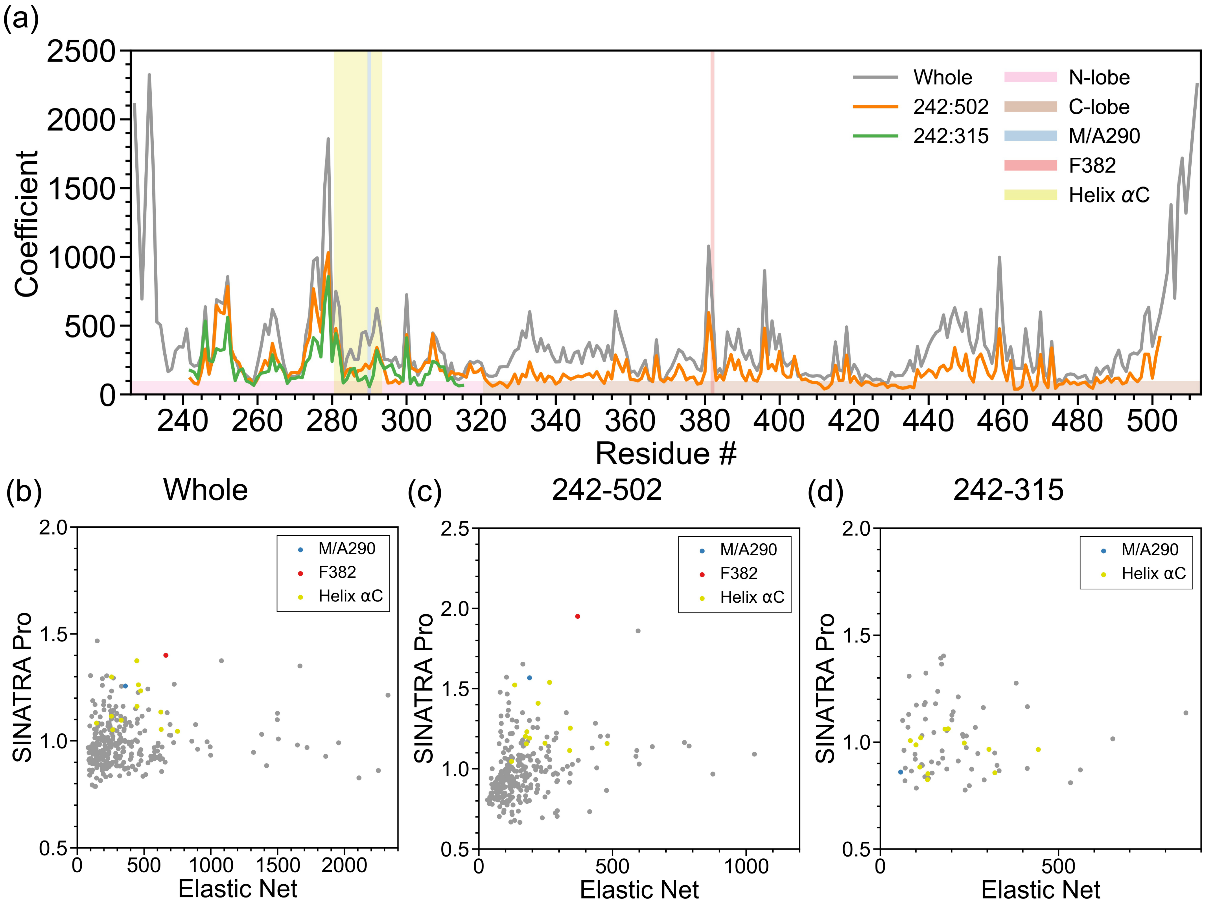

**Figure S16. Real data analyses aimed at detecting structural changes in the N-terminal pocket of the Abl1 Tyrosine protein kinase due to the Met290Val mutation in the $\alpha$C helix using atomic-level regularization with Elastic Net classification.** In this analysis, we compare the molecular dynamics (MD) trajectories of wild-type Abl1 kinase domain versus the Met290Val mutant. [14–18]. For both phenotypic classes, structural data are drawn from equally spaced intervals over a 150 ns MD trajectory (e.g., $t_{\text{MD}} = [0, 1, 2, 3, \ldots, 99] \times 1.5$ ns$+\delta$, where $\delta$ is a time offset parameter). Altogether, we have a final dataset of $N = 3000$ proteins in the study: 150 ns long interval $\times$ 15 different choices $\delta = \{0.0, 0.1, 0.2, \ldots, 1.4\}$ ns $\times$ 2 phenotypic classes (wild-type versus mutant). To generate these results, we first concatenate the $(x, y, z)$-coordinates of all atoms within each protein and treat them as features in a data frame. Next, we use Elastic Net regularization [4] to assign sparse regression coefficients to each coordinate of every atom (where the penalization term is chosen via cross-validation). Panel **(a)** shows the mean absolute coefficient of all atoms within each residue computed over each fragment-based analysis (see Material and Methods in the main text). The final row plots the correlation between the SINATRA Pro association metrics and the Elastic Net coefficients for all atoms with correspondences in the **(b)** whole protein, **(c)** fragment 242-502, and **(d)** fragment 242-315.

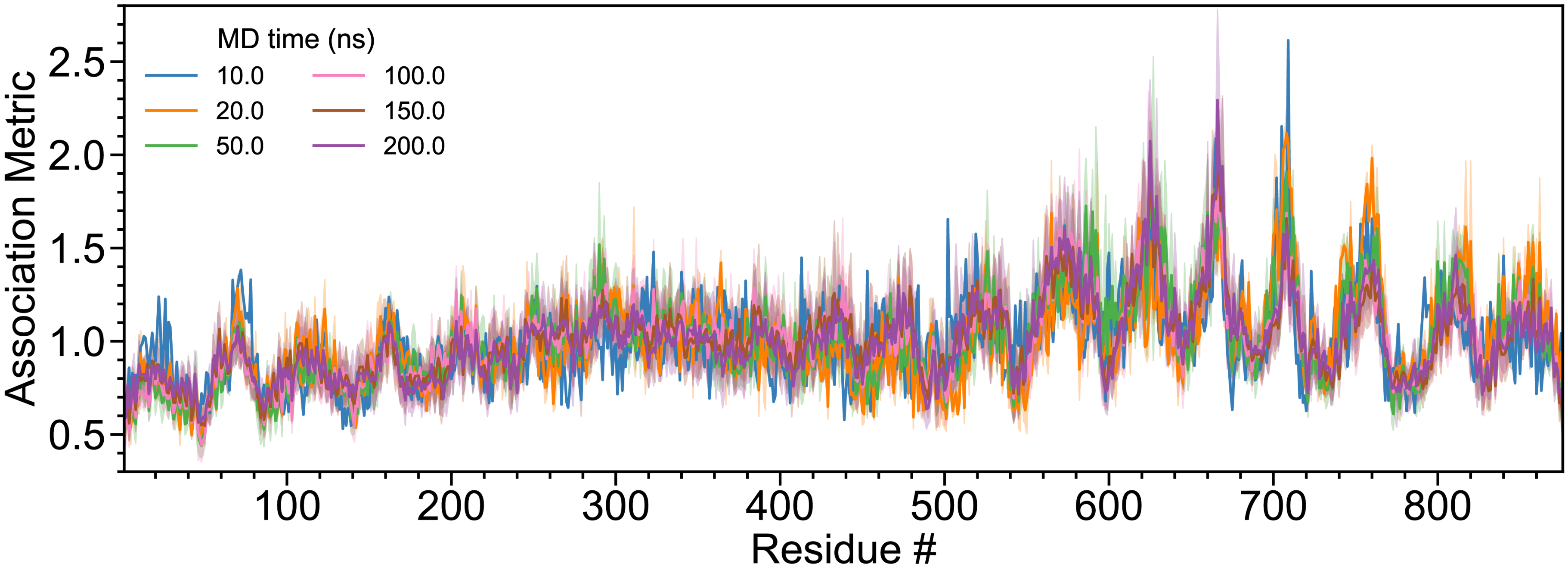

**Figure S17. Sensitivity analyses on different lengths of MD simulations aimed at detecting uncoiling of the superhelix in Importin-$\beta$ upon release of an IBB peptide using SINATRA Pro.** In this analysis, we compare the the molecular dynamics (MD) trajectories of IBB-bound Importin-$\beta$ versus unbound Importin-$\beta$ [19–21]. For both phenotypic classes, structural data are drawn from equally spaced intervals over a 10 ns (grey), 20 ns (orange), 50 ns (green), 100 ns (pink), 150 ns (brown), and 200 ns (purple) MD trajectory. As an example of how data are sampled, in the 150 ns simulation case, we have $t_{\mathrm{MD}} = [0, 1.5, 3, \ldots, 148.5]$ ns + $\delta$, where $\delta = \{0.0, 0.15, 0.3, \ldots, 1.35\}$ ns is a time offset parameter. Panels **(a)-(c)** show the mean association metrics (and their corresponding standard errors) computed for each residue within each analysis (see Material and Methods). The overlap of lines shows the robustness of SINATRA Pro to identify the same signal regardless of trajectory length.

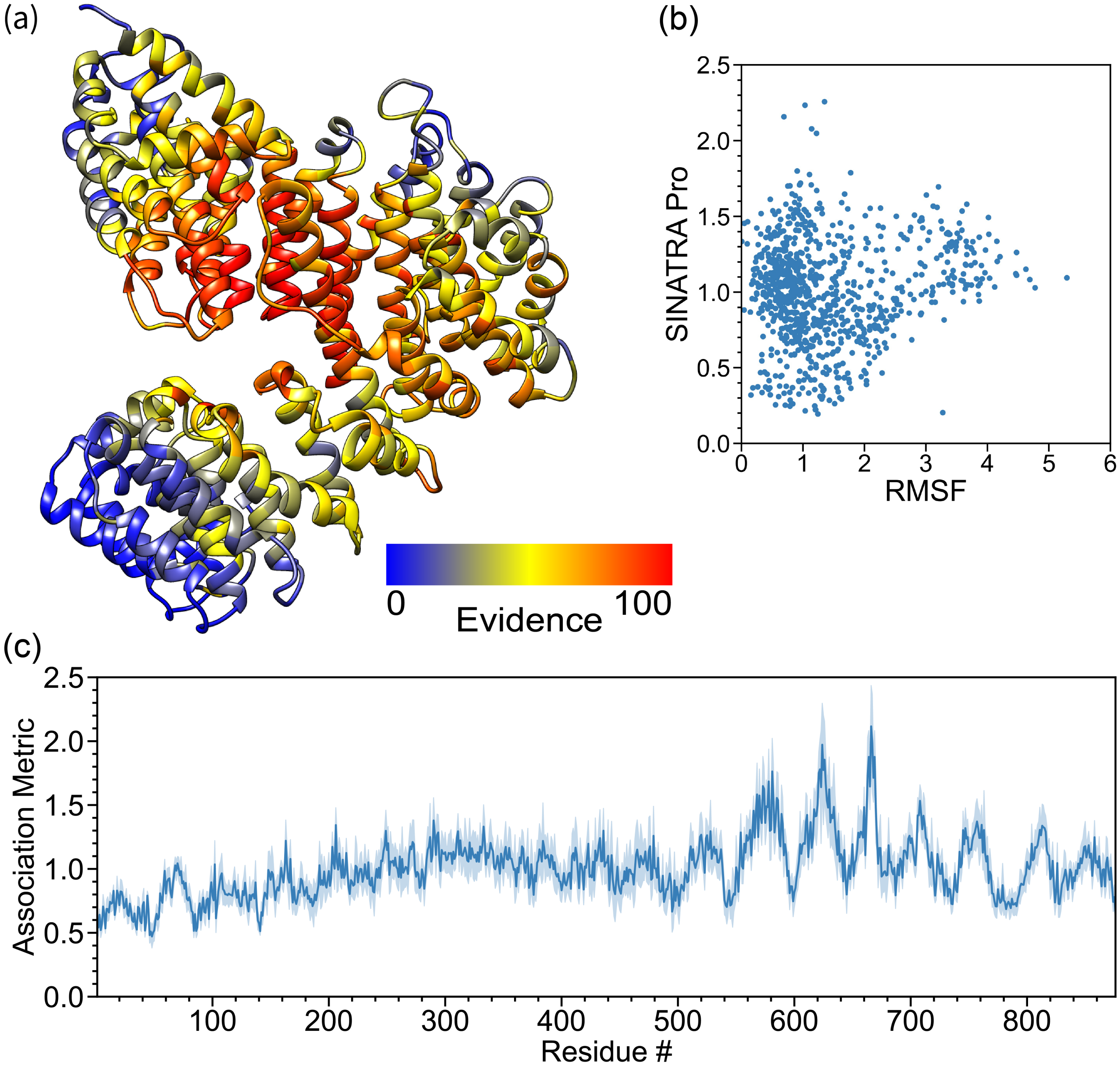

**Figure S18. Real data analyses aimed at detecting uncoiling of the superhelix in Importin-$\beta$ upon release of an IBB peptide.** In this analysis, we compare the molecular dynamics (MD) trajectories of IBB-bound Importin-$\beta$ versus unbound Importin-$\beta$ [19–21]. For both phenotypic classes, structural data are drawn from equally spaced intervals over a 100 ns MD trajectory (e.g., $t_{\text{MD}} = [0, 1, 2, 3, \ldots, 99]$ ns $+ \delta$, where $\delta$ is a time offset parameter). Altogether, we have a final dataset of $N = 2000$ proteins in the study: 100 ns long interval $\times$ 10 different choices $\delta = \{0.0, 0.1, 0.2, \ldots, 0.9\}$ ns $\times$ 2 phenotypic classes (wild-type versus mutant). This figure depicts results after applying SINATRA Pro using parameters $\{r = 6.0\,\text{Å}, c = 20, d = 8, \theta = 0.80, l = 120\}$ chosen via a grid search. The heatmap in panels **(a)** highlights residue evidence potential on a scale from $[0 - 100]$. A maximum of 100 represents the threshold at which the first residue of the protein is reconstructed, while 0 denotes the threshold when the last residue is reconstructed. Panel **(b)** plots the correlation between the SINATRA Pro association metrics and the root mean square fluctuation (RMSF) for all atoms with correspondences. Panel **(c)** shows the SINATRA Pro association metrics (and their corresponding standard errors) computed for each residue within the analysis (see Material and Methods in the main text for more details).

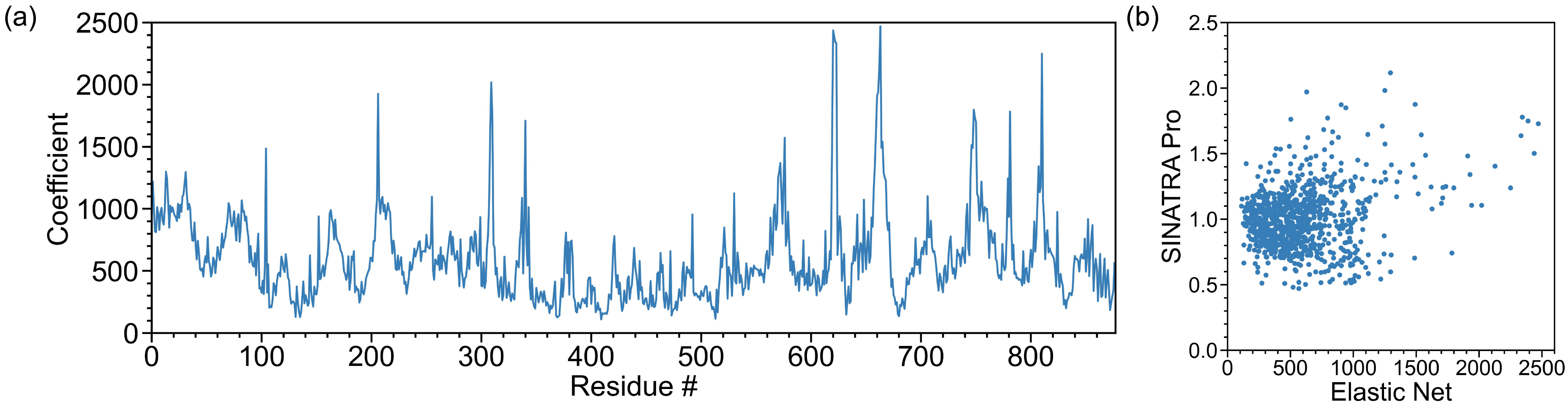

**Figure S19. Real data analyses at detecting uncoiling of the superhelix in Importin-$\beta$ upon release of IBB using atomic-level regularization with Elastic Net classification.** In this analysis, we compare the molecular dynamics (MD) trajectories of IBB-bound Importin-$\beta$ versus unbound Importin-$\beta$ [19–21]. For both phenotypic classes, structural data are drawn from equally spaced intervals over a 100 ns MD trajectory (e.g., $t_{\text{MD}} = [0, 1, 2, 3, \ldots, 99]$ ns + $\delta$, where $\delta$ is a time offset parameter). Altogether, we have a final dataset of $N = 2000$ proteins in the study: 100 ns long interval $\times$ 10 different choices $\delta = \{0.0, 0.1, 0.2, \ldots, 0.9\}$ ns $\times$ 2 phenotypic classes (wild-type versus mutant). To generate these results, we first concatenate the $(x, y, z)$-coordinates of all atoms within each protein and treat them as features in a data frame. Next, we use Elastic Net regularization [4] to assign sparse regression coefficients to each coordinate of every atom (where the penalization term is chosen via cross-validation). Panel **(a)** shows the mean absolute coefficient of all atoms within each residue computed over each fragment-based analysis (see Material and Methods in the main text). In panel **(b)**, we plot the correlation between the SINATRA Pro association metrics and the Elastic Net coefficients for all atoms with correspondences.

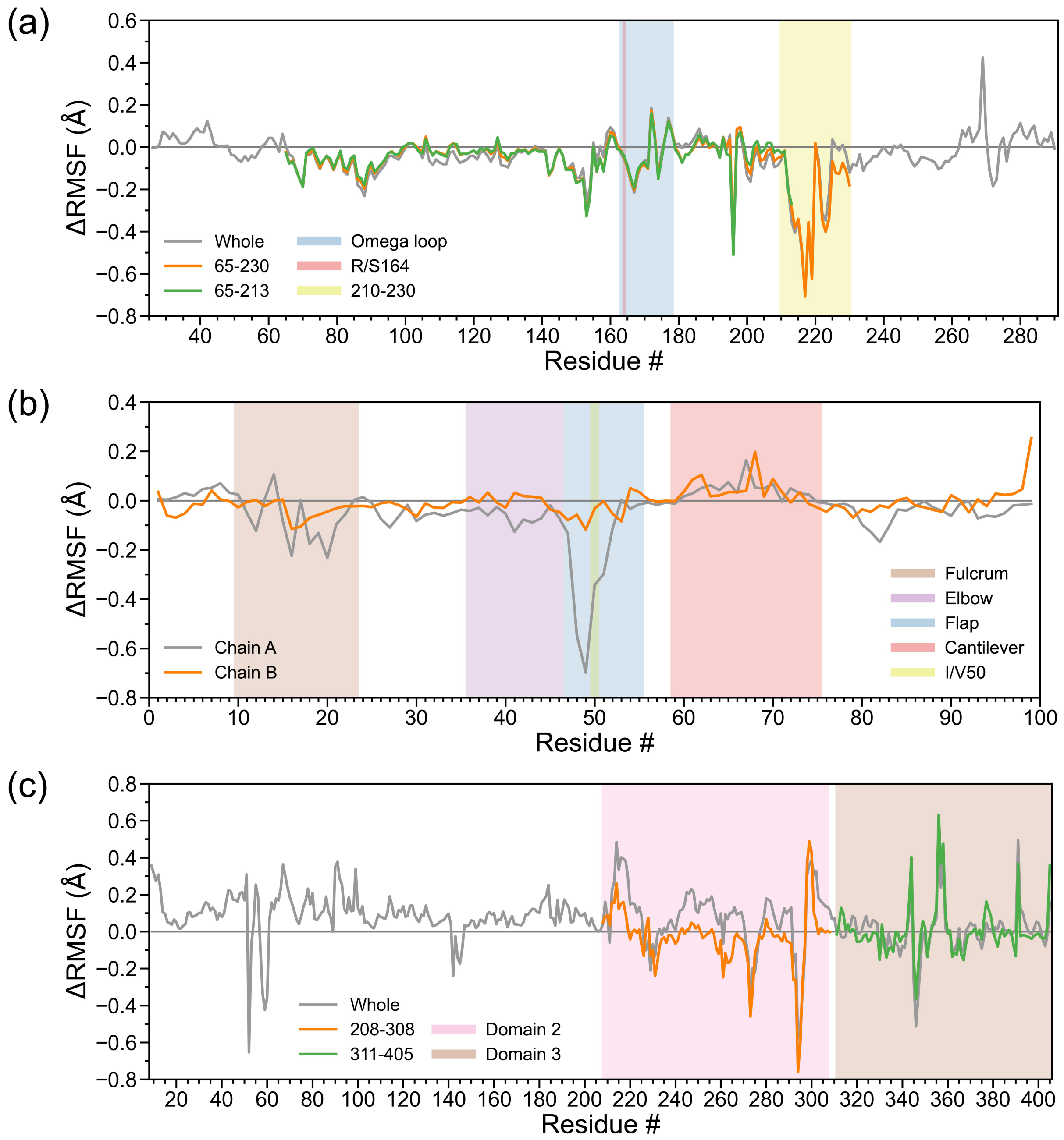

**Figure S20. Residue-level results from running root mean square fluctuation (RMSF) analysis on TEM $\beta$-lactamase, HIV-1 protease, and GTP-bound EF-Tu.** In these analyses, we compare the molecular dynamics (MD) trajectories of the alternative state for each protein to the corresponding trajectories of **(a)** Arg164Ser mutant $\beta$-lactamase, **(b)** Ile50Val mutant HIV-1 protease, and **(c)** GDP-bound EF-Tu, respectively. We analyze datasets based on different fragments of each protein. Specifically, **(a)** in TEM $\beta$-lactamase, we analyze the whole protein structure, residues 65-230, and residues 65-213; **(b)** in HIV-1 protease, we analyze chain A and chain B; and, **(c)** in EF-Tu, we analyze the whole protein structure, residues 220-310, and residues 311-405. The $y$-axis denotes the absolute difference (or $\Delta$-change) in RMSF between wild-type and mutants classes.

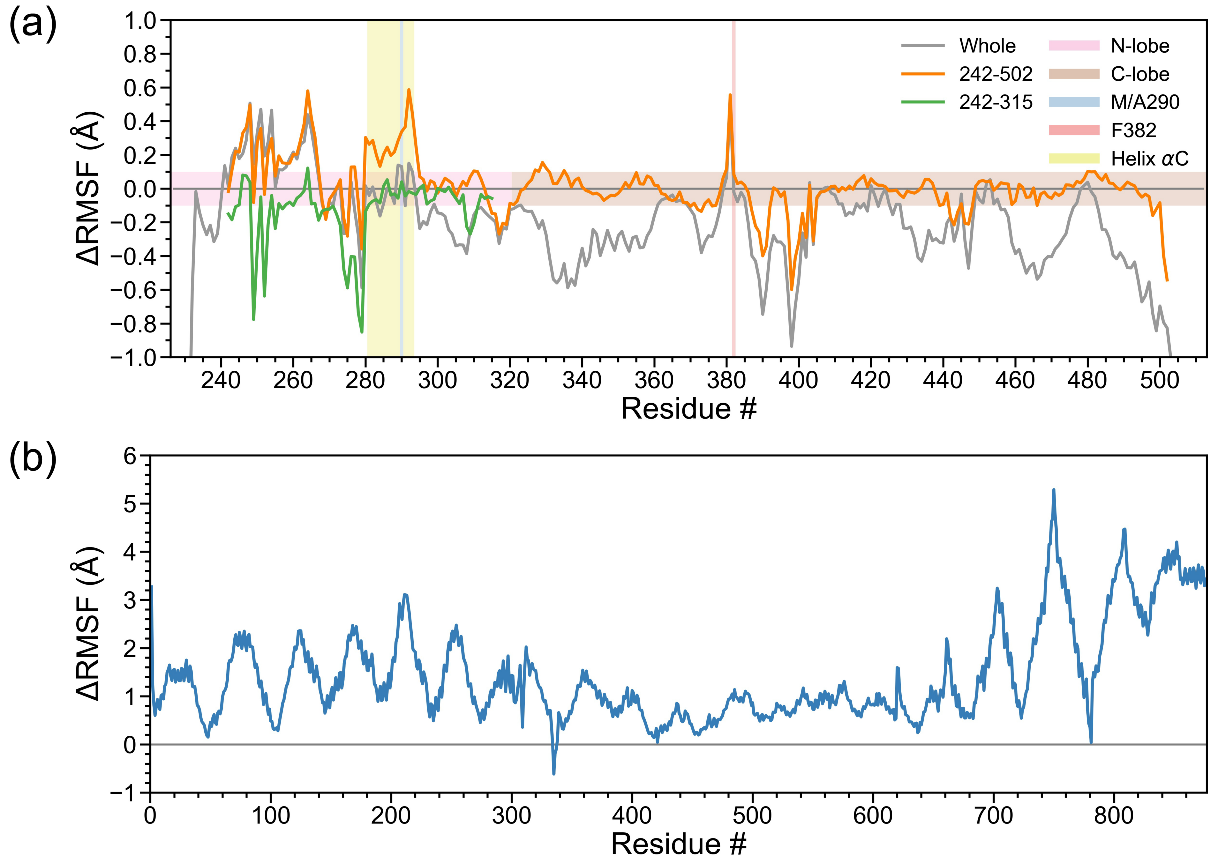

**Figure S21. Residue-level results from running root mean square fluctuation (RMSF) analysis on Abl1 and IBB-bound Importin-$\beta$.** In these analyses, we compare the molecular dynamics (MD) trajectories of the alternative state for each protein to the corresponding trajectories of **(a)** Met290Val Abl1 and **(b)** unbound Importin-$\beta$, respectively. We analyze Abl1 based on different fragments the protein. Specifically, we analyze the whole protein structure, residues 242-502, and residues 242-315. Note that, in the context of Importin-$\beta$, the superhelix includes the entire structure and so we do include any additional sub-fragment analyses. The $y$-axis on denotes the absolute difference (or $\Delta$-change) in RMSF between wild-type and mutants classes.

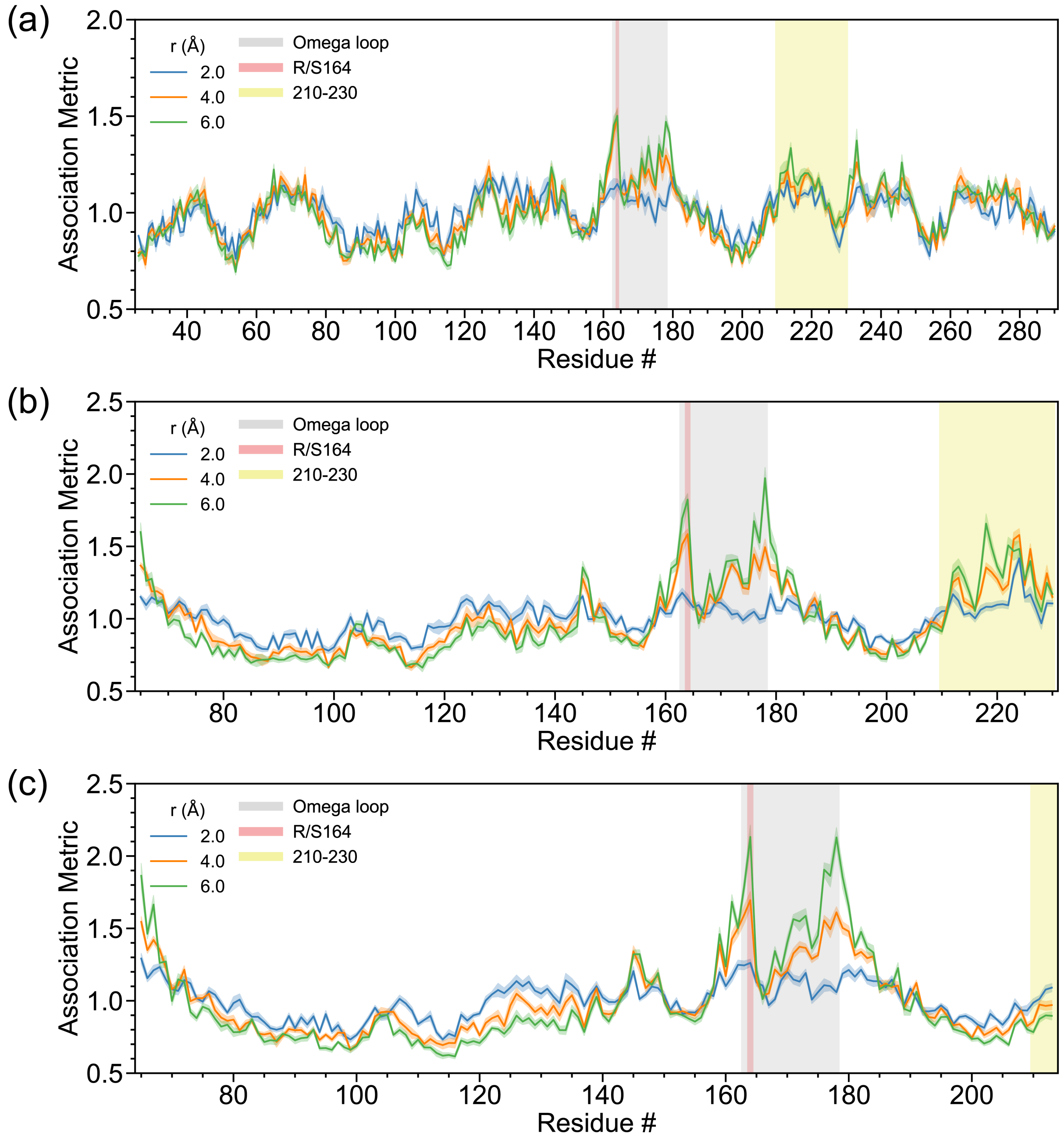

**Figure S22. Sensitivity analysis assessing the robustness of SINATRA Pro to different radius cutoffs $r$ values used to construct the simplicial complexes for TEM $\beta$-lactamase.** Recall that we use the atomic positions for each protein to create mesh representations of their 3D structures (see Fig. 1 in the main text). First, we draw an edge between any two atoms if their Euclidean distance smaller than some value $r$, namely $\text{dist}|(x_1, y_1, z_1), (x_2, y_2, z_2)| < r$. Next, we fill in all the triangles (or faces) formed by these connected edges. We treat the resulting triangulated mesh as an simplicial complex with which we can perform topological data analysis. Here, we consider the construction of mesh representations for each protein while setting $r = \{2.0, 4.0, 6.0\}$ ångströms (Å). Other SINATRA parameters were fixed: $c = 20$ cones, $d = 8$ directions per cone, $\theta = 0.80$ cap radius used to generate directions in a cone, and $l = 120$ sublevel sets per filtration. In each plot, we show the association metrics (and their corresponding standard errors) computed for each residue while analyzing **(a)** the whole protein, **(b)** fragment 65-230, and **(c)** fragment 65-213. Note that an overlap in signal shows the robustness of SINATRA Pro to this input parameter value.

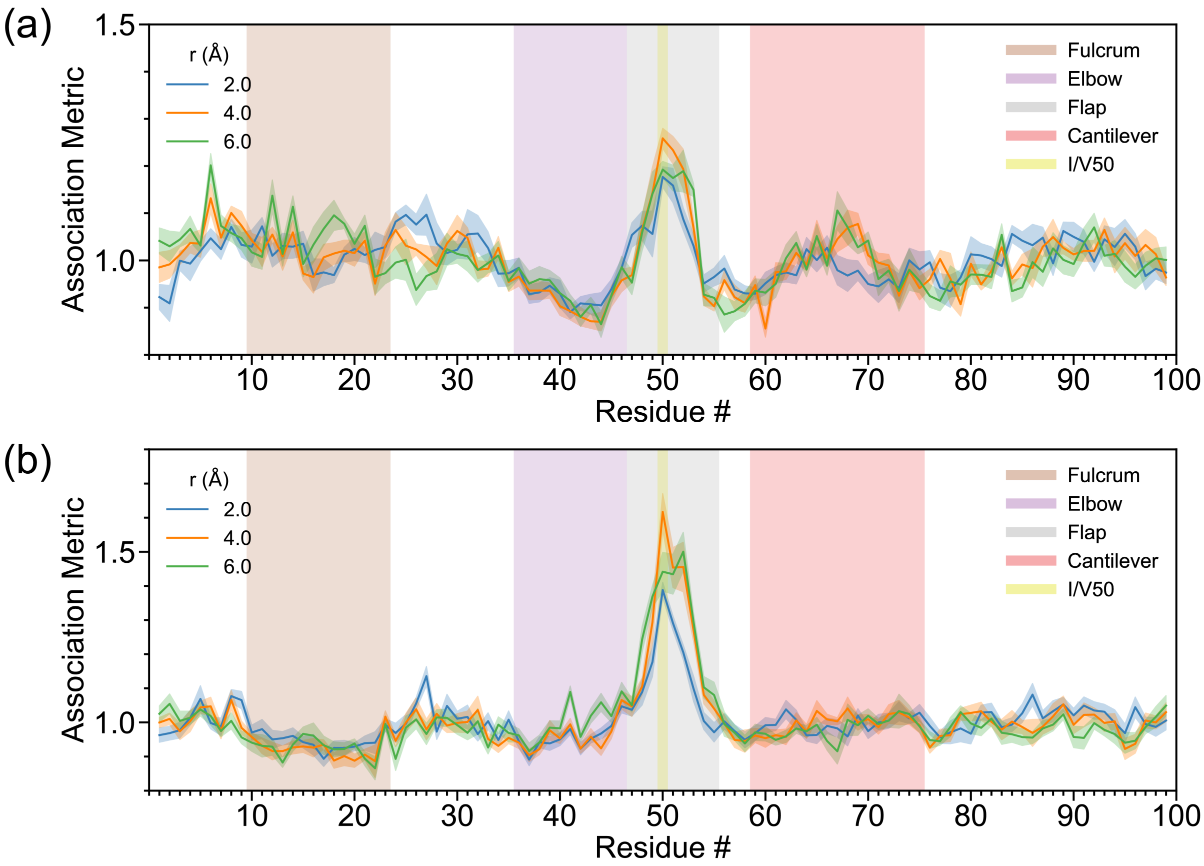

**Figure S23. Sensitivity analysis assessing the robustness of SINATRA Pro to different radius cutoffs $r$ values used to construct the simplicial complexes for HIV-1 protease.** Recall that we use the atomic positions for each protein to create mesh representations of their 3D structures (see Fig. 1 in the main text). First, we draw an edge between any two atoms if their Euclidean distance smaller than some value $r$, namely $\text{dist}|(x_1, y_1, z_1), (x_2, y_2, z_2)| < r$. Next, we fill in all the triangles (or faces) formed by these connected edges. We treat the resulting triangulated mesh as an simplicial complex with which we can perform topological data analysis. Here, we consider the construction of mesh representations for each protein while setting $r = \{2.0, 4.0, 6.0\}$ ångströms (Å). Other SINATRA parameters were fixed: $c = 20$ cones, $d = 8$ directions per cone, $\theta = 0.80$ cap radius used to generate directions in a cone, and $l = 120$ sublevel sets per filtration. In each plot, we show the association metrics (and their corresponding standard errors) computed for each residue while analyzing **(a)** chain A and **(b)** chain B. Note that an overlap in signal shows the robustness of SINATRA Pro to this input parameter value.

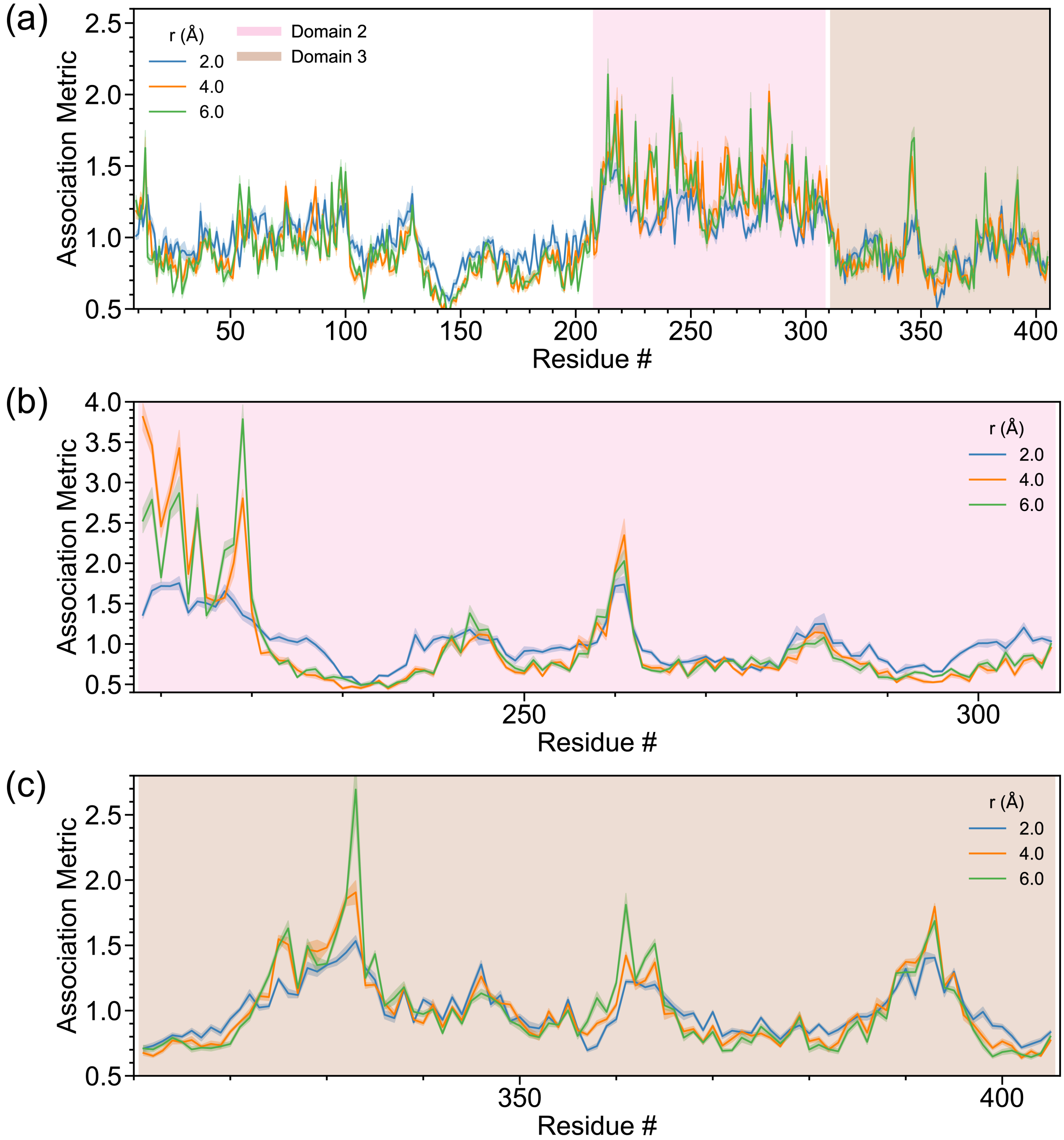

**Figure S24. Sensitivity analysis assessing the robustness of SINATRA Pro to different radius cutoffs $r$ values used to construct the simplicial complexes for EF-Tu.** Recall that we use the atomic positions for each protein to create mesh representations of their 3D structures (see Fig. 1 in the main text). First, we draw an edge between any two atoms if their Euclidean distance smaller than some value $r$, namely $\text{dist}|(x_1, y_1, z_1), (x_2, y_2, z_2)| < r$. Next, we fill in all the triangles (or faces) formed by these connected edges. We treat the resulting triangulated mesh as an simplicial complex with which we can perform topological data analysis. Here, we consider the construction of mesh representations for each protein while setting $r = \{2.0, 4.0, 6.0\}$ ångströms (Å). Other SINATRA parameters were fixed: $c = 20$ cones, $d = 8$ directions per cone, $\theta = 0.80$ cap radius used to generate directions in a cone, and $l = 120$ sublevel sets per filtration. In each plot, we show the association metrics (and their corresponding standard errors) computed for each residue while analyzing **(a)** whole protein, **(b)** fragment 208-308, and **(c)** fragment 311-405. Note that an overlap in signal shows the robustness of SINATRA Pro to this input parameter value.

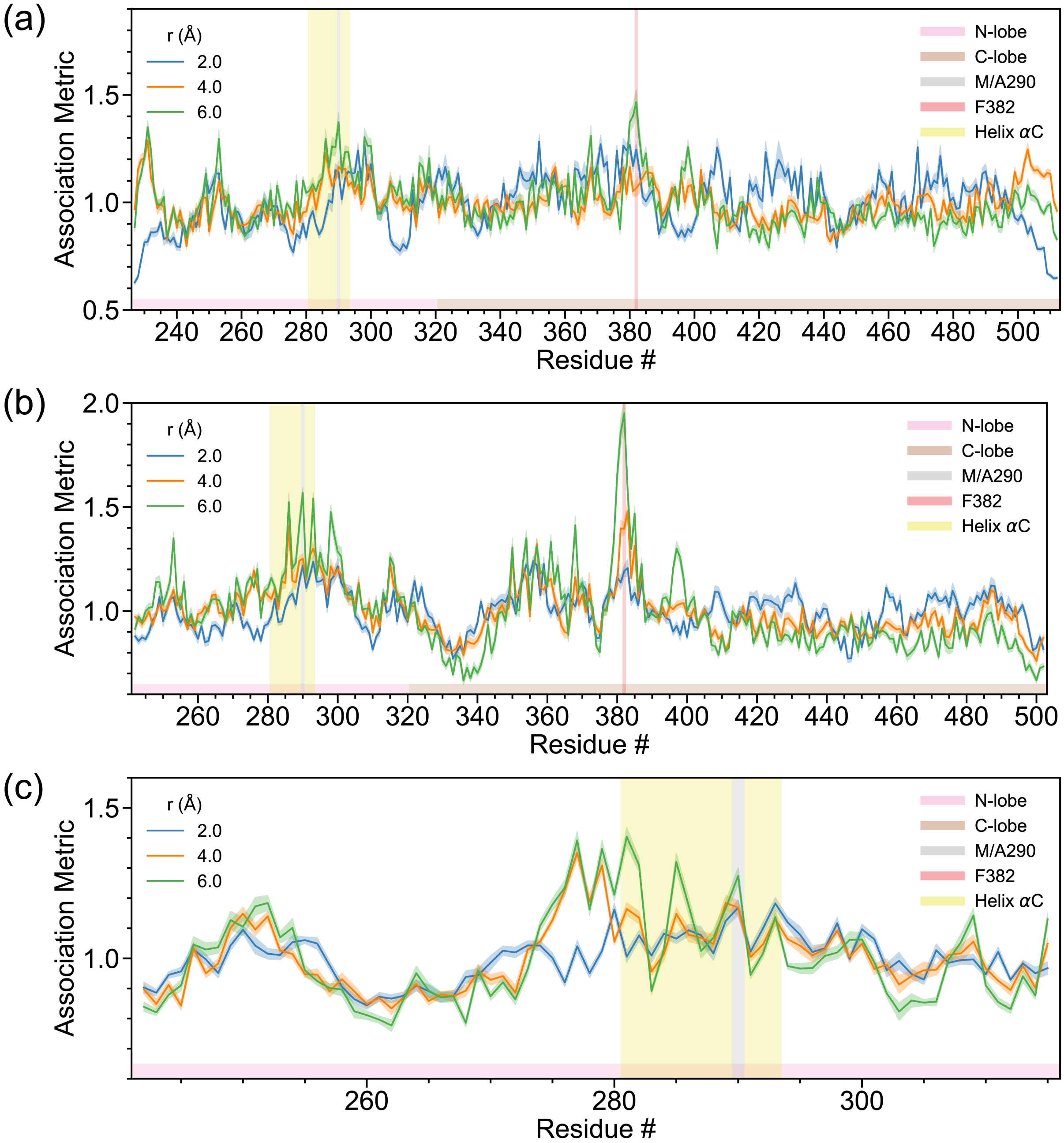

**Figure S25. Sensitivity analysis assessing the robustness of SINATRA Pro to different radius cutoffs $r$ values used to construct the simplicial complexes for Abl1.** Recall that we use the atomic positions for each protein to create mesh representations of their 3D structures (see Fig. 1 in the main text). First, we draw an edge between any two atoms if their Euclidean distance smaller than some value $r$, namely $\text{dist}|(x_1, y_1, z_1), (x_2, y_2, z_2)| < r$. Next, we fill in all the triangles (or faces) formed by these connected edges. We treat the resulting triangulated mesh as an simplicial complex with which we can perform topological data analysis. Here, we consider the construction of mesh representations for each protein while setting $r = \{2.0, 4.0, 6.0\}$ ångströms (Å). Other SINATRA parameters were fixed: $c = 20$ cones, $d = 8$ directions per cone, $\theta = 0.80$ cap radius used to generate directions in a cone, and $l = 120$ sublevel sets per filtration. In each plot, we show the association metrics (and their corresponding standard errors) computed for each residue while analyzing **(a)** the whole protein, **(b)** fragment 242-502, and **(c)** fragment 242-315. Note that an overlap in signal shows the robustness of SINATRA Pro to this input parameter value.

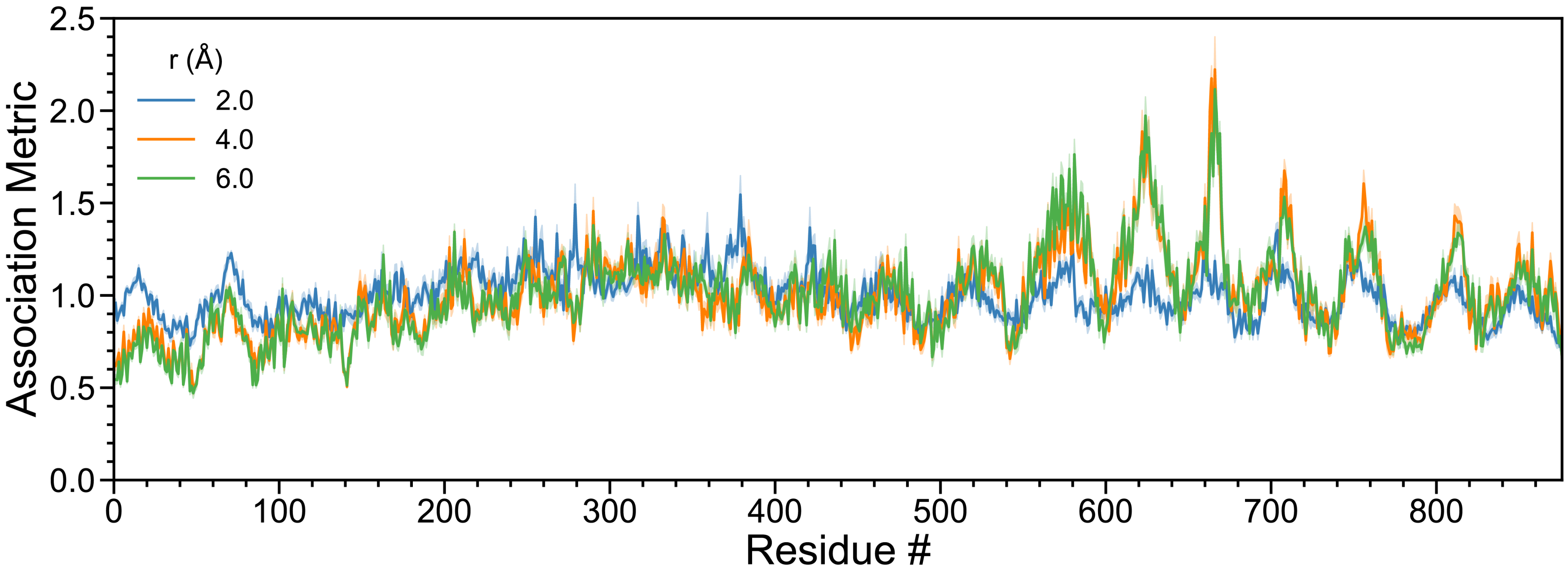

**Figure S26. Sensitivity analysis assessing the robustness of SINATRA Pro to different radius cutoffs $r$ values used to construct the simplicial complexes for Importin-$\beta$.** Recall that we use the atomic positions for each protein to create mesh representations of their 3D structures (see Fig. 1 in the main text). First, we draw an edge between any two atoms if their Euclidean distance smaller than some value $r$, namely dist$|(x_1, y_1, z_1), (x_2, y_2, z_2)| < r$. Next, we fill in all the triangles (or faces) formed by these connected edges. We treat the resulting triangulated mesh as an simplicial complex with which we can perform topological data analysis. Here, we consider the construction of mesh representations for each protein while setting $r = \{2.0, 4.0, 6.0\}$ ångströms (Å). Other SINATRA parameters were fixed: $c = 20$ cones, $d = 8$ directions per cone, $\theta = 0.80$ cap radius used to generate directions in a cone, and $l = 120$ sublevel sets per filtration. Note that an overlap in signal shows the robustness of SINATRA Pro to this input parameter value.

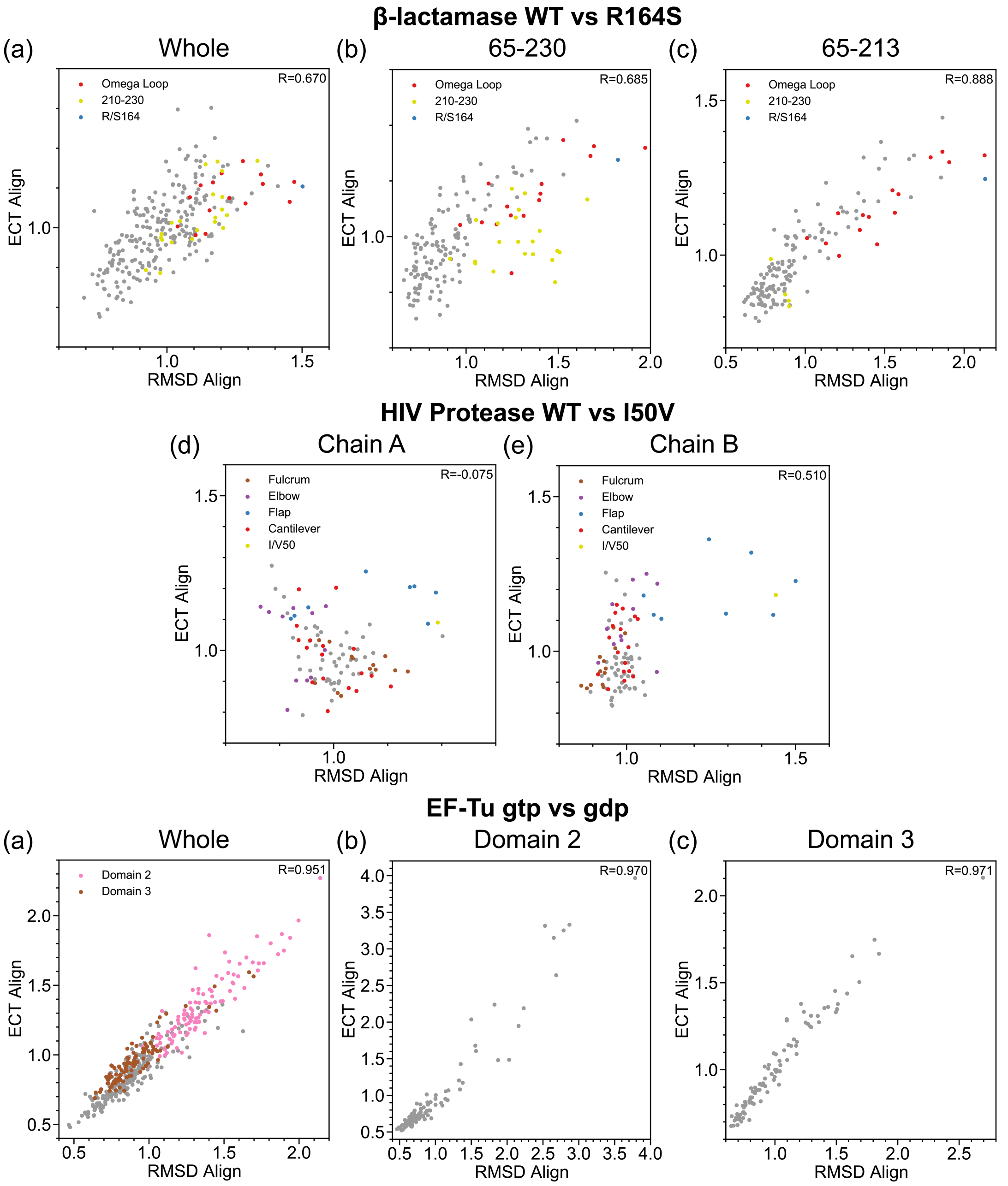

**Figure S27. Real data analysis results demonstrating the idea of running SINATRA Pro with sequence-independent (or correspondence free) structural alignment based on topological summary statistics.** Here, we perform sequence-independent structural alignment where we *implicitly* normalize the 3D protein structures by rotationally aligning their topological summary statistics. To carry out this alignment procedure, we first take each pair of protein structures and superimpose the center of mass of the backbone alpha-carbons ($C_\alpha$) atoms to the same origin. Next, we compute topological summary statistics over the mesh representation of each structure in $m = 500$ spherically uniformly distributed directions (see the Material and Methods in the main text). We take the squared Euclidean distance between any two directions to be the cost needed to align structures via their topological summaries; and we determine the "optimal" direction alignment by finding the rotation that minimizes the cumulative cost of aligning all directional pairs between proteins. We use the random sample consensus (RANSAC) method to determine the rotational matrix that aligns the angle between any two directions to be within an error threshold of 0.9 [22]. More specifically, we require that the dot product between two directions has to be larger than 0.9 to be considered aligned in RANSAC. The figures above compare the correlation ($R$) between the association metrics from SINATRA Pro calculated from the structures aligned using Euler characteristic (EC) transform and structures pre-aligned using RMSD for $\beta$-lactamase: **(a)** the whole protein, **(b)** fragment 65-230, and **(c)** fragment 65-213; monomers of HIV-1 protease: **(d)** chain A and **(e)** chain B; and fragments of the elongation factor EF-Tu: **(f)** the whole protein, **(g)** fragment 220-310 (Domain 2), and **(h)** fragment 311-405 (Domain 3). Correlations near one symbolize high agreement between the two alignment schemes.

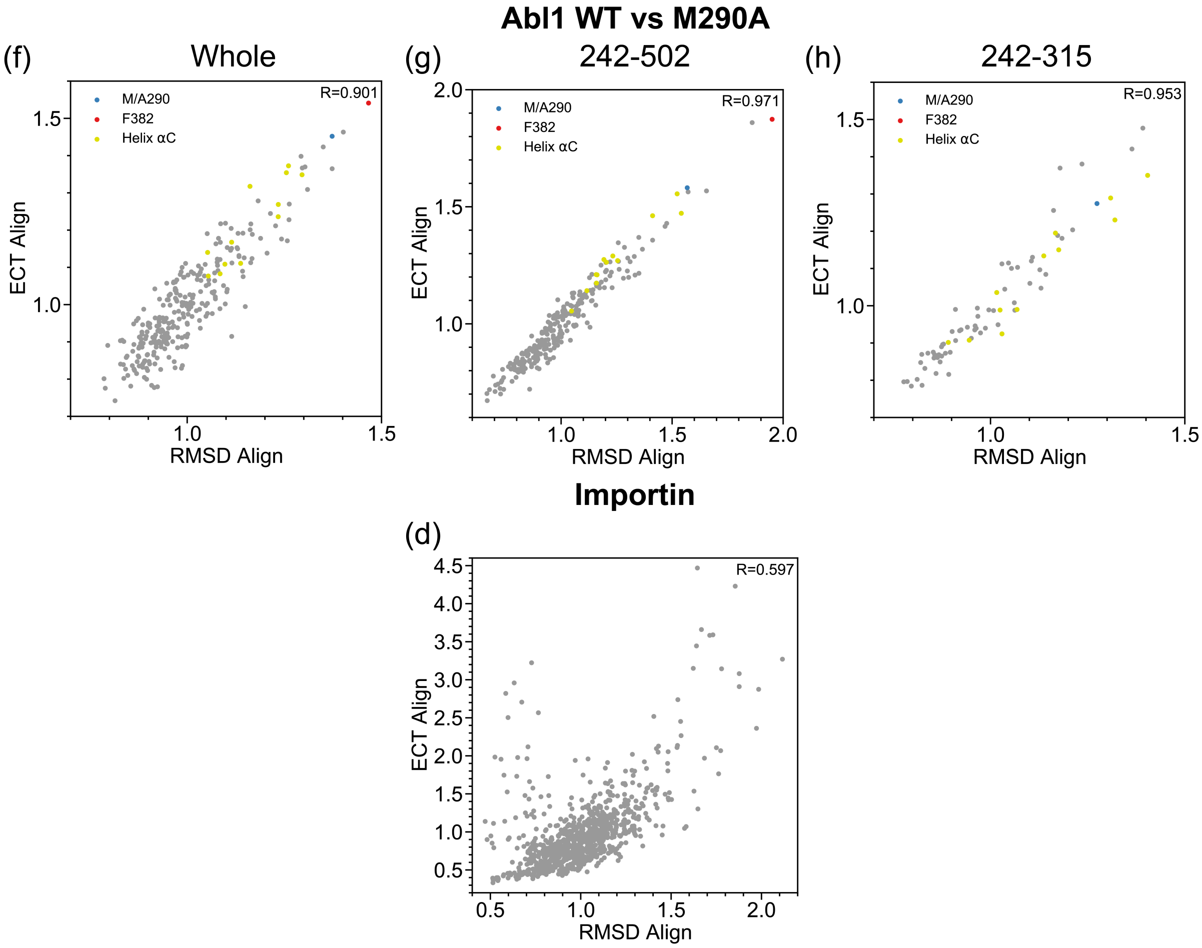

**Figure S28. Real data analysis results demonstrating the idea of running SINATRA Pro with sequence-independent (or correspondence free) structural alignment based on topological summary statistics.** Here, we perform sequence-independent structural alignment where we *implicitly* normalize the 3D protein structures by rotationally aligning their topological summary statistics. To carry out this alignment procedure, we first take each pair of protein structures and superimpose the center of mass of the backbone alpha-carbons ($C_\alpha$) atoms to the same origin. Next, we compute topological summary statistics over the mesh representation of each structure in $m = 500$ spherically uniformly distributed directions (see the Material and Methods in the main text). We take the squared Euclidean distance between any two directions to be the cost needed to align structures via their topological summaries; and we determine the "optimal" direction alignment by finding the rotation that minimizes the cumulative cost of aligning all directional pairs between proteins. We use the random sample consensus (RANSAC) method to determine the rotational matrix that aligns the angle between any two directions to be within an error threshold of 0.9 [22]. More specifically, we require that the dot product between two directions has to be larger than 0.9 to be considered aligned in RANSAC. The figures above compare the correlation ($R$) between the association metrics from SINATRA Pro calculated from the structures aligned using ECT and structures pre-aligned using RMSD for fragments of Abl1 Tyrosine protein kinase: **(a)** the whole protein, **(b)** fragment 242-502, and **(c)** fragment 242-315; and **(d)** unbound Importin-$\beta$. Correlations near one symbolize high agreement between the two alignment schemes.

#### 2 Supplementary Tables

| Total Proteins $N = 50$ | Number of Cones $c = 15$ | | | |
|---|---|---|---|---|
| | Directions per Cone $d = 4$ | | Directions per Cone $d = 8$ | |
| | Sublevel Sets $l = 25$ | Sublevel Sets $l = 50$ | Sublevel Sets $l = 25$ | Sublevel Sets $l = 50$ |
| (1) Read in PDB Structures | $45.0 \pm 0.6$ | $45.0 \pm 0.5$ | $44.6 \pm 0.0$ | $45.0 \pm 0.5$ |
| (2) Construct Meshes/Simplicial Complexes | $79.0 \pm 1.2$ | $78.1 \pm 0.2$ | $78.1 \pm 0.6$ | $78.3 \pm 0.4$ |
| (3) Compute Diff. Euler Characteristics | $74.1 \pm 0.4$ | $74.5 \pm 0.2$ | $86.0 \pm 0.7$ | $86.6 \pm 0.4$ |
| (4) Compute Atomic Variable Importance | $85.0 \pm 11.5$ | $128.3 \pm 3.1$ | $126.1 \pm 4.5$ | $461.5 \pm 10.8$ |
| (5) Reconstruct PDB Structures/Enrichments | $22.7 \pm 0.4$ | $22.7 \pm 0.1$ | $23.0 \pm 0.2$ | $23.4 \pm 0.4$ |
| **Total Runtime:** | **$305.9 \pm 11.6$** | **$348.6 \pm 3.1$** | **$357.8 \pm 4.6$** | **$694.7 \pm 0.9$** |

| Total Proteins $N = 50$ | Number of Cones $c = 20$ | | | |
|---|---|---|---|---|
| | Directions per Cone $d = 4$ | | Directions per Cone $d = 8$ | |
| | Sublevel Sets $l = 25$ | Sublevel Sets $l = 50$ | Sublevel Sets $l = 25$ | Sublevel Sets $l = 50$ |
| (1) Read in PDB Structures | $45.1 \pm 0.6$ | $45.5 \pm 0.7$ | $45.2 \pm 0.7$ | $44.9 \pm 0.5$ |
| (2) Construct Meshes/Simplicial Complexes | $85.1 \pm 13.1$ | $78.7 \pm 0.8$ | $81.4 \pm 5.8$ | $77.8 \pm 0.5$ |
| (3) Compute Diff. Euler Characteristics | $79.4 \pm 0.3$ | $80.3 \pm 1.3$ | $93.2 \pm 0.5$ | $93.7 \pm 0.4$ |
| (4) Compute Atomic Variable Importance | $82.3 \pm 0.3$ | $192.3 \pm 3.2$ | $195.6 \pm 3.4$ | $991.2 \pm 7.4$ |
| (5) Reconstruct PDB Structures/Enrichments | $22.8 \pm 0.2$ | $23.1 \pm 0.5$ | $23.4 \pm 0.4$ | $23.3 \pm 0.3$ |
| **Total Runtime:** | **$314.8 \pm 13.2$** | **$420.0 \pm 3.7$** | **$438.7 \pm 6.8$** | **$1230.9 \pm 7.5$** |

**Table S1. Empirical runtimes for running the SINATRA algorithm as a function of its free parameters and inputs.** Each entry represents the time (in seconds) it takes to run each step of the SINATRA Pro algorithm based on: *(i)* the total number of proteins analyzed $N = 50$, *(ii)* the number of cones of directions $c = \{15, 20\}$, *(iii)* the number of directions within each cone $d = \{4, 8\}$, and *(iv)* the number of sublevel sets (i.e., filtration steps) used to compute the Euler characteristic (EC) along a given direction $l = \{25, 50\}$. We simulate 10 different datasets for each combination of parameter values. Values appearing after the $\pm$ symbol are the standard deviations of these estimated times across the different runs. Each analysis was performed using simulated protein structures with $\sim$ 2700 atoms and all runtimes were computed using a central processing unit (CPU) with 8 cores and 128 gigabytes (GB) of RAM.

| Total Proteins $N = 100$ | Number of Cones $c = 15$ | | | |
|---|---|---|---|---|
| | Directions per Cone $d = 4$ | | Directions per Cone $d = 8$ | |
| | Sublevel Sets $l = 25$ | Sublevel Sets $l = 50$ | Sublevel Sets $l = 25$ | Sublevel Sets $l = 50$ |
| (1) Read in PDB Structures | $52.2 \pm 0.3$ | $50.6 \pm 0.5$ | $50.4 \pm 0.1$ | $50.4 \pm 0.1$ |
| (2) Construct Meshes/Simplicial Complexes | $159.7 \pm 2.1$ | $158.7 \pm 0.6$ | $157.8 \pm 1.1$ | $157.6 \pm 0.8$ |
| (3) Compute Diff. Euler Characteristics | $151.8 \pm 0.3$ | $152.2 \pm 0.7$ | $178.1 \pm 0.9$ | $178.4 \pm 0.6$ |
| (4) Compute Atomic Variable Importance | $89.5 \pm 0.7$ | $141.2 \pm 1.5$ | $141.8 \pm 4.8$ | $489.5 \pm 3.4$ |
| (5) Reconstruct PDB Structures/Enrichments | $46.3 \pm 0.3$ | $46.5 \pm 0.2$ | $46.8 \pm 0.1$ | $47.2 \pm 0.3$ |
| **Total Runtime:** | $\mathbf{499.5 \pm 2.3}$ | $\mathbf{549.2 \pm 1.8}$ | $\mathbf{574.9 \pm 5.0}$ | $\mathbf{923.1 \pm 3.6}$ |

| Total Proteins $N = 100$ | Number of Cones $c = 20$ | | | |
|---|---|---|---|---|
| | Directions per Cone $d = 4$ | | Directions per Cone $d = 8$ | |
| | Sublevel Sets $l = 25$ | Sublevel Sets $l = 50$ | Sublevel Sets $l = 25$ | Sublevel Sets $l = 50$ |
| (1) Read in PDB Structures | $50.4 \pm 0.1$ | $50.3 \pm 0.1$ | $50.3 \pm 0.1$ | $50.4 \pm 0.2$ |
| (2) Construct Meshes/Simplicial Complexes | $158.1 \pm 0.4$ | $157.8 \pm 1.1$ | $157.9 \pm 0.5$ | $157.0 \pm 0.9$ |
| (3) Compute Diff. Euler Characteristics | $162.7 \pm 0.6$ | $162.5 \pm 0.4$ | $193.4 \pm 0.8$ | $193.6 \pm 0.6$ |
| (4) Compute Atomic Variable Importance | $98.1 \pm 0.5$ | $211.3 \pm 2.1$ | $209.4 \pm 1.7$ | $1022.8 \pm 17.4$ |
| (5) Reconstruct PDB Structures/Enrichments | $46.8 \pm 0.2$ | $46.7 \pm 0.3$ | $47.4 \pm 0.2$ | $47.6 \pm 0.8$ |
| **Total Runtime:** | $\mathbf{516.1 \pm 0.9}$ | $\mathbf{628.6 \pm 2.4}$ | $\mathbf{658.4 \pm 1.9}$ | $\mathbf{1471.4 \pm 17.5}$ |

**Table S2. Empirical runtimes for running the SINATRA algorithm as a function of its free parameters and inputs.** Each entry represents the time (in seconds) it takes to run each step of the SINATRA Pro algorithm based on: *(i)* the total number of proteins analyzed $N = 100$, *(ii)* the number of cones of directions $c = \{15, 20\}$, *(iii)* the number of directions within each cone $d = \{4, 8\}$, and *(iv)* the number of sublevel sets (i.e., filtration steps) used to compute the Euler characteristic (EC) along a given direction $l = \{25, 50\}$. We simulate 10 different datasets for each combination of parameter values. Values appearing after the $\pm$ symbol are the standard deviations of these estimated times across the different runs. Each analysis was performed using simulated protein structures with $\sim 2700$ atoms and all runtimes were computed using a central processing unit (CPU) with 8 cores and 128 gigabytes (GB) of RAM.

| Total Proteins $N = 200$ | Number of Cones $c = 15$ | | | |
|---|---|---|---|---|
| | Directions per Cone $d = 4$ | | Directions per Cone $d = 8$ | |
| | Sublevel Sets $l = 25$ | Sublevel Sets $l = 50$ | Sublevel Sets $l = 25$ | Sublevel Sets $l = 50$ |
| (1) Read in PDB Structures | $63.5 \pm 1.3$ | $63.3 \pm 1.6$ | $64.3 \pm 2.8$ | $63.1 \pm 1.1$ |
| (2) Construct Meshes/Simplicial Complexes | $305.3 \pm 1.4$ | $304.6 \pm 2.2$ | $305.4 \pm 0.9$ | $304.2 \pm 1.1$ |
| (3) Compute Diff. Euler Characteristics | $296.1 \pm 0.8$ | $297.1 \pm 0.4$ | $347.8 \pm 1.7$ | $348.0 \pm 1.2$ |
| (4) Compute Atomic Variable Importance | $111.9 \pm 0.8$ | $158.8 \pm 0.7$ | $161.8 \pm 4.9$ | $493.3 \pm 1.1$ |
| (5) Reconstruct PDB Structures/Enrichments | $89.1 \pm 0.5$ | $89.8 \pm 1.2$ | $90.7 \pm 0.5$ | $90.9 \pm 0.3$ |
| **Total Runtime:** | $\mathbf{865.9 \pm 2.2}$ | $\mathbf{913.5 \pm 3.1}$ | $\mathbf{970.1 \pm 6.0}$ | $\mathbf{1299.5 \pm 2.3}$ |

| Total Proteins $N = 200$ | Number of Cones $c = 20$ | | | |
|---|---|---|---|---|
| | Directions per Cone $d = 4$ | | Directions per Cone $d = 8$ | |
| | Sublevel Sets $l = 25$ | Sublevel Sets $l = 50$ | Sublevel Sets $l = 25$ | Sublevel Sets $l = 50$ |
| (1) Read in PDB Structures | $63.7 \pm 1.2$ | $63.5 \pm 1.5$ | $63.9 \pm 1.1$ | $64.0 \pm 1.1$ |
| (2) Construct Meshes/Simplicial Complexes | $310.5 \pm 11.8$ | $304.5 \pm 0.6$ | $304.8 \pm 0.9$ | $304.7 \pm 1.1$ |
| (3) Compute Diff. Euler Characteristics | $320.0 \pm 0.5$ | $319.6 \pm 0.4$ | $377.0 \pm 1.4$ | $377.6 \pm 1.5$ |
| (4) Compute Atomic Variable Importance | $119.9 \pm 0.5$ | $226.4 \pm 0.5$ | $227.5 \pm 2.3$ | $1001.7 \pm 6.9$ |
| (5) Reconstruct PDB Structures/Enrichments | $91.2 \pm 0.7$ | $90.6 \pm 0.7$ | $92.8 \pm 1.1$ | $92.6 \pm 1.2$ |
| **Total Runtime:** | $\mathbf{905.3 \pm 11.9}$ | $\mathbf{1004.6 \pm 1.9}$ | $\mathbf{1066.1 \pm 3.3}$ | $\mathbf{1840.6 \pm 7.3}$ |

**Table S3. Empirical runtimes for running the SINATRA algorithm as a function of its free parameters and inputs.** Each entry represents the time (in seconds) it takes to run each step of the SINATRA Pro algorithm based on: *(i)* the total number of proteins analyzed $N = 200$, *(ii)* the number of cones of directions $c = \{15, 20\}$, *(iii)* the number of directions within each cone $d = \{4, 8\}$, and *(iv)* the number of sublevel sets (i.e., filtration steps) used to compute the Euler characteristic (EC) along a given direction $l = \{25, 50\}$. We simulate 10 different datasets for each combination of parameter values. Values appearing after the $\pm$ symbol are the standard deviations of these estimated times across the different runs. Each analysis was performed using simulated protein structures with $\sim 2700$ atoms and all runtimes were computed using a central processing unit (CPU) with 8 cores and 128 gigabytes (GB) of RAM.

### References

1. Xu B, Wang N, Chen T, Li M. Empirical evaluation of rectified activations in convolutional network; 2015. ArXiv.

2. Tibshirani R. Regression shrinkage and selection via the lasso. J R Stat Soc Series B Stat Methodol. 1996;58(1):267–288.

3. Hoerl AE, Kennard RW. Ridge regression: Biased estimation for nonorthogonal problems. Technometrics. 1970;12(1):55–67.

4. Zou H, Hastie T. Regularization and variable selection via the elastic net. J R Stat Soc Series B Stat Methodol. 2005;67(2):301–320.

5. Simonyan K, Vedaldi A, Zisserman A. Deep inside convolutional networks: Visualising image classification models and saliency maps. arXiv preprint arXiv:13126034. 2013;.

6. Stojanoski V, Chow DC, Hu L, Sankaran B, Gilbert HF, Prasad BVV, et al. A triple mutant in the Ω-loop of TEM-1 $\beta$-lactamase changes the substrate profile via a large conformational change and an altered general base for catalysis. Journal of Biological Chemistry. 2015;290(16):10382–10394. Available from: `https://pubmed.ncbi.nlm.nih.gov/25713062`.

7. Egorov A, Rubtsova M, Grigorenko V, Uporov I, Veselovsky A. The Role of the Ω-Loop in Regulation of the Catalytic Activity of TEM-Type $\beta$-Lactamases. Biomolecules. 2019;9(12). Available from: `https://www.ncbi.nlm.nih.gov/pmc/articles/PMC6995641/`.

19. Cingolani G, Petosa C, Weis K, Müller CW. Structure of importin-$\beta$ bound to the IBB domain of importin-$\alpha$. Nature. 1999;399(6733):221–229. Number: 6733 Publisher: Nature Publishing Group. Available from: https://www.nature.com/articles/20367.

20. Zachariae U, Grubmüller H. Importin-$\beta$: Structural and Dynamic Determinants of a Molecular Spring. Structure. 2008;16(6):906–915. Available from: https://www.sciencedirect.com/science/article/pii/S0969212608001445.

21. Halder K, Dölker N, Van Q, Gregor I, Dickmanns A, Baade I, et al. MD Simulations and FRET Reveal an Environment-Sensitive Conformational Plasticity of Importin-$\beta$. Biophysical Journal. 2015;109(2):277–286. Available from: https://www.ncbi.nlm.nih.gov/pmc/articles/PMC4621615/.

22. Fischler MA, Bolles RC. Random Sample Consensus: A Paradigm for Model Fitting with Applications to Image Analysis and Automated Cartography. Commun ACM. 1981;24(6):381–395. Available from: https://doi.org/10.1145/358669.358692.
